# Synonymization of the male-based ant genus *Phaulomyrma* (Hymenoptera, Formicidae) with *Leptanilla* based upon Bayesian total-evidence phylogenetic inference

**DOI:** 10.1101/2020.08.28.272799

**Authors:** Zachary H. Griebenow

## Abstract

Although molecular data have proven indispensable in confidently resolving the phylogeny of many clades across the tree of life, these data may be inaccessible for certain taxa. The resolution of taxonomy in the ant subfamily Leptanillinae is made problematic by the absence of DNA sequence data for leptanilline taxa that are known only from male specimens, including the monotypic genus *Phaulomyrma* Wheeler & Wheeler. Focusing upon the considerable diversity of undescribed male leptanilline morphospecies, the phylogeny of 35 putative morphospecies sampled from across the Leptanillinae, plus an outgroup, is inferred from 11 nuclear loci and 41 discrete male morphological characters using a Bayesian total-evidence framework, with *Phaulomyrma* represented by morphological data only. Based upon the results of this analysis *Phaulomyrma* is synonymized with *Leptanilla* Emery, and male-based diagnoses for *Leptanilla* that are grounded in phylogeny are provided, under both broad and narrow circumscriptions of that genus. This demonstrates the potential utility of a total-evidence approach in inferring the phylogeny of rare extant taxa for which molecular data are unavailable and begins a long-overdue systematic revision of the Leptanillinae that is focused on male material.

## Introduction

Over the past three decades, DNA sequences have provided great insight into phylogenetic relationships across the Metazoa, including the insects (Kjer *et al*. 2018). The application of maximum-likelihood (ML) and Bayesian statistical methods to analysis of genetic data has robustly resolved many problems that were intractable when using morphological data alone (e.g. Niehuis *et al*. 2012; Wipfler *et al*. 2019). However, DNA sequences may be unavailable for some taxa, necessitating the integration of morphological and molecular data under the same inferential framework. Fossils are the most obvious example of this: these are valuable for calibration of phylogenies in absolute time under a Bayesian approach, preferably with their topological position being inferred from the data (Ronquist *et al*. 2012; O’Reilly *et al*. 2015; Bapst et al. 2016; Matzke & Wright 2016). Although the inclusion of fossils for the purposes of “tip-dating” has received the bulk of attention in Bayesian total-evidence phylogenetic inference, the lack of molecular data may afflict rare extant taxa as well (Sánchez *et al*. 2016; Robertson & Moore 2016). This is problematic if the affinities of these taxa are not immediately clear from morphology alone.

The ant subfamily Leptanillinae (Hymenoptera: Formicidae) is an apt test case for methods to resolve this problem. A group of small, hypogaeic ants largely restricted to the Old World tropics and subtropics, the Leptanillinae are understood to be one of the earliest-diverging lineages in the ant crown-group (Rabeling *et al*. 2008; Kück *et al*. 2011; Borowiec *et al*. 2019; Boudinot *et al*. submitted). Three out of eight described genera are known from both workers and males: *Opamyrma* Yamane, Bui & Eguchi, 2008 (Yamada *et al*. 2020), *Protanilla* Taylor, 1990 (Griebenow, in press), and *Leptanilla* Emery, 1870 (e.g. Ogata *et al*. 1995). Males of *Anomalomyrma* Taylor, 1990 are unknown. Four leptanilline genera—*Scyphodon* Brues, 1925; *Phaulomyrma* Wheeler & Wheeler, 1930; *Noonilla* Petersen, 1968; and *Yavnella* Kugler, 1986*—* have been described solely from males, as have many species of *Leptanilla* (cf. Bolton 1990). Recent molecular data indicate that the type species of *Yavnella* and a specimen provisionally assigned to *Phaulomyrma* are nested within a clade of putative *Leptanilla* morphospecies (Borowiec *et al*. 2019). Moreover, although *Scyphodon anomalum* Brues, 1925 and *Noonilla copiosa* Petersen, 1968 exhibit bizarre autapomorphies such as hypertrophied mandibles (Brues 1925) and a ventromedian genital “trigger” (Petersen 1968), respectively, these ants are otherwise similar to males attributed to *Leptanilla* (Boudinot 2015).

This indicates a need for a systematic revision of the Leptanillinae, but almost all published taxonomic studies of the group have been descriptive without recourse to molecular phylogeny, with the exceptions being revisions to our concept of the subfamily. Multi-locus DNA datasets demonstrated that the enigmatic Afrotropical genus *Apomyrma* Gotwald, Brown & Lévieux, 1971 is closely related to the Amblyoponinae rather than the Leptanillinae (Brady *et al*. 2006; Moreau *et al*. 2006), and that the superficially similar Asian genus *Opamyrma* is in fact sister to the remaining Leptanillinae (Ward & Fisher 2016). None of these studies focused upon the Leptanillinae or the internal phylogeny of this clade. Such a study must confront two challenges: first, the lack of DNA sequences for certain critical taxa across the Leptanillinae (e.g., *Scyphodon*), which hampers any attempt to confidently resolve relationships among these; second, the definition of genera based only upon males, which prevents an integrated phylogenetic classification of the Leptanillinae, since phenotypes of only one sex are considered.

The dissociation of leptanilline castes results from collecting bias. Subterranean workers have been largely collected with *lavage de terre* methodology (López *et al*. 1994; Wong and Guénard 2016), Winkler trapping (Belshaw & Bolton 1994; Leong *et al*. 2018), and subterranean pitfall traps (Wong & Guénard 2016; Man *et al*. 2017); whereas male leptanillines are typically collected by sweeping foliage or by deploying Malaise or pan traps (Robertson 2000). None of these methods are likely to collect males in association with workers, nor is the queen caste often collected in association with conspecifics. Contrasting with the alate condition observed in most ants, queens described from the tribe Leptanillini are completely wingless and blind (Emery 1870; Kutter 1948; Masuko 1990; López *et al*. 1994; Ogata *et al*. 1995), meaning that these are no more likely to be collected than corresponding workers. Queens belonging to other leptanilline lineages (*Opamyrma* and the Anomalomyrmini) are alate so far as is known (Bolton 1990; Baroni Urbani & de Andrade 2008; Borowiec *et al*. 2011; Chen *et al*. 2017; Hsu *et al*. 2017; Man *et al*. 2017), save for an apparent record of queen polyphenism in an undescribed *Protanilla* (Billen *et al*. 2013), but are infrequently collected.

Therefore, the bulk of known leptanilline diversity, most of it undescribed, is represented by exclusively male material. In some cases, molecular data are inaccessible for male morphotaxa due to paucity of suitably recent specimens, obliging a total-evidence approach to infer the phylogeny of these lineages. This study uses such an approach to resolve the position of the male-based species *Phaulomyrma javana* Wheeler & Wheeler, 1930, the sole species included in this genus. Here, the phylogeny of the Leptanillinae is inferred jointly from 10 protein-coding genes, 28S rDNA, and 41 discrete male morphological characters under a Bayesian statistical framework. This is the first combined-evidence Bayesian analysis to include the Leptanillinae and is novel among studies of ant phylogeny in its inclusion of exclusively male morphological characters (Barden *et al*. [2017] used both worker and male morphology in their Bayesian total-evidence inference). Despite the absence of nucleotide sequences for *P. javana* a Bayesian total-evidence approach facilitates the inclusion of this terminal and its confident phylogenetic placement. Based upon the results of these joint molecular and morphological phylogenetic analyses, a revised male-based definition of *Leptanilla* is provided, and *Phaulomyrma* is synonymized with that genus.

## Materials and Methods

### Taxon Sampling

Thirty-five terminals were included in total (Tables 1-2). Discrete morphological data were scored for those 33 terminals for which male material was known. *Anomalomyrma boltoni* Borowiec, Schulz, Alpert & Baňar, 2011 and *Leptanilla revelierii* Emery, 1870 were represented in this study by workers alone. The latter was included on account of its status as the type species of that genus: regardless of future systematic revision to the Leptanillinae, the concept of the genus *Leptanilla* will not exclude this species. DNA sequences for the outgroup *M. heureka* were obtained from a worker ant, as published in Borowiec *et al*. (2019). Most putative morphospecies were represented by singletons (Table 1), but phenotypic variation within those morphospecies for which material was abundant (e.g., *Leptanilla* zhg-my02) is minimal, and so gives no reason to suspect heterospecificity among the specimens referred to these morphospecies.

**Table 1.**
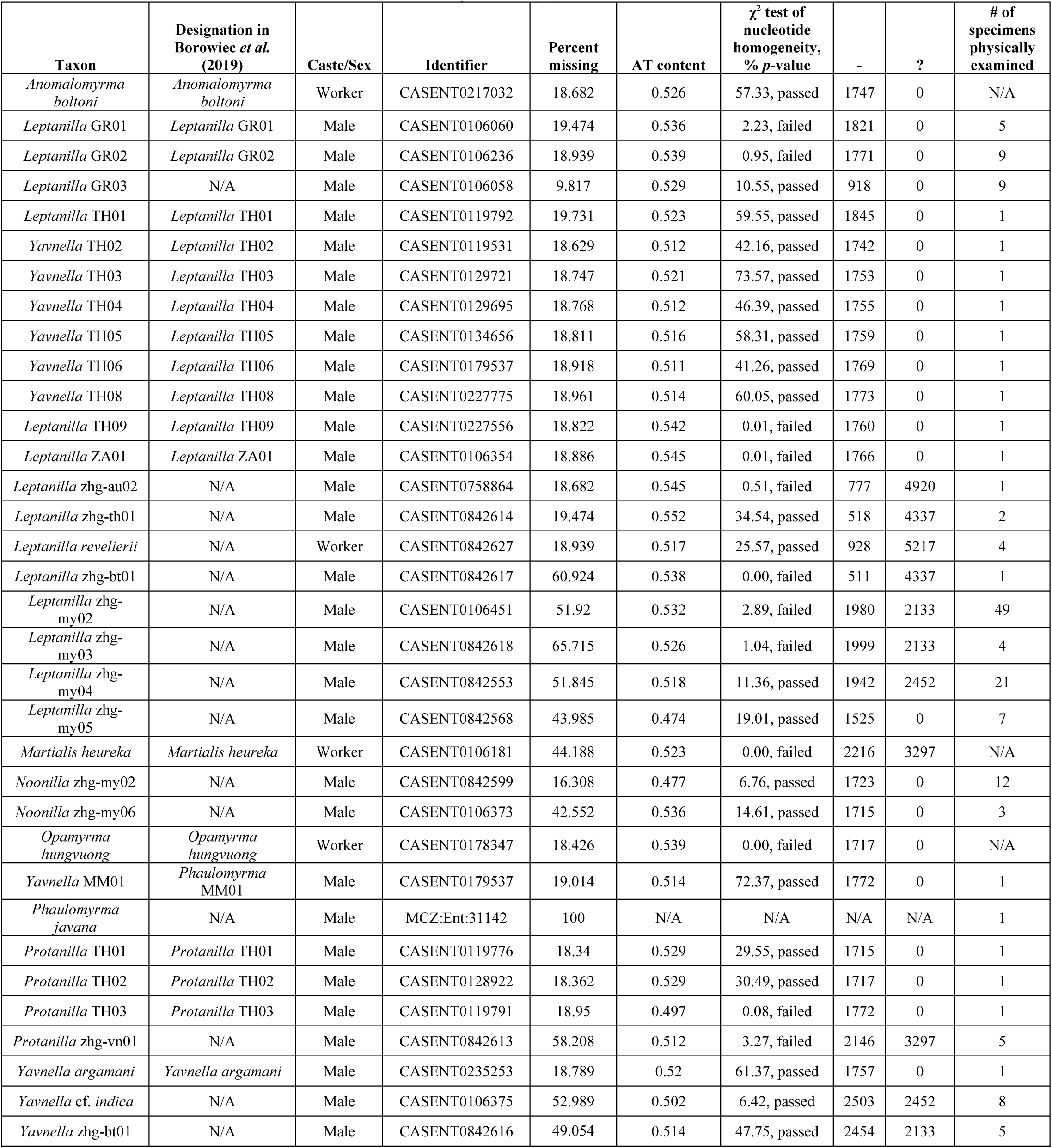
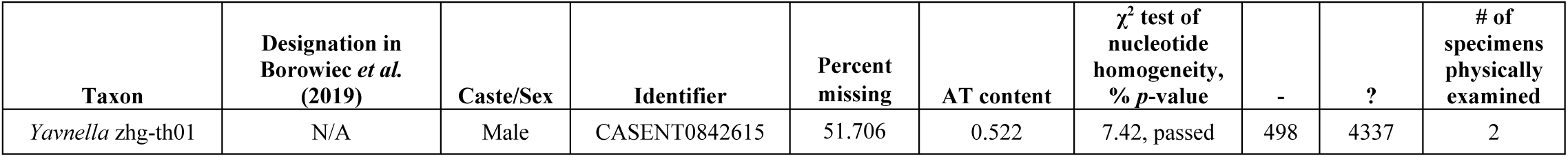
Summary statistics for full 9,351-bp DNA legacy-locus alignment. “–” = absent base; “” = unknown base. χ^2^ test of nucleotide homogeneity executed with IQ-Tree 1.6.10 (Nguyen *et al*. 2015) on the CIPRES Science Gateway (v. 3.3) (Miller *et al*. 2010).

**Table 2.**
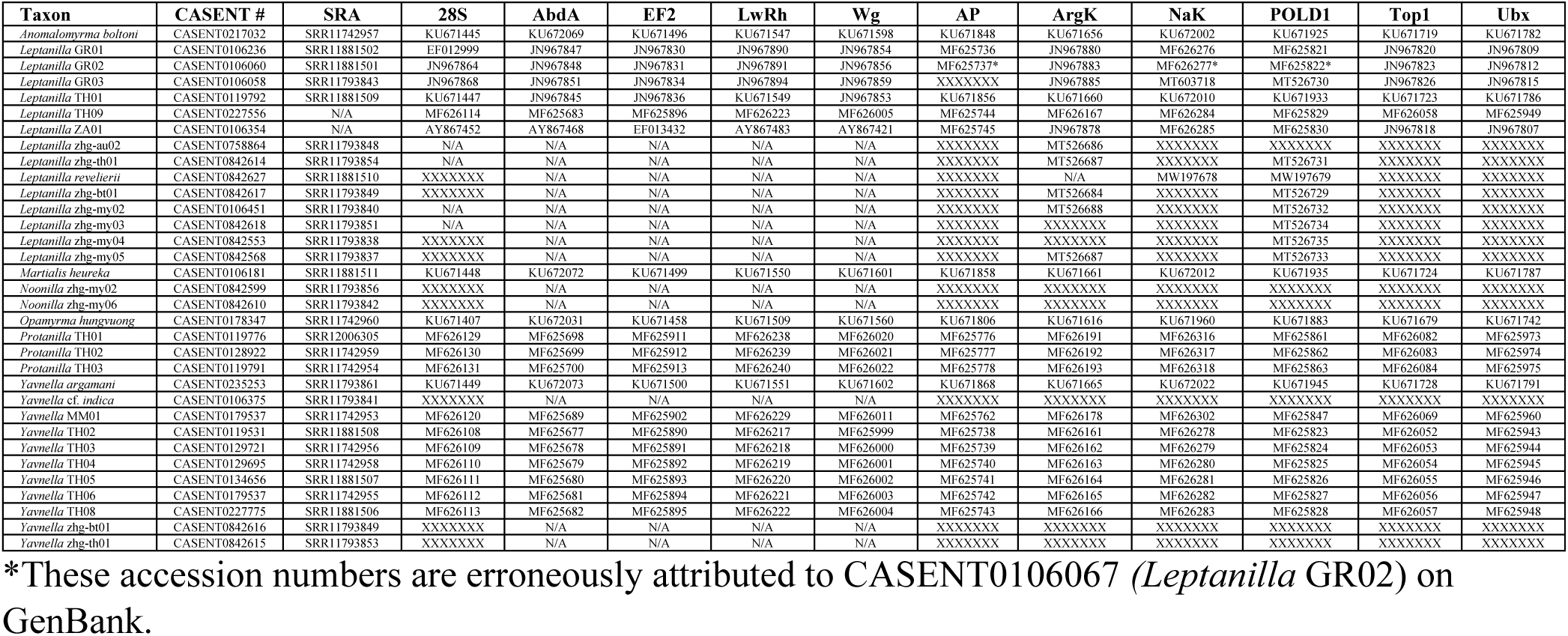
NCBI and SRA accession numbers for DNA sequences used in Bayesian total-evidence inference.

Representatives of all male-based genera were included in total-evidence analyses, except for *Scyphodon*. These include both *Yavnella argamani* Kugler, 1986 and *Yavnella* cf. *indica*, along with two undescribed *Yavnella* morphospecies from Bhutan and Thailand, respectively; *Phaulomyma javana*; and two morphospecies of *Noonilla* identified as such according to the definition given by Petersen (1968). *Leptanilla* TH02-6 and −08, along with *Phaulomyrma* MM01, were placed in those genera by Borowiec *et al*. (2019) and/or Boudinot (2015) but are here identified as *Yavnella* (Table 1) according to the definition of Kugler (1986).

Material is deposited in the following repositories: the Bohart Museum of Entomology, University of California, Davis, CA, USA (UCDC); the California Academy of Sciences, San Francisco, CA, USA (CASC); the California State Collection of Arthropods, Sacramento, CA, USA (CSCA); the Lund Museum of Zoology, Lund, Sweden (MZLU); and the Australian National Insect Collection, Canberra, Australia (ANIC).

### Molecular Dataset

Total-evidence phylogenetic inference was based upon 11 nuclear loci: *28S ribosomal DNA* (*28S*)*, abdominal-A* (*abdA*)*, arginine kinase* (*argK*)*, antennapedia* (*Antp*)*, elongation factor 1-alpha F2 copy* (*EF1*α*F2*)*, long wavelength rhodopsin* (*LW Rh*)*, NaK ATPase* (*NaK*)*, DNA pol-delta* (*POLD1*)*, topoisomerase I* (*Top1*)*, ultrabithorax* (*Ubx*), and *wingless* (*Wg*). I derived these “legacy loci” for 19 terminals from the alignment of Borowiec *et al*. (2019) (doi: 10.5281/zenodo.2549806) but expanded to include autapomorphic indels and introns, and constituting 11,090 bp. Legacy loci for *Leptanilla* GR03 were derived from Ward & Sumnicht (2012). For further detail on the protocols for the extraction and amplification of these genetic data, refer to Ward *et al*. (2010) and Ward & Fisher (2016). I added 14 terminals to this “legacy-locus” intron-inclusive dataset by retrieving orthologous loci from phylogenomic data acquired with the ultra-conserved element (UCE) probe set *hym*-*v2* (Branstetter *et al*. 2017). *P. javana* was the only terminal for which molecular data were not obtained: this species is known only from two slide-mounted syntypes collected in 1907.

DNA was extracted non-destructively using a DNeasy Blood and Tissue Kit (Qiagen Inc., Valencia, CA) according to manufacturer instructions. DNA was quantified for each sample with a Qubit 2.0 fluorometer (Life Technologies Inc., Carlsbad, CA). Phylogenomic data were obtained from these taxa using the *hym-v2* probe set, with libraries being prepared and target loci enriched using the protocol of Branstetter *et al*. (2017). Enrichment success and size-adjusted DNA concentrations of pools were assessed using the SYBR FAST qPCR kit (Kapa Biosystems, Wilmington, MA) and all pools were combined into an equimolar final pool. The contents of this final pool were sequenced by an Illumina HiSeq 2500 at the University of Utah’s High Throughput Genomics Facility or an Illumina HiSeq 4000 at Novogene, Sacramento, CA. The FASTQ output was demultiplexed and cleansed of adapter contamination and low-quality reads using *illumiprocessor* (Faircloth 2013) in the PHYLUCE package. Raw reads were then assembled with *trinity* v. 2013-02-25 (Grabherr *et al*. 2011) or SPAdes v. 3.12.0 (Bankevich *et al*. 2013). The possibility of genetic contamination and/or mis-assembly in the UCE samples was tested by inferring a phylogeny from a concatenated UCE alignment, unpartitioned, using IQ-Tree 1.6.10 (Nguyen *et al*. 2015) on the CIPRES Science Gateway (v. 3.3) (Miller *et al*. 2010) with the GTR+G model of substitution for 1,000 ultrafast bootstrap replicates (Hoang *et al*. 2018): this phylogeny was plausible given preliminary hypotheses, providing no positive evidence of sequence contamination or mis-assembly. Summary statistics for these UCE assemblies were computed using *statswrapper.sh* in *BBMap* (Bushnell 2014), and provided in Supplemental Table 1.

In the cases of the 14 terminals not included in Ward & Sumnicht (2012) or Borowiec *et al*. (2019) for which molecular data could be obtained, legacy loci orthologous with those used by Borowiec *et al*. (2019) were then recovered from genome-scale data as follows using PHYLUCE (Faircloth 2016). I derived sequences representing each locus for *Leptanilla* GR02 from the alignment ANT-exon-sequences-40-taxa-reduced.fasta published by Branstetter *et al*. (2017), given the comparative completeness of the matrix for that species, and its phylogenetic position nested well within the Leptanillinae. These sequences were then used analogously to probes.

Species-specific contig assemblies were obtained using *phyluce_assembly_match_contigs_to_probes.py* (min_coverage = 50, min_identity = 85), a list of legacy loci shared across all taxa was generated using *phyluce_assembly_get_match_counts.py*, and separate FASTA files for each locus were created using these outputs. Sequences were aligned separately by locus using MAFFT (Katoh *et al*. 2009) implemented with the command *phyluce_assembly_seqcap_align.py*, and these sequences were then trimmed with Gblocks (Castresana 2000) as implemented by the wrapper script *phyluce_assembly_get_gblocks_trimmed_alignment_from_untrimmed.py* (settings: b1 = 0.5, b2 = 0.5, b3 = 12, b4 = 7). Alignment statistics for the output FASTA files were calculated with *phyluce_align_get_align_summary_data.py*. Finally, a matrix that was 80% complete with respect to locus coverage was generated using the script *phyluce_align_get_only_loci_with_min_taxa.py*. This contained 7 out of the 10 protein-coding loci that I attempted to recover using the exon-based bioinformatic protocol of Branstetter *et al*. (2017), in addition to 28S rDNA. Legacy loci recovered from UCE assemblies often included non-coding sequences adjacent to the regions included in Borowiec *et al*. (2019), which were trimmed manually in AliView. In whichever cases those loci had been recovered, sequences for the taxa represented only in the dataset of Borowiec *et al*. (2019) were then aligned with the recovered legacy loci using the online MAFFT interface (Katoh *et al*. 2019) with default settings. In cases where legacy loci were not successfully recovered or were incomplete relative to preexisting Sanger-derived sequences, these loci were derived from the datasets of Borowiec *et al*. (2019) or, in the case of *Leptanilla* GR03, Ward & Sumnicht (2012). These data were concatenated with UCE-derived sequences across all FASTA files, inasmuch all sequences for each morphospecies were derived from the same specimen; and all loci were concatenated to produce a final alignment, which was 9,351 bp in length. Further summary statistics for this final alignment are provided in Table 1 and Supplemental Table 2. Alignment was unambiguous once all loci were brought into their respective reading frames. GenBank accession numbers for all loci used in this study are provided in Table 2.

Those terminals for which loci were obtained using the 11 nuclear loci from *Leptanilla* GR02 as “probes” according to the modified PHYLUCE protocol cited above (“Molecular Dataset”) (Faircloth 2016), with 80% locus coverage implemented in *phyluce_align_get_only_loci_with_min_taxa.py*, exhibit low coverage relative to those that were sequenced prior to this study (Table 1). Therefore, a 9,062-bp legacy-locus alignment was created that includes only those data published prior to this study (Ward & Sumnicht 2012; Borowiec *et al*. 2019), with 20 terminals. These sequences can be used to test the possibility that missing data would have an appreciable effect on phylogenetic inference phylogenetic analyses.

### Morphological Dataset

Forty-one discrete binary morphological characters were coded for all 33 morphospecies known from males. All these specimens were examined with a Leica MZ75 compound microscope or by reference to images on AntWeb, except for the male of *M. heureka* and *O. hungvuong*, in the cases of which observations were derived from Boudinot (2015: Figs. 11-12) and Yamada *et al*. (2020: Figs. 11-13), respectively, or from the textual descriptions by those authors. I imaged specimens when necessary using a JVC KY-F75 digital camera and compiled color photographs from these with the Syncroscopy AutoMontage Program. Scanning electron microscopy was undertaken using a Hitachi TM4000 tabletop microscope. Morphological terminology follows the Hymenoptera Anatomy Ontology (Yoder *et al*. 2010), with some exceptions being derived from Bolton (2003) and Boudinot (2018). The character coding scheme was binary and non-additive (Pleijel 1995). Missing data were scored as ‘?’. Autapomorphic characters were included. Numerical scores for all morphological characters are presented in the Supplemental Table 3.

Non-additive binary coding has been criticized for its susceptibility to redundancy (Strong & Lipscomb 1999), stipulation of compound characters, and the inadvertent conflation of morphological absences that are not hierarchically equivalent (Brazeau 2011). These problems largely result from careless character delimitation. I compensated for these potential flaws by defining and using only characters that do not logically depend upon other characters. Definitions of morphological character states are provided in the Appendix.

### Phylogenetic Analyses

For the two legacy-locus molecular datasets, the partitioning scheme was inferred with PartitionFinder2 v. 2.1.1 (Guindon *et al*. 2010; Lanfear *et al*. 2012, 2017) on the CIPRES Science Gateway, with subsets being asserted *a priori* according to locus and codon position. Introns were included. Models with I+G extensions were excluded from consideration due to undesirable behavior in a model-based framework (Yang 1996). As an alternative *ad hoc* partitioning scheme for the 9,351-bp alignment, I respectively partitioned all exonic loci so that 1^st^-2^nd^ codon positions were placed in their own partition separate from the 3^rd^, and modeled nucleotide substitution in all partitions under GTR+G. Using AMAS (Borowiec 2016), the full 9,351-bp and 9,062-bp molecular alignments were respectively split according to partition scheme(s) for partitioned Bayesian total-evidence inference.

In total-evidence and morphology-only Bayesian phylogenetic analyses, the Mkv model (Lewis 2001) was used to model substitution of morphological character states, albeit with stationary frequencies of character states treated as free parameters (Felsenstein 1981) in order to accommodate asymmetry in character state frequencies. Variation in evolutionary rate among characters was accommodated by drawing rates from a gamma-distributed prior probability distribution (+G), approximated with 8 discrete categories *k*.

All phylogenetic analyses were performed in a Bayesian statistical framework using RevBayes v. 1.0.11 (Höhna *et al*. 2017) compiled on Ubuntu Linux v. 13.04. The following phylogenetic analyses were implemented: one using the 41-character male morphological dataset alone; one using the 9,351-bp molecular dataset alone; two total-evidence analyses using the 9,351-bp molecular alignment, respectively with algorithmic or *ad hoc* partitioning schemes as described above; and a total-evidence analysis using the 9,062-bp molecular alignment, partitioned algorithmically as described above with PartitionFinder2. Each analysis consisted of four independent Markov chain Monte Carlo (MCMC) chains, each run for 50,000 generations. Trees were sampled every 10 generations, with the first 25% of the run being discarded as burn-in. MCMCs with respect to all continuous parameters were considered converged if the effective sample sizes as given in Tracer v. 1.7.1 (Rambaut *et al*. 2018) were ≥200, with sufficiency of MCMC mixing across posterior probability landscapes being qualitatively assessed using traces of the respective log-likelihoods of each parameter across the course of the analysis. Maximum *a posteriori* trees were compiled from this sample of each run, with node support expressed as Bayesian posterior probability (BPP).

### Data Availability and Nomenclature

All nucleotide and morphological data along with PartitionFinder2 configuration files, RevBayes scripts, and output of all phylogenetic analyses, are available at the Dryad Digital Repository (doi:10.25338/B8GP7C). Sequence Read Archives (SRAs) of raw UCE reads, and UCE assemblies, are publicly available on NCBI (Table 2).

This article has been registered in Zoobank (www.zoobank.org). The LSID number is 5F3BECF6-3715-47B3-8D0F-DE7D66E1DA0A.

## Results

Bayesian total-evidence inference of leptanilline phylogeny using the 9,351-bp legacy-locus dataset under the two partitioning schemes resulted in similar topologies, with none of the differences affecting the composition or interrelationship of major clades. All Bayesian total-evidence phylogenies inferred under the *ad hoc* partitioning schemes are provided on Dryad.

Most nodes in these phylogenies were supported with BPP≥0.95. Those nodes supported with BPP≤0.95 were scattered and shallow (Fig. 1), meaning that the interrelationships among all major leptanilline clades are well-resolved. Although the sampling of the Leptanillinae was more extensive than that of Borowiec *et al*. (2019), our inferences were largely congruent. Bayesian total-evidence inference from the 9,062-bp alignment also drew a consilient conclusion (Fig. 2), indicating that the taxonomically biased distribution of missing data in the 9,351-bp legacy-locus dataset does not have an appreciable effect on the backbone of inferred leptanilline phylogeny. Phylogenetic inference from the 9,351-bp alignment alone, and therefore excluding *P. javana*, fully corroborates the conclusions of total-evidence Bayesian phylogenetic inference with high Bayesian posterior probabilities overall, while inference from the morphological dataset alone was insufficient to resolve the phylogeny of the Leptanillinae (see Dryad). All discussion from here on refers to the phylogeny inferred under the partitioning scheme derived with PartitionFinder2 (Fig. 1) for the 9,351-bp molecular alignment, unless otherwise noted.

**Fig. 1.**
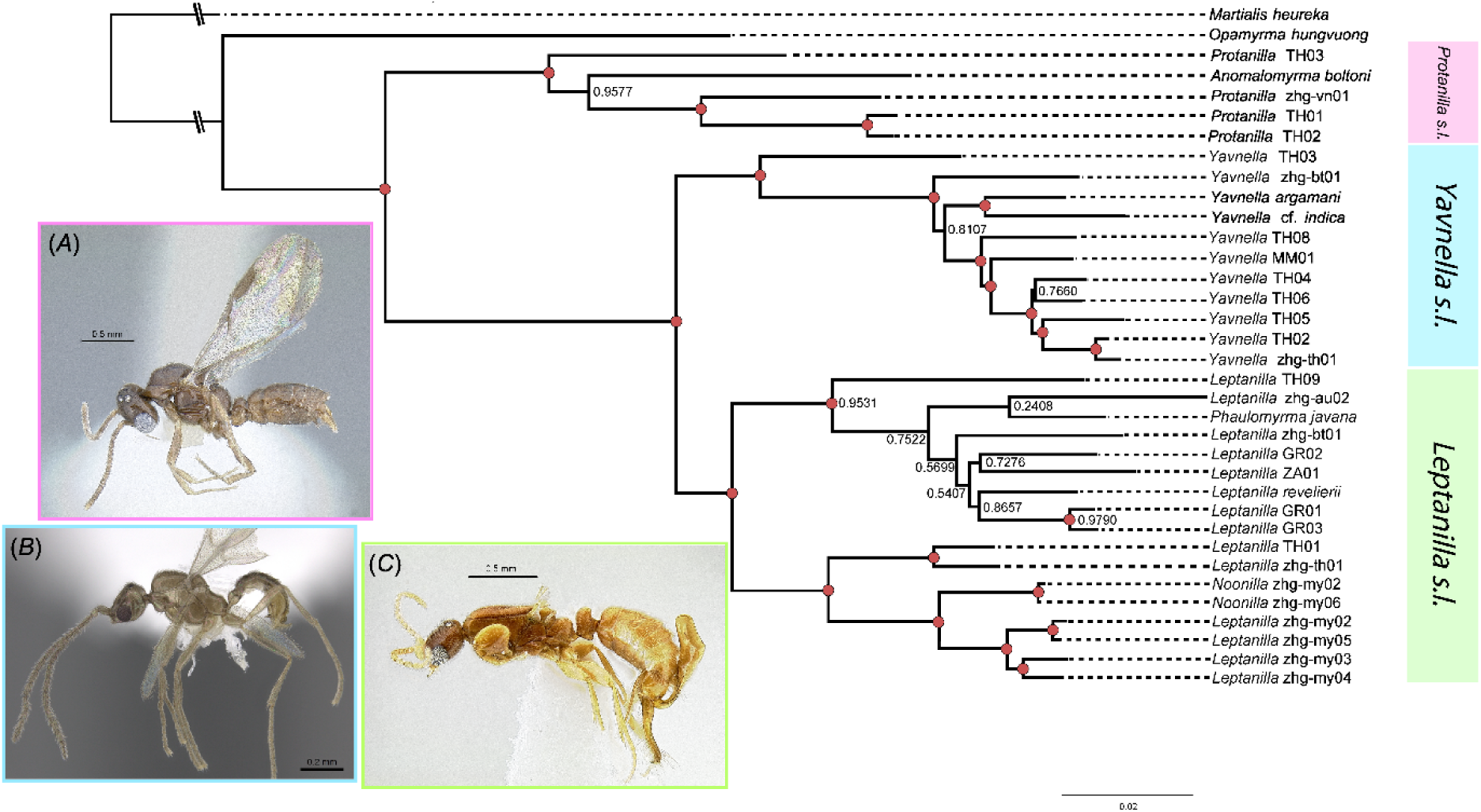
Bayesian total-evidence phylogeny of the Leptanillinae under partitioning scheme inferred with PartitionFinder2 for 9,351-bp legacy-locus alignment. Phylogeny was rooted *a posteriori* on *Martialis heureka*. Nodes with BPP≥0.95 marked in red. (A) *Protanilla* zhg-vn01 (CASENT0842613); (B) *Yavnella* TH08 (CASENT022755; Shannon Hartman); (C) *Leptanilla* zhg-my02 (CASENT0106416).

**Fig. 2.**
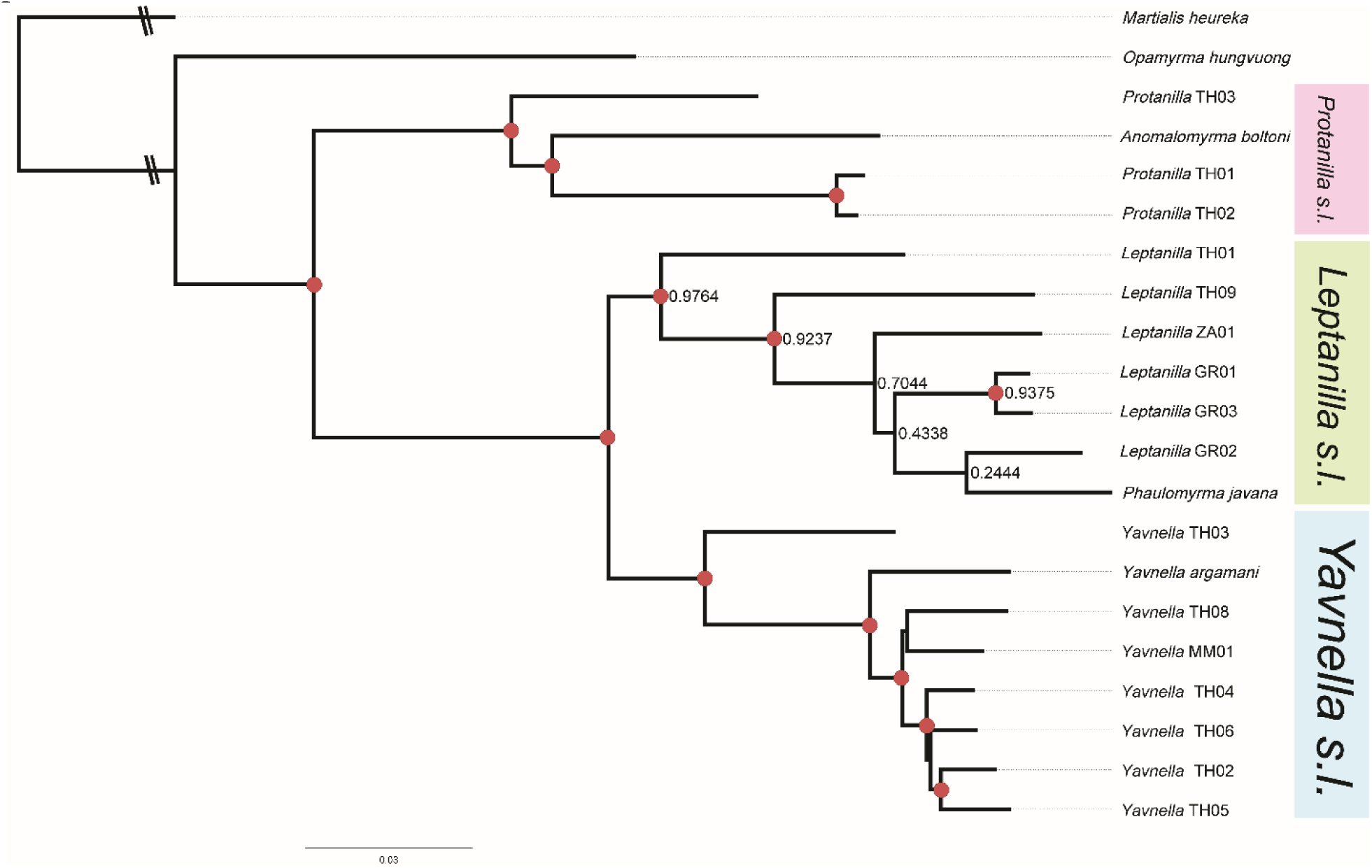
Bayesian total-evidence phylogeny of the Leptanillinae under partitioning scheme inferred with PartitionFinder2 for 9,062-bp legacy-locus alignment. Phylogeny was rooted *a posteriori* on *Martialis heureka*. Nodes with BPP≥0.95 marked in red.

The clade corresponding to the tribe Anomalomyrmini (labeled as *Protanilla sensu lato* in Fig. 1) is recovered with maximal support (BPP = 1), with *A. boltoni* sister to all sampled *Protanilla* save *Protanilla* TH03—thus rendering *Protanilla* paraphyletic—well-supported (BPP = 0.9577). This same topology was recovered by Bayesian total-evidence analysis from the 9,062-bp alignment, with higher support (BPP = 0.9947) (Fig. 2). Borowiec *et al*. (2019) recovered *A. boltoni* as sister to *Protanilla* TH03 with weak support irrespective of statistical framework, albeit with more extensive sampling within the Anomalomyrmini, as did total-evidence inference from the 9,351-bp dataset under the *ad hoc* partitioning scheme (BPP = 0.6535). However, the internal topology of the Anomalomyrmini does not have any bearing upon the status of *Phaulomyrma* relative to other male-based leptanilline genera, nor its status relative to *Leptanilla*.

*Noonilla*, *Yavnella argamani* and *Yavnella* cf. *indica*, *Leptanilla revelierii*, and *Phaulomyrma javana* were firmly recovered within a clade corresponding to the Leptanillini (BPP = 1). As in Borowiec *et al*. (2019) the Leptanillini bifurcate robustly, with *Y. argamani* (and *Yavnella* cf. *indica*, which was not included in Borowiec *et al*. [2019]) recovered in a clade otherwise without described representatives, which is hereinafter designated *Yavnella sensu lato* (BPP = 1). Although morphologically diverse (Fig. 3), the male morphospecies that comprise the sister-group to *Yavnella s. l.* are distinguished from that clade by 1) clypeus with a medial axis no longer than the diameter of the torulus, where the epistomal sulcus is distinct; and 2) pronotum and mesoscutum that are not extended posteriorly in profile view. Since *L. revelierii* is recovered within this clade, it is hereinafter referred to as *Leptanilla sensu lato* (BPP = 0.9964). *Leptanilla s. l.* bifurcates into two well-supported clades: one is broadly Eurasian and Australian in its representation (with a single Afrotropical representative), including *L. revelierii* and *P. Javana* (BPP = 0.9531); the other is Indo-Malayan, and includes *Noonilla* (BPP = 0.9839) (Figs. 1, 4). Since *L. revelierii* is included within the Eurasian-Australian clade, this clade is hereinafter referred to as *Leptanilla sensu stricto.* The two circumscriptions of the name *Leptanilla* presented here are supported by male morphology (see Discussion).

**Fig. 3.**
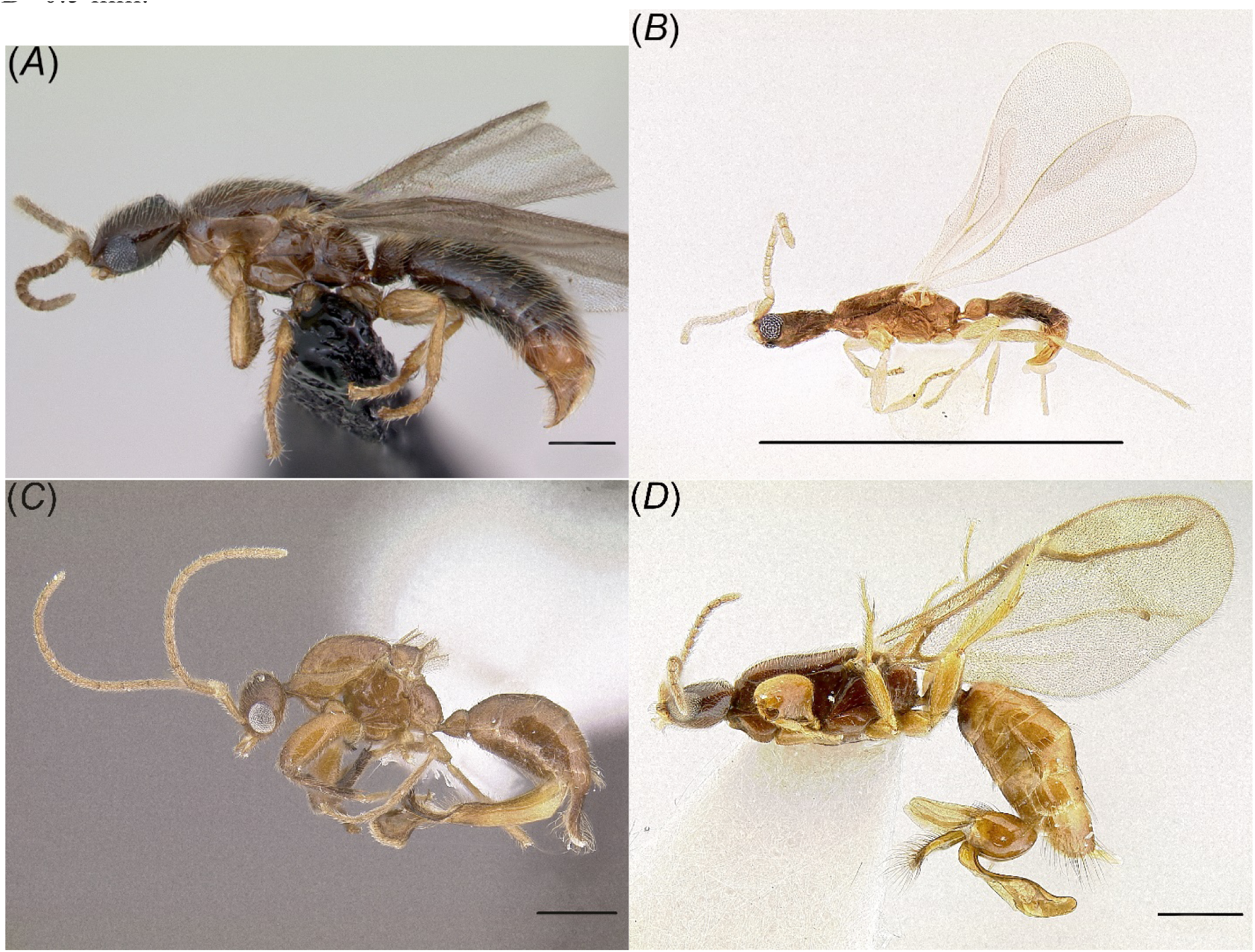
Selected diversity of male *Leptanilla s. l.* (A) *Leptanilla* TH01 (CASENT0119792; April Nobile); (B) *Leptanilla* zhg-bt02 (CASENT084612; not sequenced in this study); (C) *Noonilla* zhg-my04 (CASENT0842610; not sequenced in this study); (D) *Leptanilla* (Bornean morphospecies-group) zhg-my05 (CASENT0842571). Scale bar A=0.2 mm.; B=1 mm.; C-D=0.5 mm.

*Noonilla* (BPP = 0.9999) is sister to a clade represented by highly distinctive male morphospecies, recovered with maximal support (BPP = 1) (Figs. 1, 4), that are immediately recognizable by bizarre metasomal processes (heretofore hypothesized to be extensions of the gonocoxae *sensu* Boudinot [2018] [Boudinot 2015: Fig. 10D]) and a comb-like row of robust bristles on the protibia (Fig. 5) in combination with a putatively grasping profemur. These morphospecies remain undescribed. Boudinot (2015: p. 33) adduced the grasping profemur of *Noonilla* as an autapomorphy of that genus, which could justify terming the undescribed clade as *Noonilla*_cf; but this profemoral condition is more widespread across male Leptanillini than Boudinot (2015) was aware, and better sampling is required to infer whether the grasping profemur is a synapomorphy of *Noonilla* and this undescribed clade. I therefore provisionally refer to said clade as the “Bornean morphospecies-group”: while present sampling is too sparse to judge whether this clade is precinctive to Borneo, available material exclusively originates on that island. Of the 9 terminals recovered in the Indo-Malayan subclade, only *Leptanilla* TH01 was included in Borowiec *et al*. (2019) or in the 9,062-bp legacy-locus alignment. The rather disparate morphospecies *Leptanilla* TH01 and *Leptanilla* zhg-th01 are recovered as a clade with high support (BPP = 0.9961), and this clade is in turn sister to *Noonilla* + Bornean morphospecies-group (Figs. 1, 4). *Leptanilla* zhg-th01 is unique among the Leptanillinae in possessing a recurved mesoscutellar horn (Fig. 6B).

**Fig. 4.**
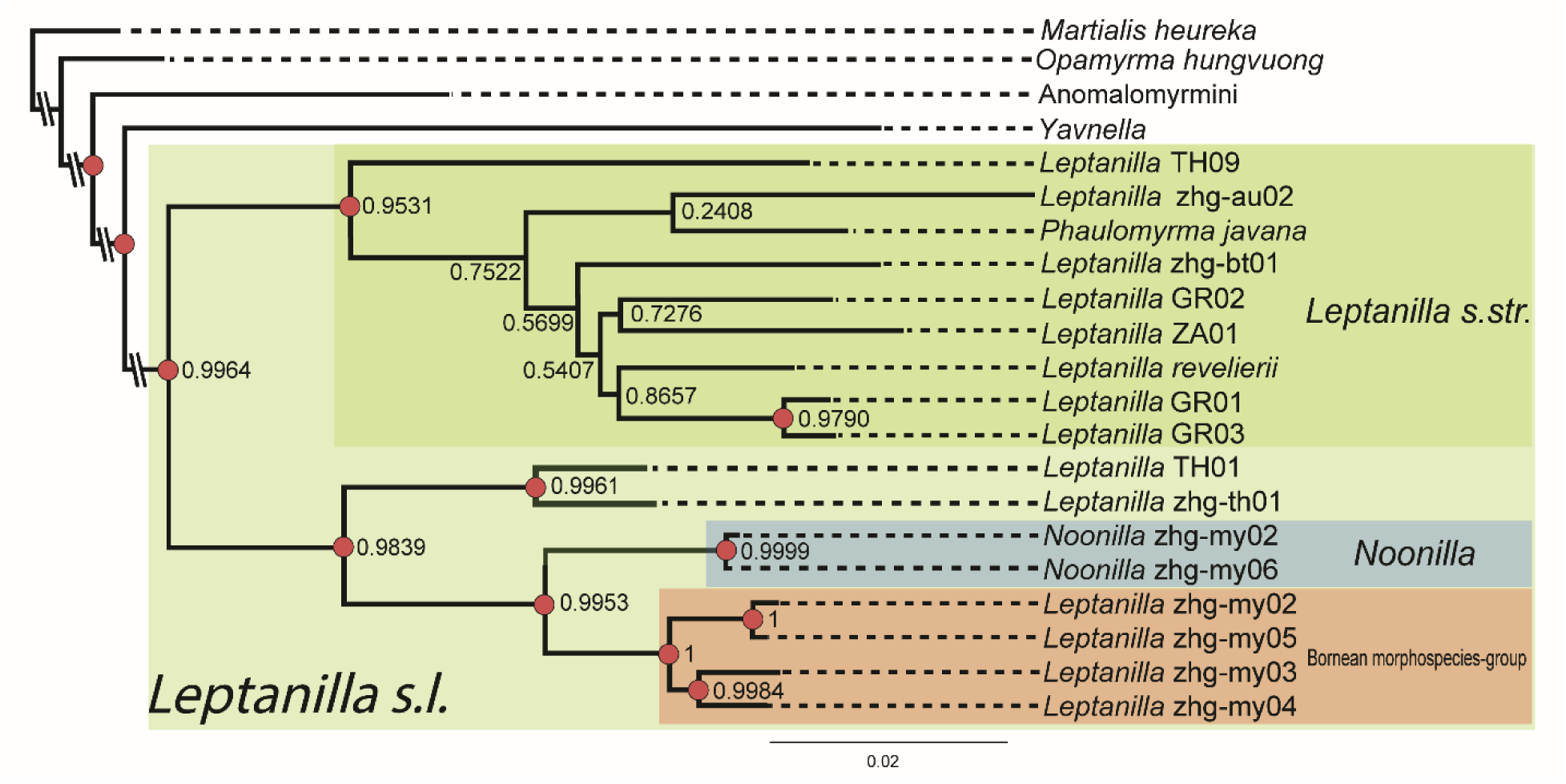
Bayesian total-evidence phylogeny of *Leptanilla s. l.* under partitioning scheme inferred with PartitionFinder2 for 9,351-bp legacy-locus alignment. Phylogeny was rooted *a posteriori* on *Martialis heureka*. Nodes with BPP≥0.95 marked in red.

**Fig. 5.**
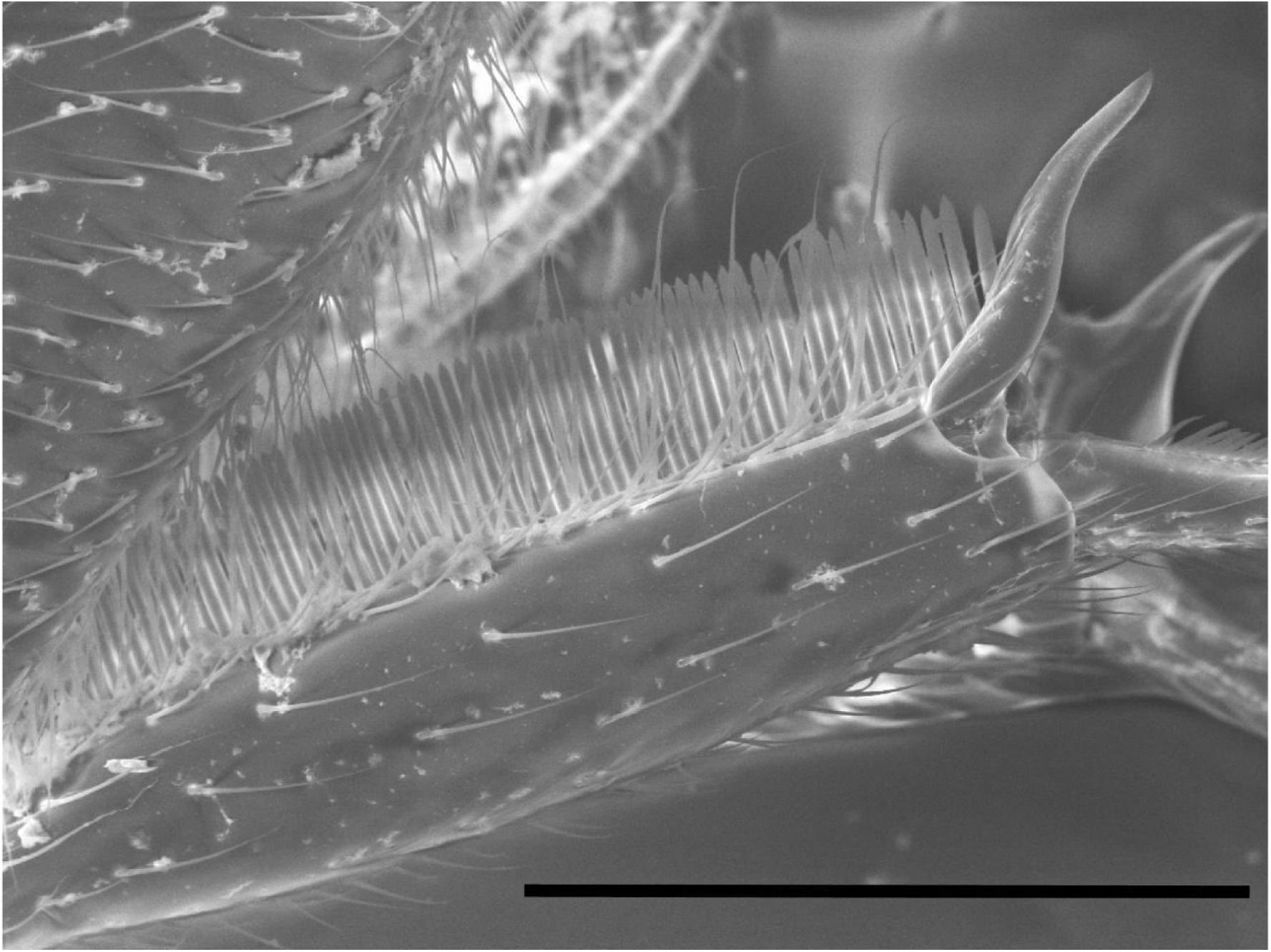
Protibia of *Leptanilla* zhg-my04 (CASENT0842555). Scale bar = 0.2 mm.

**Fig. 6.**
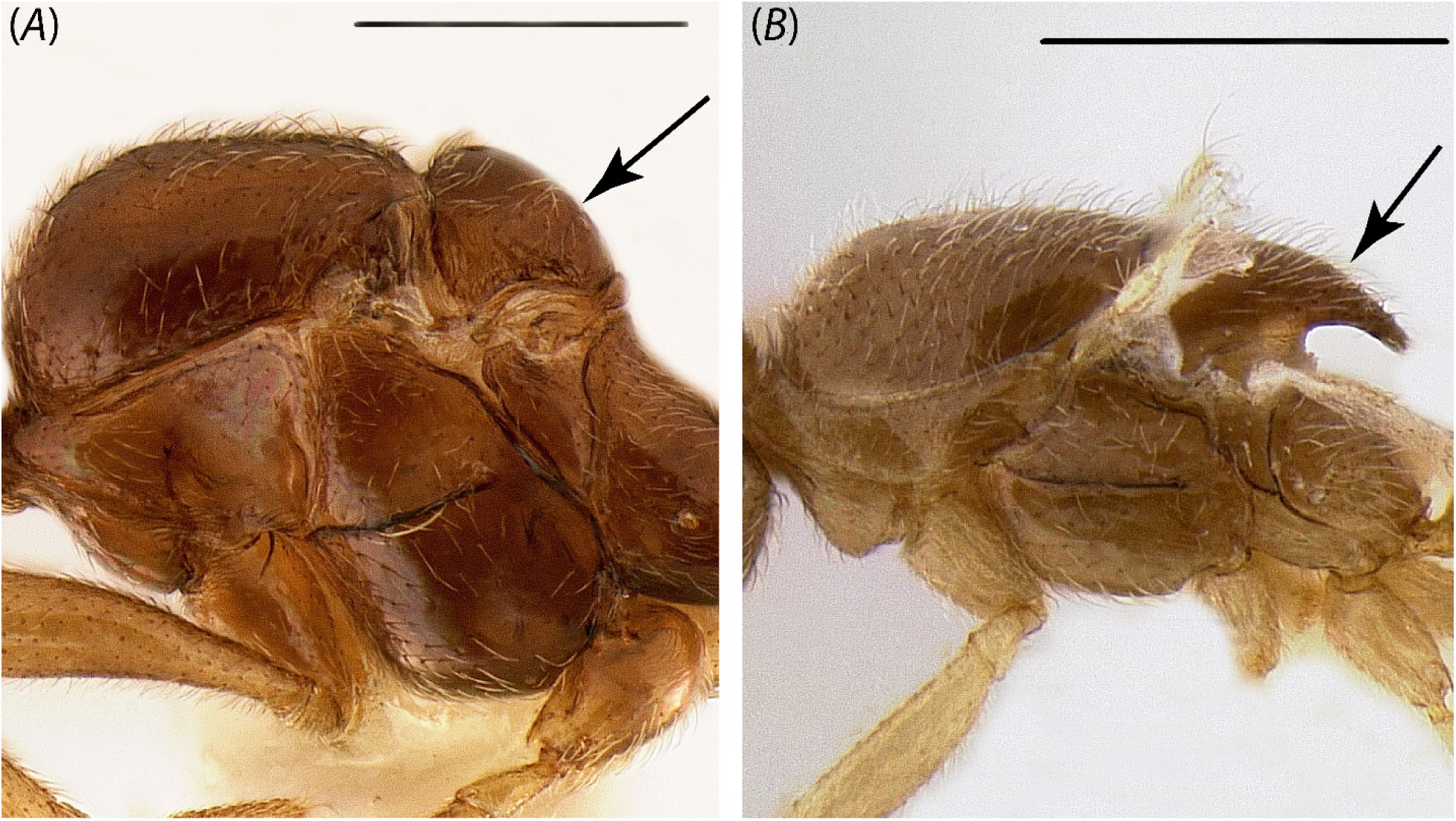
Presence (B: *Leptanilla* zhg-th01; CASENT0842619) versus absence (A: *Yavnella* zhg-th01; CASENT0842621) of the posterior prolongation of the mesoscutellum in male Leptanillini. Scale bar = 0.3 mm.

The support values of internal nodes within *Leptanilla s. str.* are generally poor under Bayesian total-evidence inference from the 9,351-bp legacy-locus alignment, with the placement of *Leptanilla* ZA01 and *Leptanilla* zhg-bt01 differing according to partitioning scheme. The position of *P. javana* cannot be confidently resolved within this clade, but the basalmost node of *Leptanilla s. str.* is well-supported, whether inferred under an algorithmic (BPP = 0.9531) or *ad hoc* (BPP = 0.9585) partitioning scheme. While the internal phylogeny of *Leptanilla s. str.* cannot be resolved with Bayesian total-evidence inference, the monophyly of this clade is probable under the model and partitioning schemes used. The topology of *Leptanilla s. str.* is likely subject to strong stochastic error due to the inclusion of *P. javana*, for which molecular data are entirely absent. This is supported by Bayesian phylogenetic inference from molecular data alone, which with only one exception recovers the internal phylogeny of *Leptanilla s. str.* with BPP≥0.95 (see Dryad).

Bayesian total-evidence inference from the 9,062-bp alignment (which does not include *Leptanilla revelierii*, zhg-au02 or zhg-bt01) gives mediocre support to *Leptanilla s. str.* (BPP = 0.9237) inclusive of *P. javana*, but provides a phylogeny consilient with the results of other phylogenetic analyses (Fig. 2). The recovery of *P. javana* within *Leptanilla s. str.* is therefore supported by Bayesian total-evidence inference. Qualitatively, male morphological characters support *Leptanilla s. str.* (see Discussion).

*P. javana* and the taxon dubbed *Phaulomyrma* MM01 by Boudinot (2015) and Borowiec *et al*. (2019) were recovered distant from one another in the leptanilline phylogeny (Figs. 1-2, 4). Total-evidence phylogenetic inference recovered the latter terminal within *Yavnella s. l.*, indicating that it was incorrectly assigned to *Phaulomyrma* by these authorities, corroborating morphological evidence (see Discussion). An undescribed male morphospecies referred to as *Phaulomyrma* by Boudinot (2015: Fig. 4F) was not sequenced in this study but also conforms morphologically to *Yavnella s. l*., and so likewise was incorrectly identified as *Phaulomyrma*. Conversely, *P. javana* is here recovered within *Leptanilla s. l.*, and moreover within *Leptanilla s. str.* (BPP = 0.9531).

## Discussion

### Delimitation of Subclades in Leptanillinae using Male Morphology

Male morphological characters corroborate inferred phylogeny at nodes of variable depth. *O. hungvuong* and the four male representatives of the Anomalomyrmini included in the present study can easily be distinguished from male Leptanillini by the presence of a pterostigma (although wing venation may be inaccessible due to deciduous wings in some male Leptanillini [pers. obs.]) and the absence of an ocellar tubercle. Griebenow (in press) provides a formal description of female-associated male *Protanilla* and a male-based definition of the leptanilline tribes, as well as *O. hungvuong*. *Yavnella s. l.* is likewise well-supported (Figs. 1-2), as is *Leptanilla s. l.*, with the former clade diagnosed almost entirely by morphological symplesiomorphies: the only putative autapomorphy of *Yavnella s. l.* is concavity of the propodeum in profile view (Fig. 7A), which was previously noted by Kugler (1986) as being distinctive to *Yavnella*.

**Fig. 7.**
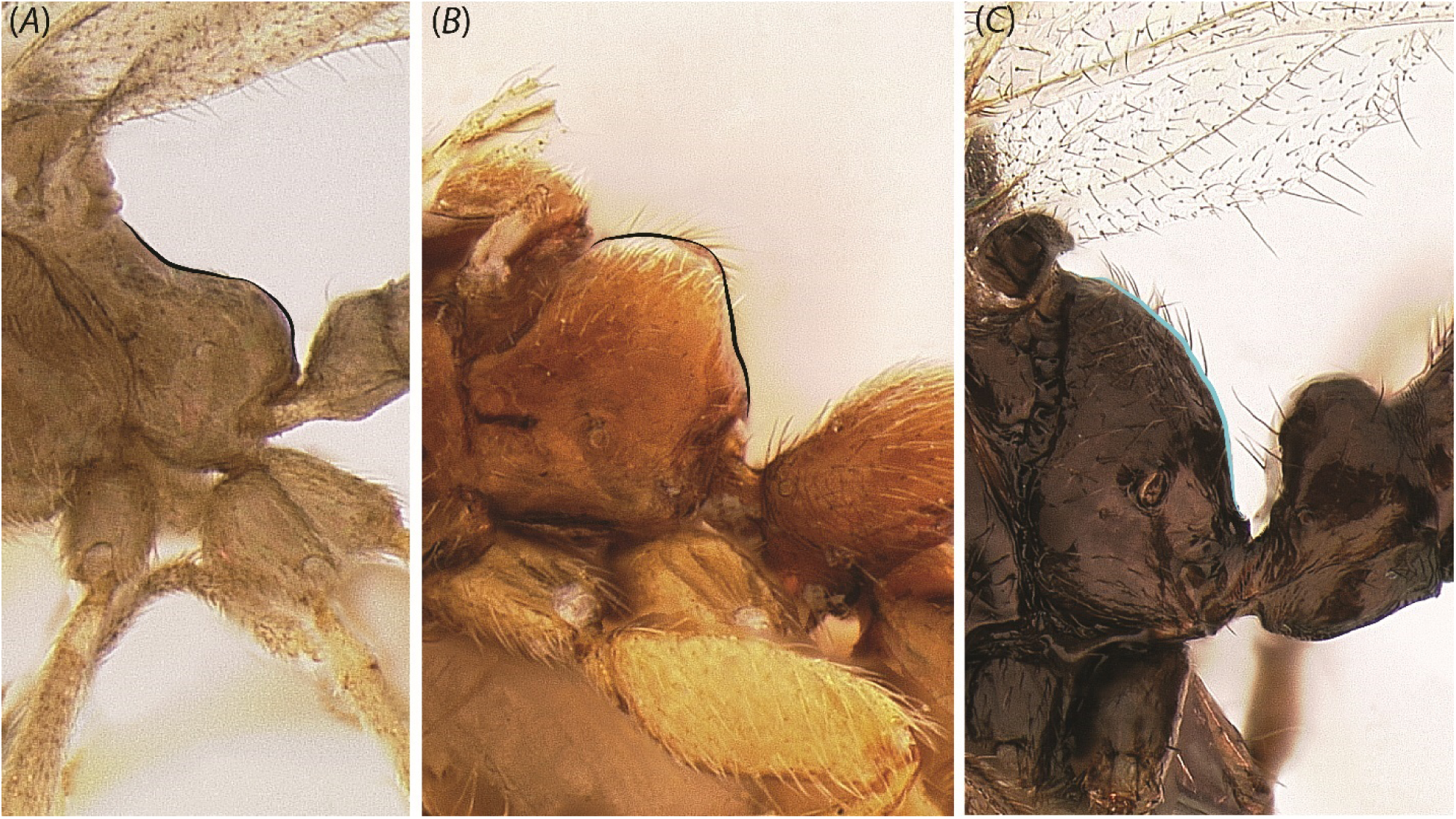
Conditions of the propodeum in the Leptanilllinae. (A) Concave (*Yavnella* zhg-bt01; CASENT0106384); (B) convex with distinct dorsal face (*Leptanilla* zhg-my02; CASENT0106456); (C) convex without distinct dorsal face (*Protanilla lini* [OKENT0011097]; male described by Griebenow, in press) (not sequenced in this study).

*Leptanilla s. str.* is identifiable relative to other subclades of *Leptanilla s. l.* based upon the following combination of male morphological characters: absence of posterior mesoscutellar prolongation (observed in *Leptanilla* zhg-th01 and *Leptanilla* TH01); propodeum convex and without distinct dorsal face (Fig. 7C); gonopodites articulated (otherwise among the Leptanillini articulated only in *Leptanilla* zhg-th01 and some *Noonilla*); gonocoxae fully separated ventrally (this character state [Fig. 8A] elsewhere observed among sampled Leptanillini in all *Yavnella s*. *l.* except for Yavnella TH03; and *Leptanilla* zhg-th01); and penial sclerites dorsoventrally compressed along their entire length, entire, and lacking sculpture (Fig. 9A), this character state elsewhere observed in Leptanillini only among *Yavnella s. l.* (excluding *Yavnella* TH03) and *Leptanilla* zhg-th01.

**Fig. 8.**
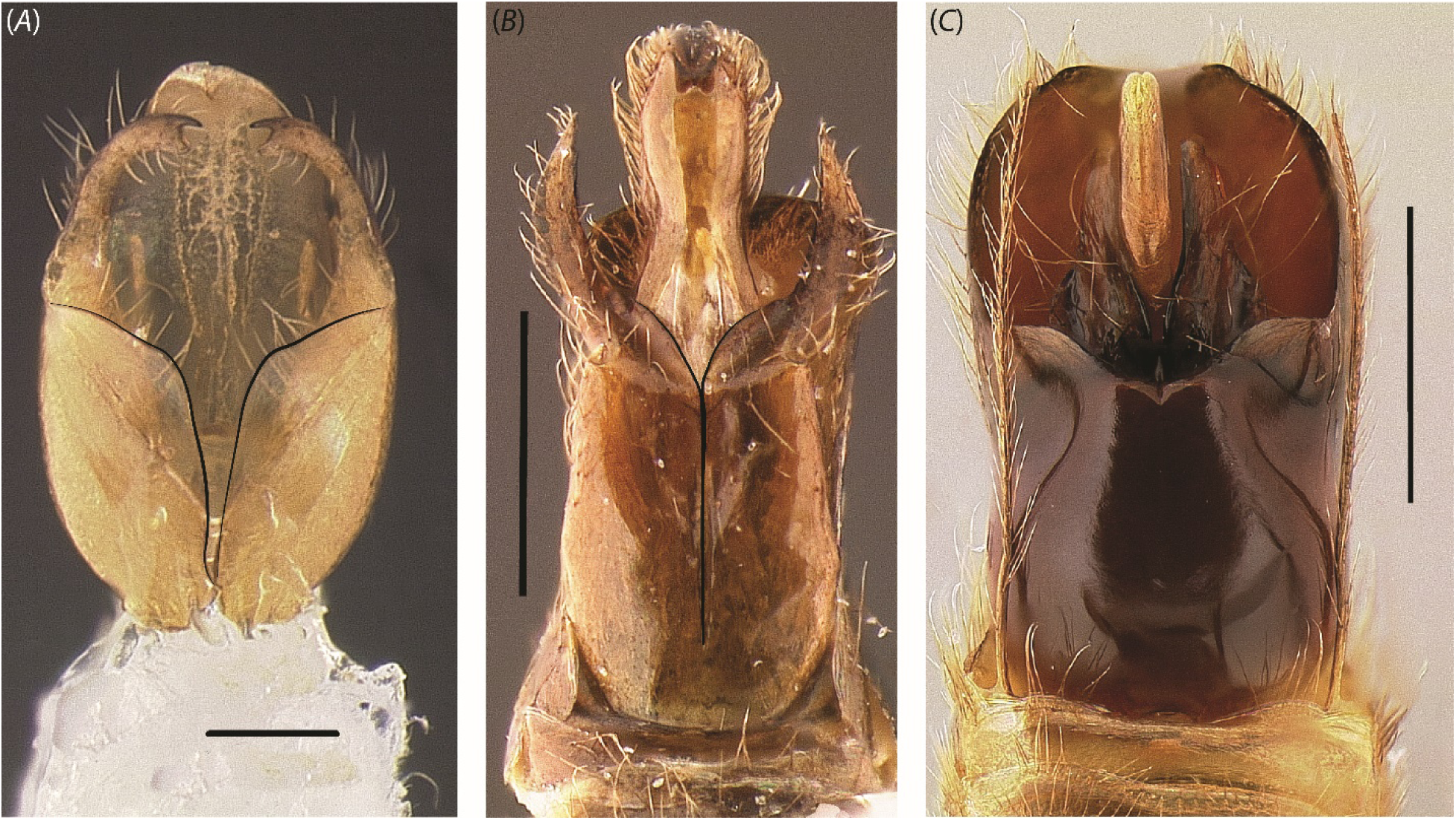
Ventral view of male genitalia across the Leptanillini. (A) *Leptanilla* ZA01 (CASENT0106354), (B) *Noonilla* zhg-my02 (CASENT0842595); (C) *Leptanilla* zhg-my04 (CASENT0842553). Scale bar A = 0.1 mm.; B = 0.3 mm.; and C = 0.5 mm.

**Fig. 9.**
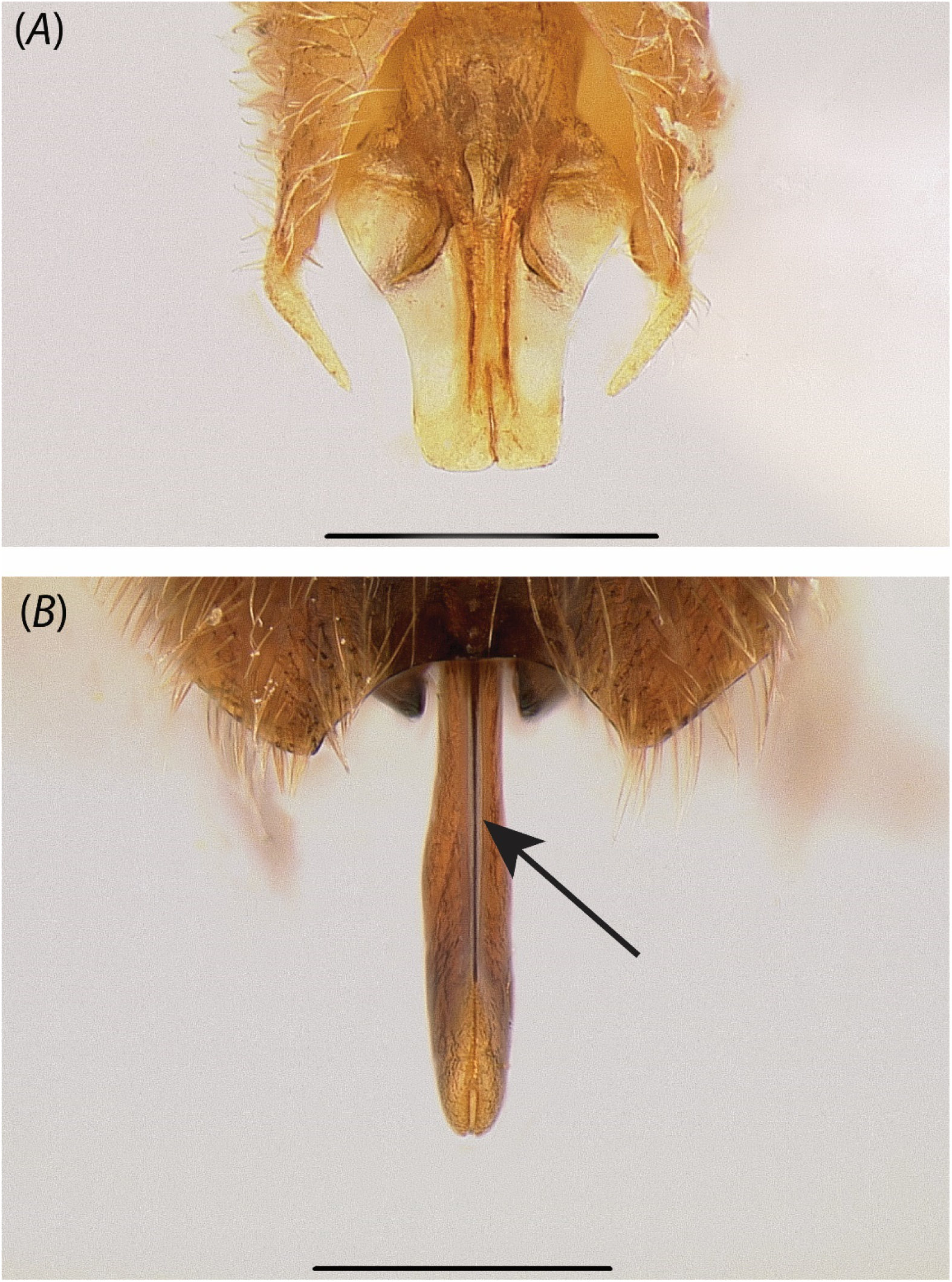
Dorsoventral (A) (*Yavnella* zhg-th01; CASENT0842620) vs. lateromedial (B) (*Leptanilla* zhg-my04; CASENT0842553) compression of the penial sclerites in posterodorsal view. Dorsomedian carina marked with arrow. Scale bar = 0.3 mm.

*Leptanilla* TH09 is weakly recovered as sister to remaining *Leptanilla s. str.*, including *P. javana*, under all Bayesian total-evidence analyses (Figs. 1-2, 4). Therefore, the phylogeny of *P. javana* relative to other *Leptanilla s. str.* would not be resolved if that clade were delimited to exclude *Leptanilla* TH09. However, *Leptanilla* TH09 conforms fully to the diagnosis of *Leptanilla s. str.* given above, and aside from apomorphies of the foreleg (a perhaps opposable calcar and apical probasitarsal seta; Fig. 10A) is not a phenotypic outlier among the terminals representing *Leptanilla s. str.* Nor given the weak BPP of *Leptanilla* TH09 as sister to the remainder of *Leptanilla s. str.* is there probabilistic support for qualitatively defining that clade to exclude *Leptanilla* TH09. Therefore, *P. javana* can be confidently placed within *Leptanilla s. str.*, despite the inability of Bayesian total-evidence inference from these data and under these models to resolve its position within that clade.

**Fig. 10.**
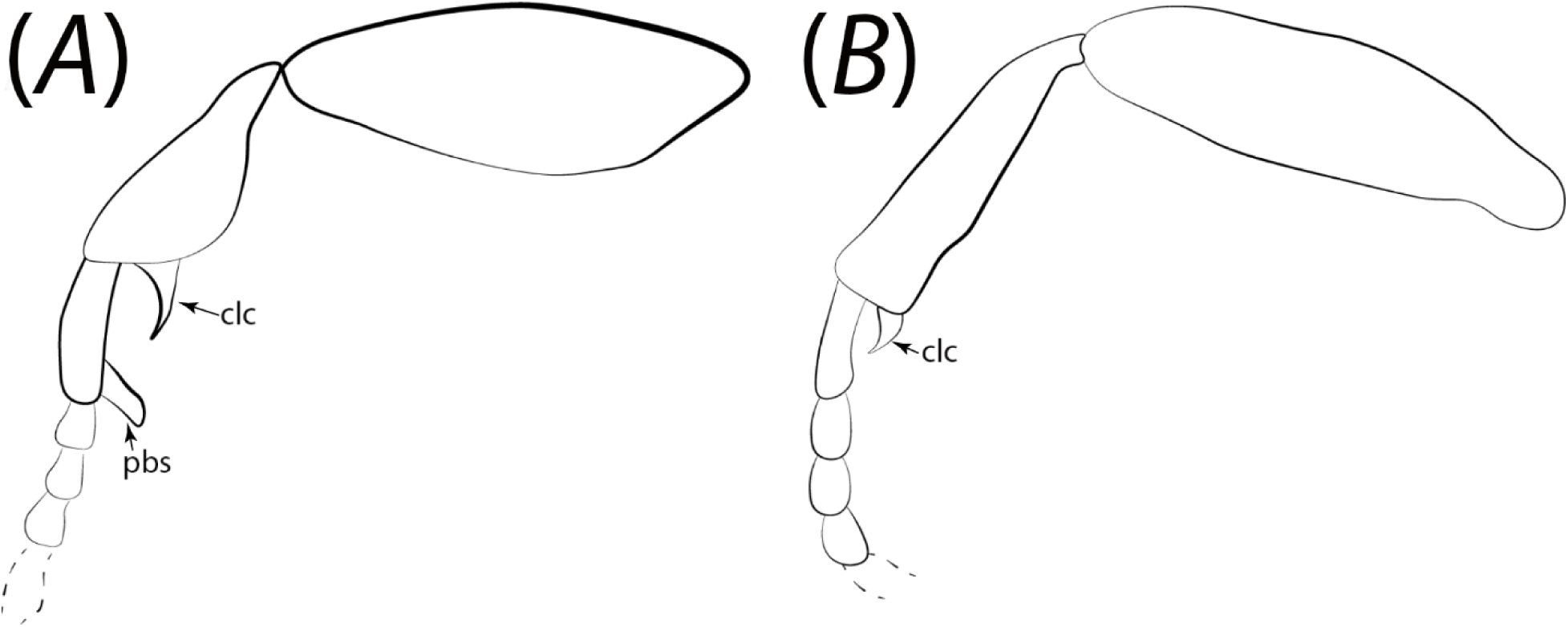
Profemur, protibia, and basal protarsomeres of (A) *Leptanilla* TH09 (CASENT0842664) and (B) *Leptanilla* zhg-bt01. Abbreviations: clc = calcar; pbs = probasitarsal seta. Not to scale.

Unlike *Scyphodon*, *Noonilla*, and even the male-based species *Leptanilla palauensis* Smith, 1953 (Petersen, 1968: p. 593), the status of *Phaulomyrma* as a leptanilline—and as an ant—has never been debated. Wheeler & Wheeler (1930) established the genus based upon the presence of wing veins and “unusually large genitalia” (Wheeler & Wheeler 1930: p. 193), transferring also *Leptanilla tanit* Santschi, 1907 to *Phaulomyrma*. Their argument regarding wing venation has no merit, given that the forewing venation of *P. javana* falls within the range of variation observed in putative *Leptanilla* morphospecies (Petersen 1968: pp. 594-595), with all leptanilline males examined by Boudinot (2015) exhibiting at least one compound abscissa on the forewing.

Petersen (1968: p. 597) even referred to *Leptanilla* and *Phaulomyrma* as “nearly identical” (when comparing these taxa to *L. palauensis*), and returned *L. tanit* to *Leptanilla*, but refrained from synonymizing *Leptanilla* and *Phaulomyrma* on account of the apparent uniqueness of the genitalia of *P. javana* as illustrated by Wheeler and Wheeler (1930: Figs. 2A, C). In passing, Taylor (1965: p. 365) also mentioned *Phaulomyrma* as being “possibly synonymous” with *Leptanilla*.

Examination of a syntype of *P. javana* (lectotypified below) demonstrates that its genitalia are consistent with other sampled male *Leptanilla s. str.* to the exclusion of males within the Indo-Malayan sister-group of *Leptanilla s. str.* (Fig. 11). Although the preservation of this specimen on a slide prevents us from directly confirming stylar articulation, the sharply recurved styli are consistent with the syndrome seen in dried male leptanillines with articulated gonopodites (Kugler 1986; Ward and Sumnicht 2012), indicating that the gonopodites are articulated in *P. javana*. *Contra* Fig. 2C of Wheeler and Wheeler (1930), the volsellae of *P. javana* are not discernible *in situ* (Fig. 11D). If their condition is truly “plate-like” as described by Wheeler and Wheeler (1930: p. 196), the volsellae of *P. javana* resemble those observed in undescribed Sicilian male morphospecies attributed to *Leptanilla* (Scupola & Ballarin 2009). Dissection of Anatolian *Leptanilla* GR03, and Spanish material that closely resembles sequenced males of *Leptanilla s. str.*, demonstrates that the volsellae are likewise lamellate in these morphospecies, having much the same condition as in *Leptanilla africana* (Baroni Urbani 1977: Fig. 37) (not included in this study). Therefore, given the phylogeny of *P. javana* and its morphological conformity to *Leptanilla s. str.* there is no justification for maintaining the genus *Phaulomyrma*. It is a nomenclatural irony that Wheeler & Wheeler (1930: p. 193) note that the derivation of the genus name is from the Greek *phaulus*, which they translate as “trifling or paltry”: the justification for establishing *Phaulomyrma* as a genus was trifling indeed.

**Fig. 11.**
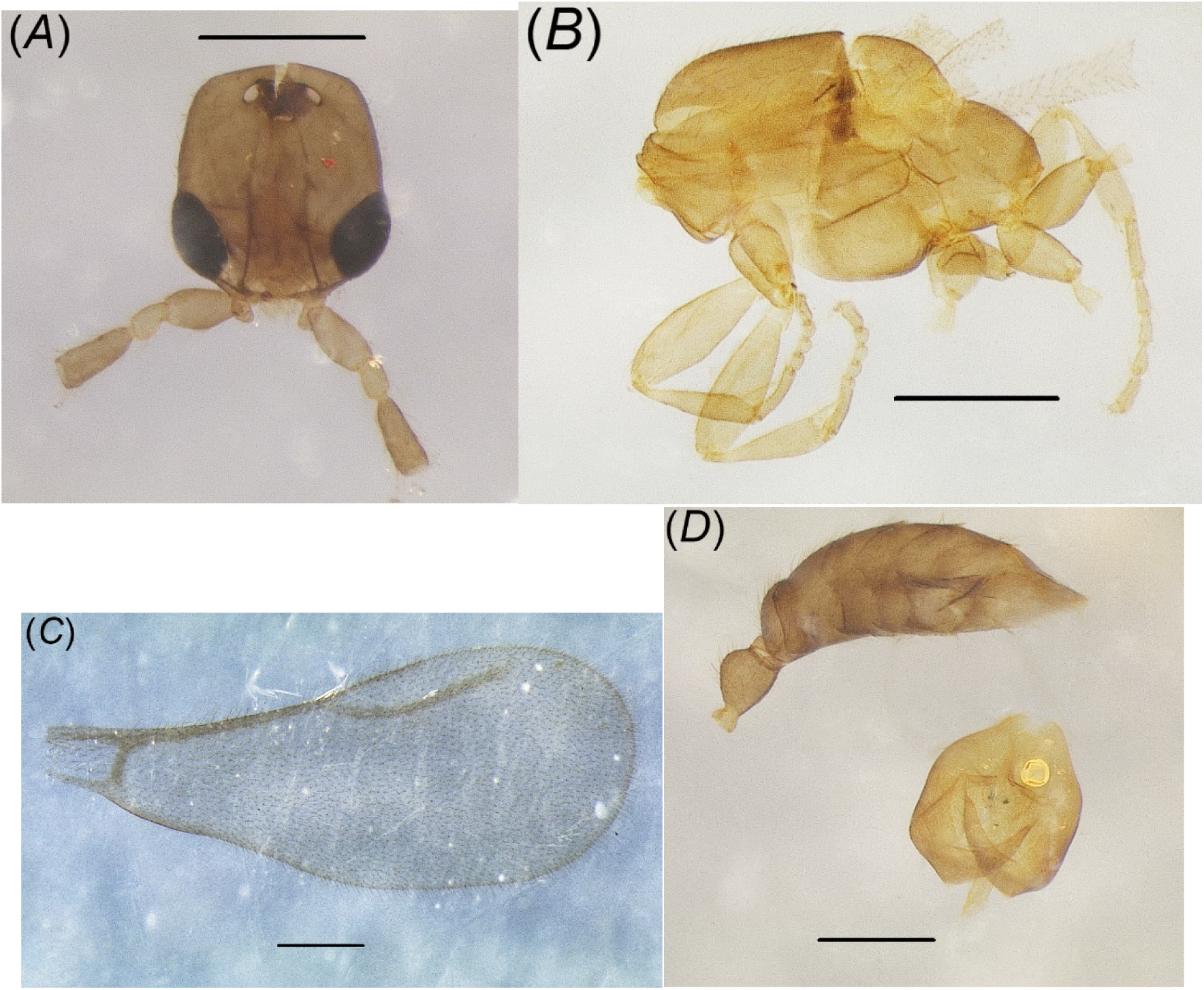
Lectotype of *Phaulomyrma javana* as designated by this study (MCZ:Ent:31142). (A) Full-face view; (B) profile view of mesosoma; (C) forewing; (D) metasoma and genitalia. Scale bar = 0.2 mm.

A complete male-based diagnosis of *Leptanilla* relative to other Leptanillinae under both broad and strict circumscriptions of *Leptanilla* is provided below, with putative synapomorphies for the two circumscriptions represented in italics. Only genital characters could be scored for *Leptanilla* ZA01.

> ***Leptanilla javana*** (Wheeler & Wheeler, 1930) comb. nov.
>
> Figs. 11A-D.
>
> *Phaulomyrma javana* Wheeler & Wheeler 1930: 193. Figs. 1, 2C.
>
> *Phaulomyrma javana –* Petersen 1968: 293. Figs. 16A-C.

### Lectotype

INDONESIA • ♂; Jawa Barat, “Buitenzorg” [Bogor]; Mar. 1907; F.A.G. Muir leg.; MCZ 31142.

### Paralectotype

Same data as for lectotype (no accession code).

> **Genus *Leptanilla* Emery, 1870**
>
> Type species: *Leptanilla revelierii* Emery, 1870: 196.

= *Leptomesites* Kutter, 1948 (286). Synonymy by Baroni Urbani, 1977 (433). Holotype deposited at MHNG (Muséum d’Histoire Naturelle, Geneva).

= *Phaulomyrma* Wheeler & Wheeler, 1930 (193); **syn. nov.** Lectotype and paralectotype deposited at MCZC (Museum of Comparative Zoology, Cambridge, Massachusetts).

*Male diagnosis of* Leptanilla s. l. *relative to other Leptanillinae*

1. Mandibles articulated to gena (Fig. 12B).
2. *Medial axis of clypeus no longer than diameter of torulus, when epistomal sulcus is distinct*.
3. Antennomere 3 shorter than scape.
4. Ocelli present and set on tubercle (Fig. 13) (with exception of *Leptanilla* [Bornean morphospecies-group] zhg-my05).
5. *Pronotum and mesoscutum posteriorly extended (Fig. 14B-C)*.
6. Notauli absent.
7. Pterostigma absent.
8. Propodeum not concave in profile view.

**Fig. 12.**
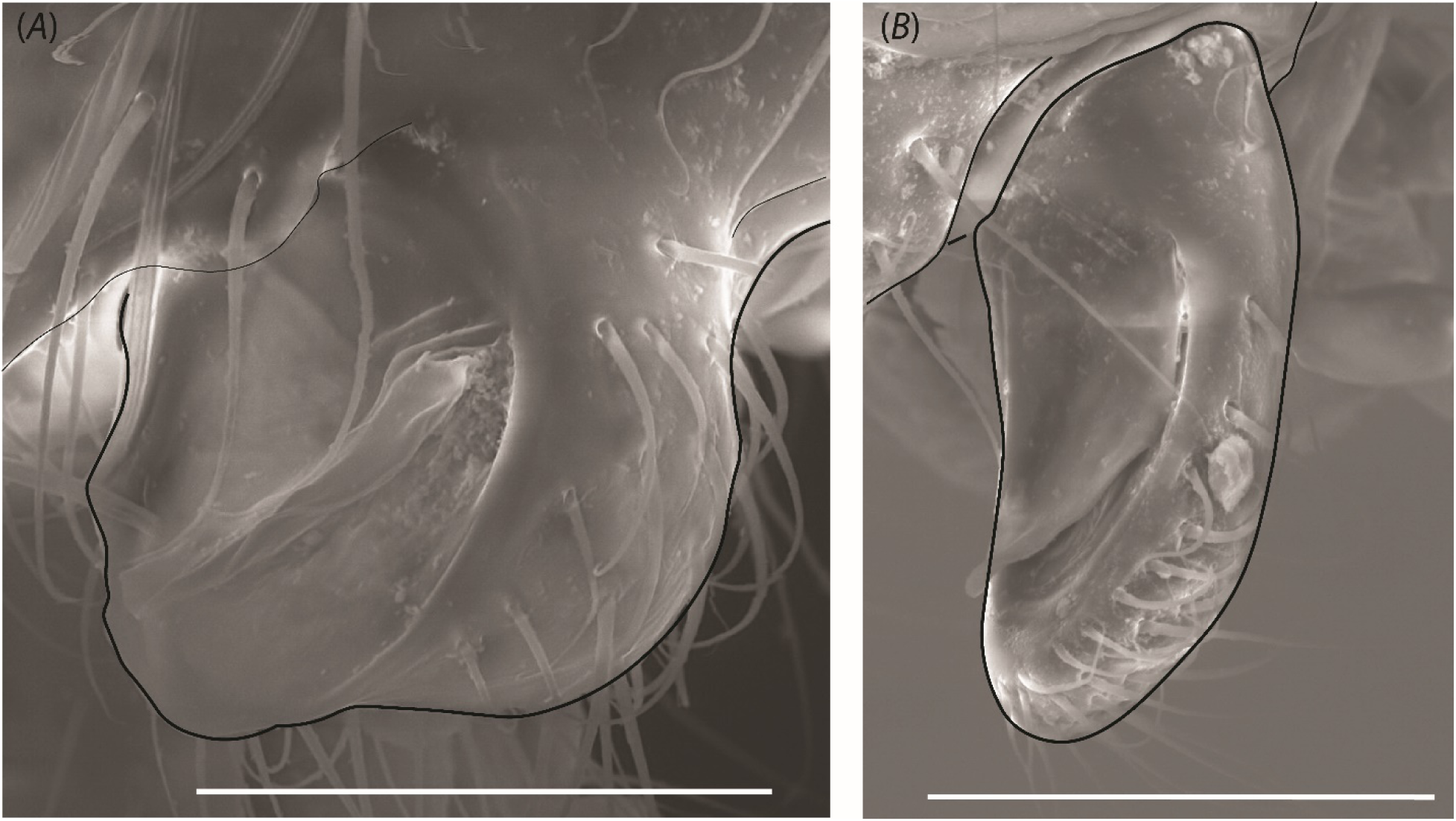
Mandible of (A) *Yavnella* cf. *indica* (CASENT0106377) and (B) *Leptanilla* zhg-my03 (CASENT0842618). Scale bar A=0.03 mm.; B=0.05 mm.

**Fig. 13.**
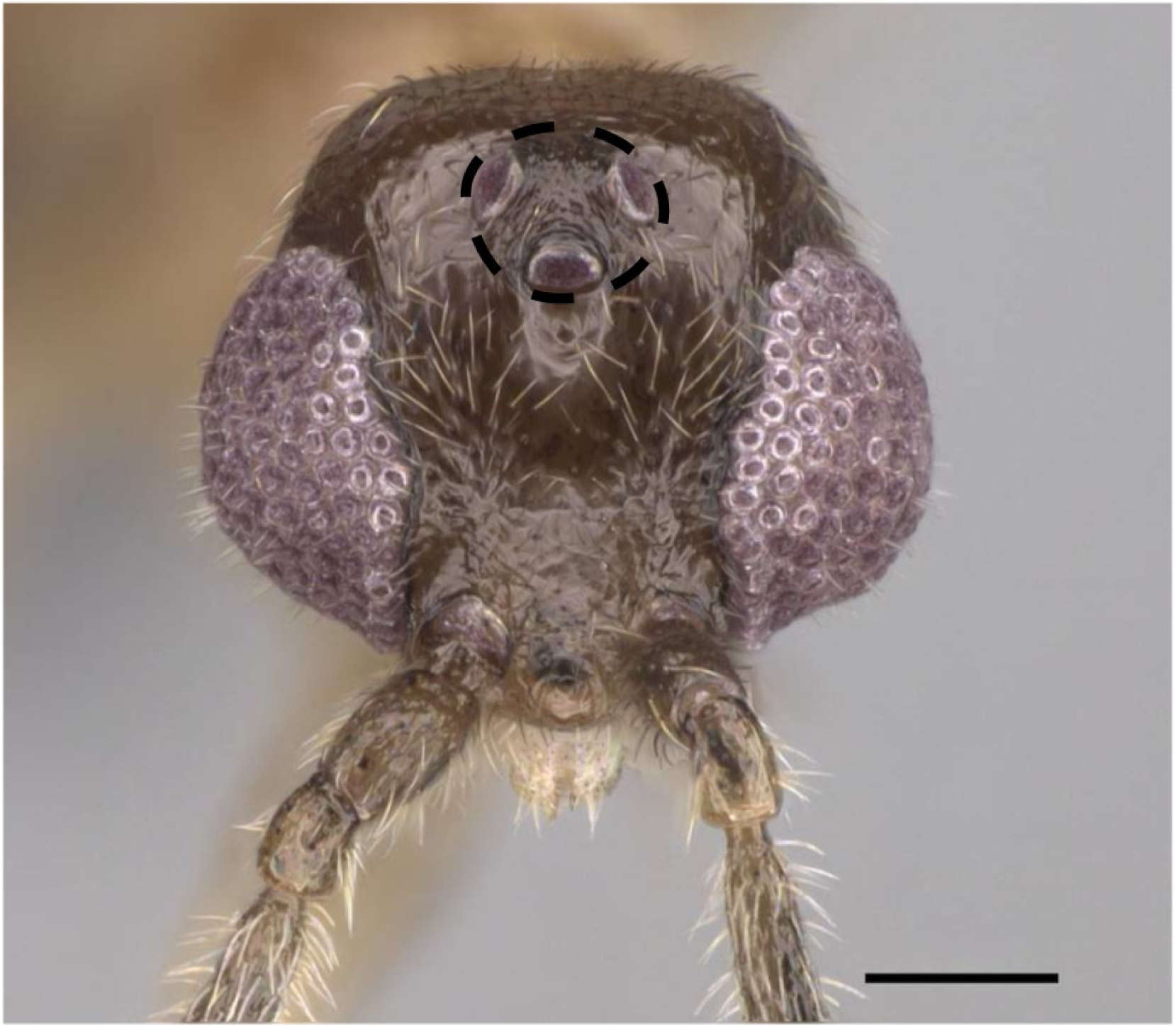
Full-face view of *Yavnella* TH02 (CASENT0119531; Michele Esposito), with ocellar tubercle marked. Scale bar=0.1 mm.

**Fig. 14.**
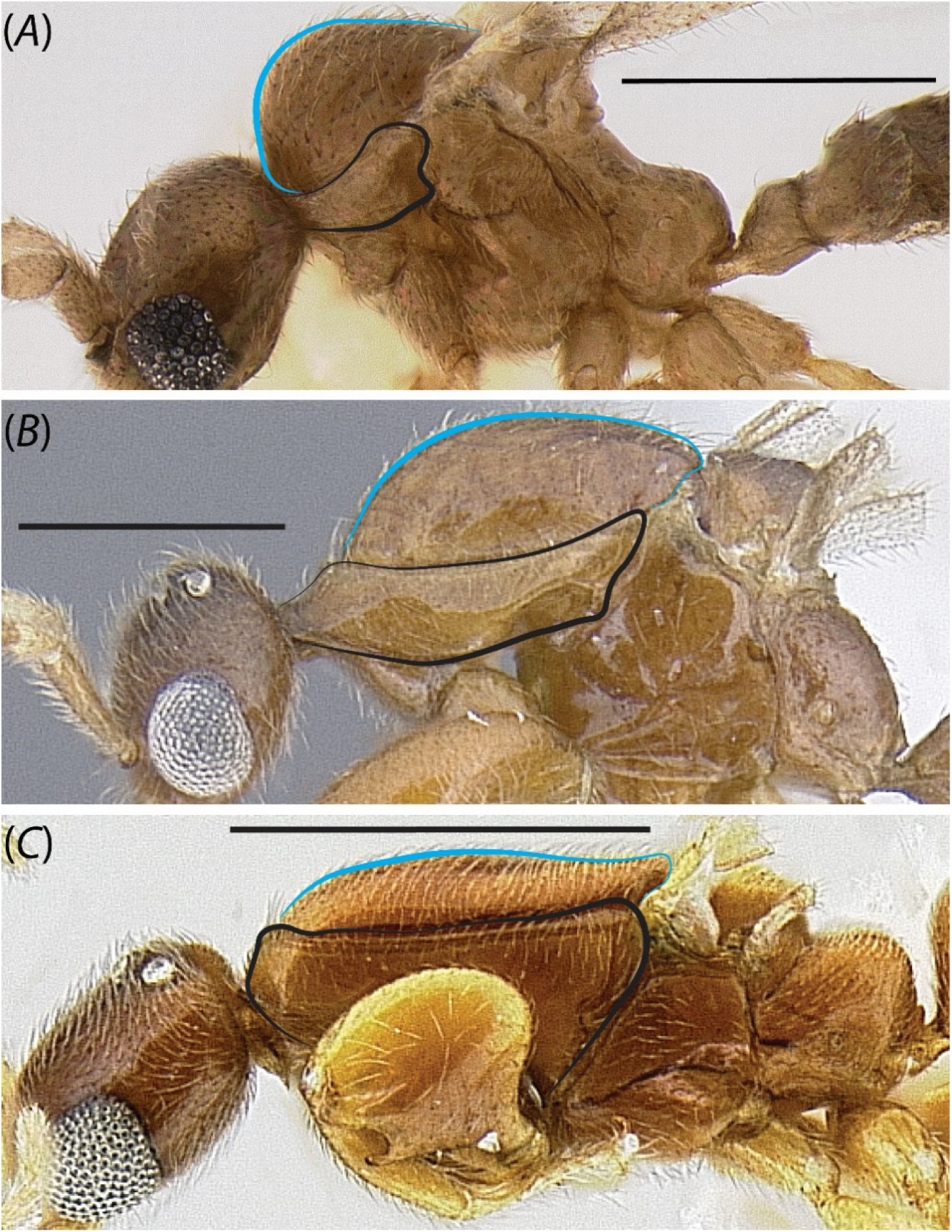
Profile of pronotum (black) and mesoscutum (blue) in male Leptanillini. (A) *Yavnella* zhg-bt01 (CASENT0106384); (B) *Noonilla* zhg-my04 (CASENT0842610; not sequenced in this study); (C) *Leptanilla* (Bornean morphospecies-group) zhg-my03 (CASENT0106416).

*Male Diagnosis of* Leptanilla s. str. *relative to other* Leptanilla s. l.

9. Anteromedian ocellus and compound eye not intersecting line parallel to dorsoventral axis of cranium.

10. Profemoral ventral cuticular hooks absent.

11. Ventromedian protibial comb-like row of setae absent.

12. Infuscation at juncture of Rf and 2s-rs+Rs+4-6 absent.

13. *Antero-admedian line absent* (HAO: 0000128).

14. Mesoscutellum not posteriorly prolonged.

15. Propodeum convex in profile view, without distinct dorsal face.

16. Abdominal sternite IX without posterolateral filiform processes.

17. Abdominal tergite VIII broader than long.

18. Gonocoxae medially separated*.

19. Gonopodites articulated.

20. *Volsella lamellate, entire distally, without denticles*\*.

21. Penial sclerites dorsoventrally compressed, dorsomedian carina absent, ventromedian carina sometimes present.

22. Phallotreme situated at penial apex, without vestiture.

*These character states observed so far as is possible with available specimens.

*Notes*.
1. The mandibles are fused to the gena (Fig. 12A) in sampled *Yavnella s. l.* except for *Yavnella* TH04: this character state is seen elsewhere in male ants only among some *Simopelta* spp. (Ponerinae) (Brendon Boudinot, pers. comm.).
2. The epistomal sulcus is often difficult to distinguish in *Leptanilla s. l.*, but the anteroposterior reduction of the clypeus can be inferred by the situation of the toruli at the anterior-most margin of the head (cf. Boudinot 2015: p. 30).
3. Antennomere 3 is longer than the scape in all sampled *Yavnella s. l.* except for *Yavnella* TH05.
4. Ocelli are entirely absent in *Yavnella* TH03 and *Yavnella* zhg-bt01. The ocellar tubercle is absent in the Anomalomyrmini and *O. hungvuong*; within *Leptanilla s. l.* it is absent in *Leptanilla* zhg-my05, which is here inferred to be a secondary loss.
5. As noted by Petersen (1968: p. 87), *N. copiosa* contrasts with other described male Leptanillinae by the lack of an “elongated, laterally compressed” mesosoma. *Yavnella* was described by Kugler (1986) as sharing this condition, which Petersen (1968) adduced as plesiomorphic for the Leptanillinae. While the relative modification of the mesosoma—here approximated by the proportions of the pronotum and mesoscutum— forms a morphocline across the male Leptanillinae, this morphocline is discontinuous, with a gap between the morphospace occupied by *Leptanilla s. l.* (Fig. 14B-C) and that occupied by *O. hungvuong*, the Anomalomyrmini, and *Yavnella s. l.* (Fig. 14A). Future sampling of male Leptanillinae may close this gap in morphospace, which would limit the diagnostic utility of pronotal and mesonotal length.
6. The absence of notauli is a synapomorphy of the tribe Leptanillini. The notauli in *Protanilla* TH01 and *Protanilla* zhg-vn01, in the tribe Anomalomyrmini, are homoplastically absent.
7. The absence of the pterostigma (Figs. 15A, C) is a synapomorphy of the Leptanillini.
8. The convexity of the propodeum in profile view is plesiomorphic for the Leptanillinae. Its concave condition in *Yavnella* (Kugler 1986) is apomorphic for that genus.
9. The anteromedian ocellus is not situated orthogonally to the compound eye in profile view in *Leptanilla s. str. Leptanilla* TH01 and zhg-th01, the Bornean morphospecies-group, and all examined *Noonilla*. The concomitant prognathy of the male cranium is unique among male Leptanillinae to *Leptanilla s. l.*, and as adduced by Petersen (1968), this condition appears apomorphic among the Leptanillinae.
10. A profemoral ventral cuticular hook (Fig. 16B) is unique among the morphospecies sampled herein to *Leptanilla* (“Bornean morphospecies-group”) zhg-my02 and −05.
11. The ventromedian comb-like row of setae on the protibia is an autapomorphy of the Bornean morphospecies-group.
12. The infuscation observed in the Bornean morphospecies-group at the juncture of Rf and 2s-rs+Rs+4-6 (Fig. 15C) is not enclosed anteriorly by an abscissa and appears to be homoplasious with the pterostigma observed in male Anomalomyrmini. Infuscation of the forewing is otherwise absent in the Leptanillini.
13. The antero-admedian line is present among sampled Leptanillini only among some *Yavnella s. l*.
14. The mesoscutellum is posteriorly prolonged in *Leptanilla* TH01 and *Leptanilla* zhg-th01 (Fig. 6B). The differences in mesoscutellar shape between these morphospecies (see Appendix) are such that the homology of posterior mesoscutellar prolongation is uncertain.
15. The propodeum has a distinct planar to depressed dorsal face in the Bornean morphospecies-group (Fig. 7B). This condition is an autapomorphy of that clade.
16. The posterior margin of abdominal sternite IX is variously emarginate to entire in male Leptanillinae or with a posteromedian process (e.g., *Protanilla* zhg-vn01, *Yavnella* TH03), but posterolateral filiform processes of abdominal sternite IX are an autapomorphy of the Bornean morphospecies-group.
17. Abdominal tergite VIII is longer than broad only in *Noonilla* (Fig. 17B), *Scyphodon* and a bizarre male morphospecies from Côte d’Ivoire (CASENT0102373) for which molecular data are unavailable.
18. The gonocoxae exhibit partial (Fig. 8B) to full (Fig. 8C) medial fusion at least in ventral view in *Yavnella* TH03, *Noonilla*, and all sampled members of the Bornean morphospecies-group. Within *Leptanilla s. l.*, complete lack of medial gonocoxal fusion (Fig. 8A) is a symplesiomorphy of *Leptanilla s. str.*, *Leptanilla* TH01, and *Leptanilla* zhg-th01.
19. Articulation of the gonopodites encompasses both cases in which conjunctival membrane is visible between the gonocoxa and stylus, and those in which the stylus is recurved relative to the gonocoxa without apparent conjunctival membrane. This character state is a symplesiomorphy of *Leptanilla s. str.*, and among *Leptanilla s. l.* included in this study is observed in *Noonilla* zhg-my02 and −6, and *Leptanilla* zhg-th01.
20. The volsellae cannot be observed without dissection in many male Leptanillinae (e.g., *Noonilla*), limiting my ability to assess their condition. However, *Leptanilla s. str.* contrast with the Anomalomyrmini, *Yavnella s. l.*, and the Bornean morphospecies-group in that the volsellae (where visible) are dorsoventrally flattened, entire, and lacking sculpture (Fig. 18). This is one of only two synapomorphies of *Leptanilla s. str.* relative to other *Leptanilla s. l*.
21. Dorsoventral compression at the penial apex is also observed in *Yavnella s. l.* (except for *Yavnella* TH03). In the Indo-Malayan sister clade of *Leptanilla s. str.* the penial sclerites are lateromedially compressed to subcircular, at least basally. *Leptanilla* zhg-th01 exhibits an intermediate condition, with the penial apex being lateromedially compressed and this condition less pronounced towards the base.
22. Position of the phallotreme with distal margin adjoining the penial apex appears to be ancestral for the Leptanillini. The phallotreme is shifted basally in *Leptanilla* zhg-my02 and –5 (Fig. 19B), *Noonilla*, and *Scyphodon*. The outline of the phallotreme is subcircular in these morphotaxa. Setae surrounding the phallotreme are observed in *Noonilla* and *Scyphodon*; this character state is likely a synapomorphy of these genera.

**Fig. 15.**
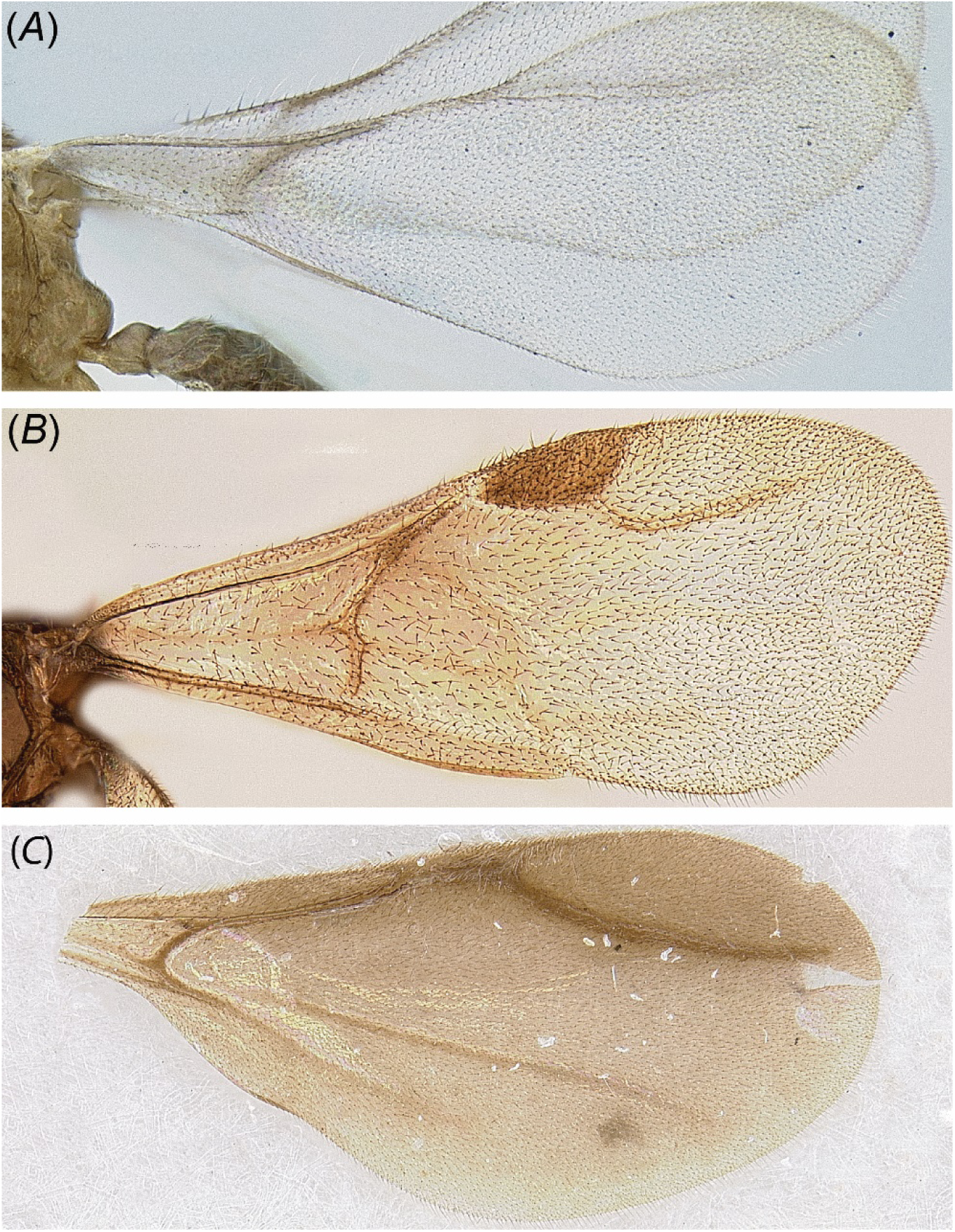
Examples of male forewing venation across the Leptanillinae. (A) *Yavnella* zhg-bt01 (CASENT0106384); (B) *Protanilla* zhg-vn01 (CASENT0842613); (C) *Leptanilla* (Bornean morphospecies-group) zhg-my05 (CASENT0842571).

**Fig. 16.**
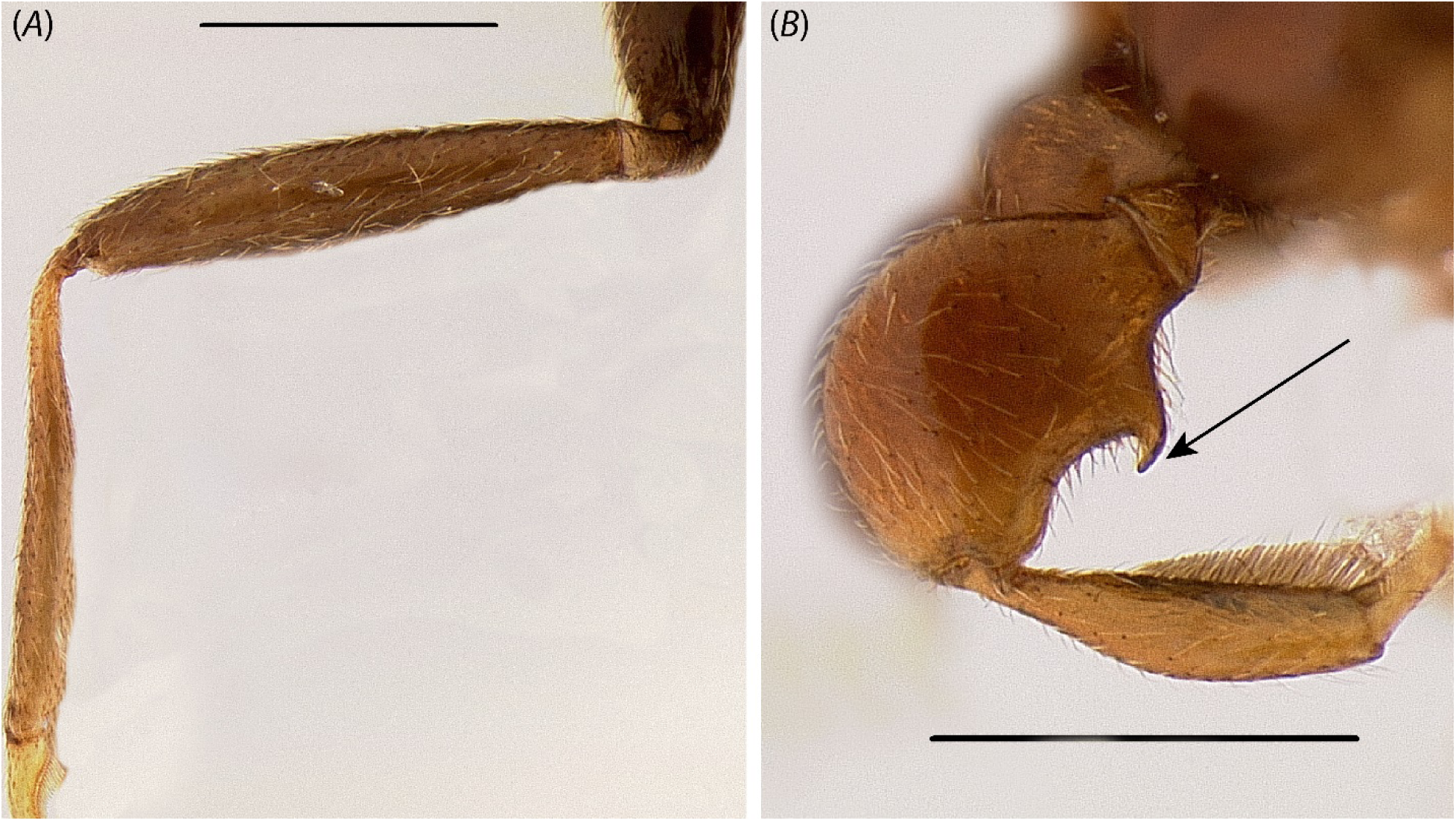
Foreleg of (A) *Yavnella argamani* (CASENT0235253) and (B) *Leptanilla* zhg-id01 (CASENT0842626; not sequenced in this study). Scale bar=0.3 mm.

**Fig. 17.**
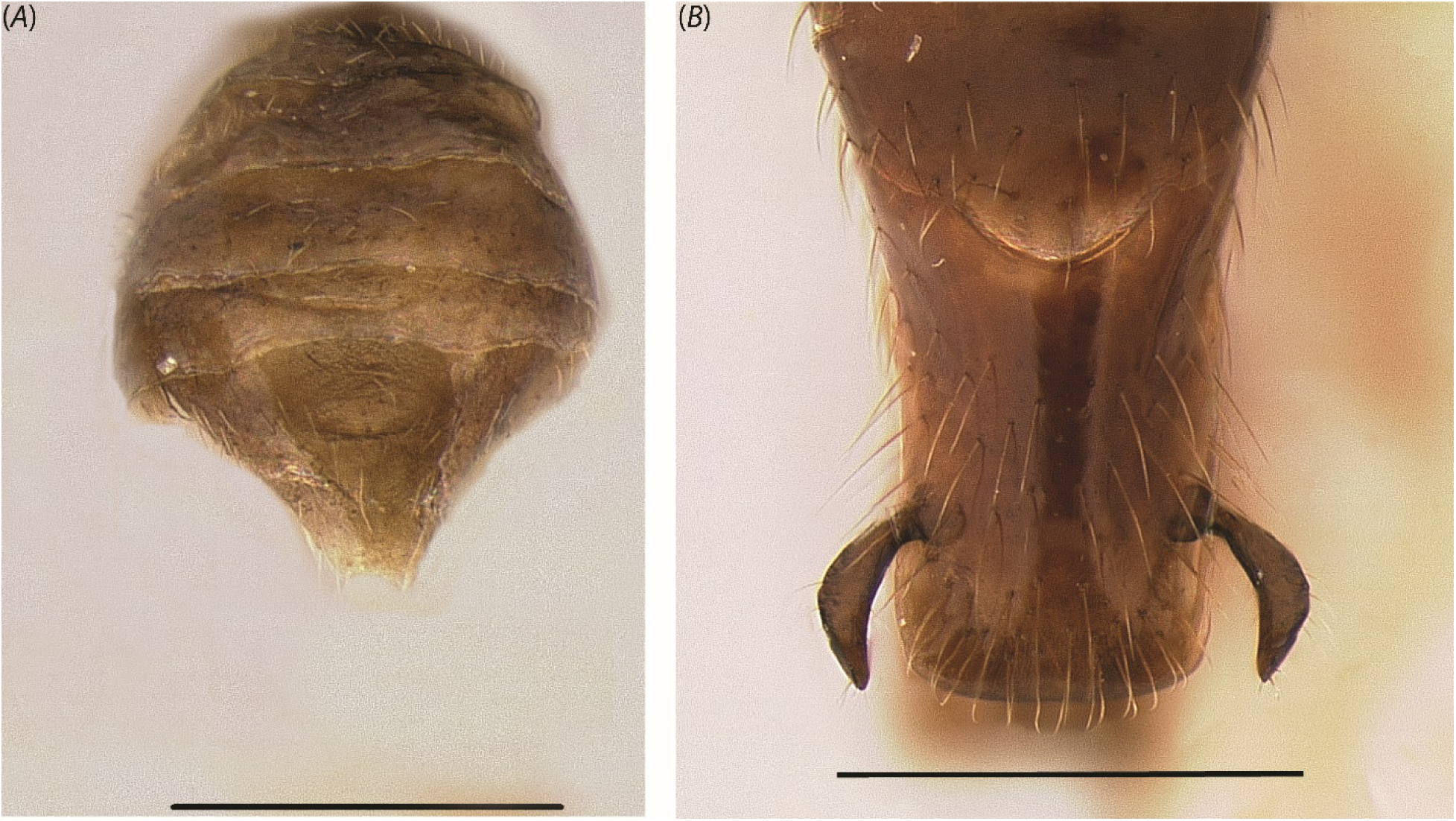
Posterior view of abdominal tergite VIII in male Leptanillini. (A) *Yavnella* zhg-th01 (CASENT0842620) and (B) *Noonilla* zhg-my02 (CASENT0842592). Scale bar = 0.3 mm.

**Fig. 18.**
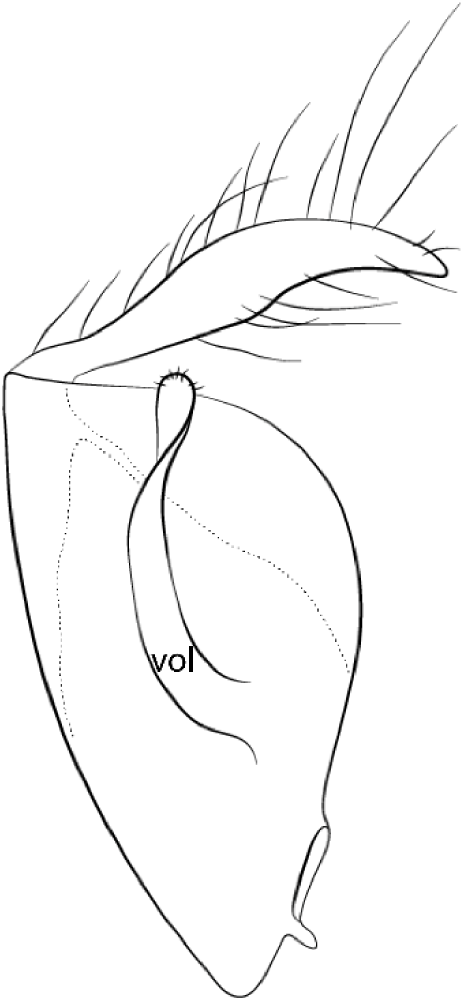
Gonopodite and volsella (*vol*) of *Leptanilla africana*, sketched after Baroni Urbani (1977: Fig. 37) by M. K. Lippey. Top of image is distal to body.

**Fig. 19.**
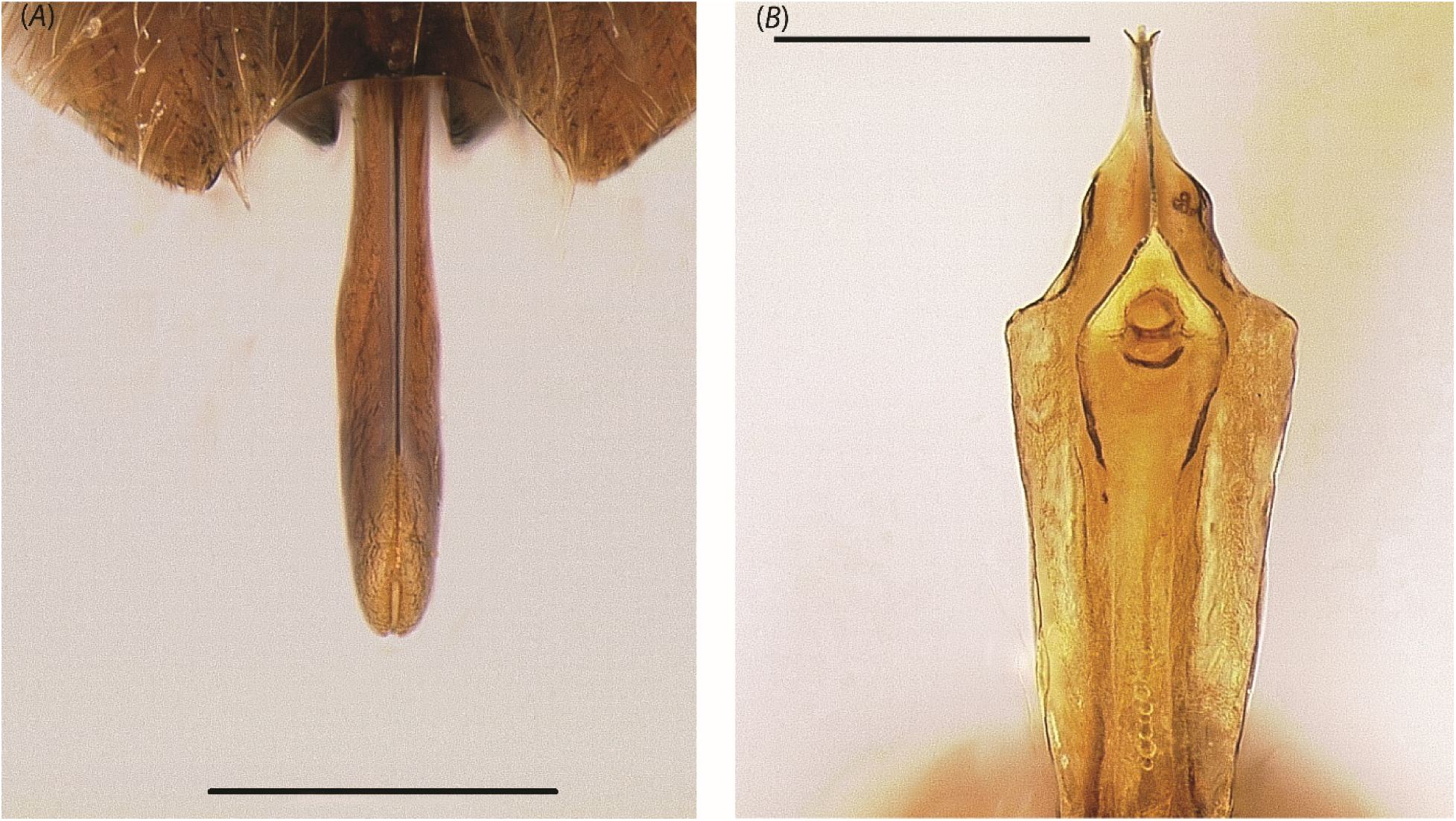
Dorsoposterior view of phallotreme in (A) *Leptanilla* zhg-my04 (CASENT0842553); ventroposterior view of phallotreme in (B) *Leptanilla* zhg-my05 (CASENT0106432). Scale bar A = 0.3 mm.; B = 0.4 mm.

### Goals of Future Research

Two described male-based species of *Leptanilla* are worth noting here as requiring further study and acquisition of fresh material: *L. palauensis*, which was transferred with some reservation to *Leptanilla* from *Probolomyrmex* (Formicidae: Proceratiinae) by Taylor (1965); and *Leptanilla astylina.* Examination of the holotype of *L. palauensis* demonstrates that according to the morphological hypotheses made herein this species can be confidently referred to *Leptanilla s. l.*, but beyond that its affinities are unclear. Based upon available illustrations (Petersen 1968: Fig. 1) *L. astylina* likewise can be placed in *Leptanilla s. l.*, and closely resembles *Leptanilla s. str.*, excluding its genitalia, which to judge from Petersen (1968) are unlike those of any specimen that was examined in this study, and exclude it from the definition of *Leptanilla s. str.* given herein.

The case of *Scyphodon* must also be briefly addressed here. Examination of a specimen attributable to this monotypic male-based genus shows that it can be placed in *Leptanilla s. l.* As reported by Petersen (1968), the genitalia of *Scyphodon* conspicuously resemble those of *Noonilla*: there is no reason to conclude that *Scyphodon* belongs within *Leptanilla s. str.,* and I predict that *Scyphodon* is either sister to, or nested within, *Noonilla*. Future total-evidence Bayesian phylogenetic inference will resolve the relation of *Scyphodon* to other *Leptanilla s. l*.

Future acquisition and examination of novel material may necessitate revision of the male diagnosis of *Leptanilla* provided here, but this diagnosis is robust to all morphological observations made with sequenced material. As *Yavnella s. l.*, *Noonilla* and the Bornean morphospecies-group are known only from males, and *L. revelierii* is known only from female castes, no argument can yet be made regarding the ranking of the former clades relative to *Leptanilla*. *Yavnella* is here ranked as a genus, but the description of *Yavnella* workers may reveal a morphological basis for subjective arguments for the subsumption of *Yavnella* within *Leptanilla*. The delimitation of genera within the Leptanillini—including the status of *Noonilla* and undescribed male morphospecies more closely related to that genus than to *L. revelierii*— therefore depends not only upon phylogenetic resolution of the many lineages known only from male material, but upon the morphology of corresponding workers. Future molecular sequencing will be needed to associate workers and/or gynes to leptanilline lineages that are known only from males: such an effort has successfully linked *Protanilla lini* (Anomalomyrmini) with previously unassociated males (Griebenow, in press).

## Conclusions

I have here demonstrated the utility of discrete morphological data within a total-evidence framework that includes molecular data in inferring the phylogeny of an ant taxon known only from male morphology. Using probabilistic models, the phylogenetic position of *P. javana* is robustly inferred in conjunction with taxa for which only molecular data, or both these and male morphological data, are available. In that phylogeny, *P. javana* and *L. revelierii* are confidently recovered within a subclade easily diagnosed by male morphological characters; disregarding future retrieval of worker material and/or novel male specimens, *Phaulomyrma* can be synonymized with *Leptanilla* despite continued uncertainty in the bounds of the latter genus. Future work will employ this Bayesian total-evidence approach to infer the affinity of other, more peculiar leptanilline taxa for which molecular data are unavailable. With a robust phylogeny inferred for the Leptanillinae that is congruent with male morphology, the parallel taxonomy that bedevils this little-understood group of ants can begin to be resolved.

## Acknowledgments

First and foremost, I must thank Ziad Khouri for his generosity in providing indispensable assistance and conceptual advice on the writing of scripts for the Bayesian phylogenetic analyses upon which this project relied. I thank Jadranka Rota (MZLU), Debbie Jennings (ANIC), Kevin Williams (CSCA), and Brian Fisher (CASC) for loans of material for examination and non-destructive DNA extraction; Stefan Cover was of great help in facilitating access to a syntype of *Leptanilla. javana*. I also thank my present and past lab-mates Jill Oberski and Matt Prebus for their diligent and painstaking work enriching UCEs from the specimens that were used in this study (sometimes with my help, sometimes not). In the realm of data collection, I am grateful to Eli Sarnat, Steve Heydon and Lynn Kimsey for allowing me the usage of equipment for this study; the aid of Michael Branstetter in providing advice on the retrieval of legacy loci from UCE datasets; Ziv Lieberman in advising me to use *BBMap*; and my labmate Brendon Boudinot, who was an enduring source of informative feedback on the coding of morphological observation into discrete character states. Lastly, I must thank my adviser Phil Ward for his invaluable feedback on the construction and finer details of this manuscript, along with past work in acquiring the Sanger-sequenced data that I utilized in this study, and for tutoring me in the delimitation of protein-coding loci from flanking introns. This research was supported by the University of California, Davis and by NSF grant DEB-1932405 to P. S. Ward.

## Appendix

Definition of binary presence/absence morphological characters. Note that all non-genital morphological data are missing in *Leptanilla* ZA01, since all that remained of this specimen after destructive DNA extraction was the male genitalia. Missing observations are noted for other terminals where relevant. Males of *Protanilla lini* Terayama, 2009 were identified as such by molecular data (Griebenow, in press).

1. Mesal protibial margins carinate: a sclerotized carina (Fig. 20A) is present (1) on the mesal margin of the ventral protibial surface in *Noonilla*. This character could not be scored in *Leptanilla* TH01. Under the alternative character state (0) the mesal protibial face is convex (Fig. 20B) to carinate.
2. Ventral cuticular hook present on profemur: the lateral margin of the ventral profemoral surface is ventrally produced into a hook-like structure (Figs. 16B, 21B) (1) in *Leptanilla* (Bornean morphospecies-group) zhg-my02 and -my05. The morphospecies imaged in Figs. 16B, 21B is closely related to these (Griebenow, in press) but was not sequenced in this study. Under the alternative character state (0) there are no cuticular extensions of the profemur (Fig. 16A). This character could not be scored in *Leptanilla* TH01.
3. Row of ventral protibial bristles present: a single medial row of parallel-sided setae is present (1) on the ventral protibial surface only in the “Bornean morphospecies-group” (Fig. 21B). These are robust by comparison to adjacent unmodified setae. Under the alternative character state (0) setae on the protibial venter are not robust, parallel-sided, and arranged in a single medial row (Fig. 21A). This character could not be scored in *Leptanilla* TH01.
4. Head inclusive of compound eyes wider than long: this character state is observed (1) in *O. hungvuong*; all male Anomalomyrmini sampled herein; all *Yavnella s. l.* except for *Yavnella* TH05, −8, and *Yavnella* MM01; *Leptanilla* (Bornean morphospecies-group) zhg-my04; and *Noonilla* zhg-my06 (Fig. 22B). Under the alternative character state (0) the head inclusive of the compound eyes is narrower than long in full-face view (Fig. 22A).
5. Clypeus broader than torular diameter along medial axis: this character state is observed (1) in *M. heureka*; *O. hungvuong*; in all Anomalomyrmini sampled herein; and in all *Yavnella s. l.* for which observations are available (Fig. 23B). Clypeus narrower than torular diameter along medial axis (0) (Fig. 23A) may therefore be diagnostic for *Leptanilla s. l.* This character could not be scored in *Yavnella* TH03, −5 and −8; *Yavnella* zhg-bt01; *Phaulomyrma* MM01; *Leptanilla* zhg-th01; and *Leptanilla* GR01-3, zhg-au02 and zhg-bt01.
6. Anterior tentorial pits situated directly anterior to toruli: the anterior tentorial pits are situated directly anterior to the toruli, in whole (Fig. 24B) or in part (1) so that at least some portion of the anterior tentorial pit intersects an anteroposterior axis drawn through the torulus, in *M. heureka*, *O. hungvuong*, all Anomalomyrmini save *Protanilla* TH01, and all *Yavnella s. l.* save *Yavnella* TH05 and MM01. Under the alternative character state (0), the anterior tentorial pits are situated anterolaterad the toruli or may not be readily discernible (Fig. 24A), so that no part of the anterior tentorial pit intersects an anteroposterior axis drawn through the torulus. This character could not be scored in *Yavnella* TH03 and −8, *Leptanilla* (Bornean morphospecies-group) zhg-my02, *Leptanilla* zhg-au02, and *Leptanilla javana*.
7. Antennomere 3 longer than scape: this character state (1) (Fig. 25B) is observed in *Protanilla* TH03 and all *Yavnella s. l.* except for *Yavnella* TH05. Under the alternative character state (0) the scape is shorter than (Fig. 25A) or subequal in length to antennomere 3. This character could not be scored in *O. hungvuong* or *Leptanilla* zhg-au02.
8. Mandible articulated to gena: the base of the male mandible is visibly fused to the gena (0) in all *Yavnella s. l.* for which observations are available (Fig. 12A), except for Yavnella TH04. In all other terminals in which this character can be assessed a complete point of articulation to the gena is visible (1) (Fig. 12B). This character could not be scored in *Yavnella* TH03 and MM01; and *Leptanilla* zhg-au02, -TH09 and *javana*.
9. Occipital margin angularly emarginate in dorsal view: the occiput is coded as angularly emarginate in dorsal view (1) if the posterolateral corners of the occipital margin are produced; this character state is observed in *Leptanilla* TH01 (Fig. 26B), *Leptanilla* zhg-th01, and the Bornean morphospecies-group except for *Leptanilla* zhg-my03. Under the alternative character state (0) the occiput is linear to shallowly emarginate (Fig. 26A).
10. Mesoscutum convex in profile view: the mesoscutum is scored as convex (1) if not planar to shallowly convex (0) (Fig. 27A). Mesoscutal convexity (1) (Fig. 27B) is present in *M. heureka*, *O. hungvuong*, the Anomalomyrmini, *Yavnella s. l.*, and *Leptanilla* (Bornean morphospecies-group) zhg-my04.
11. Notauli present: the presence (1) or absence (0) (Fig. 28A) of notauli is always unambiguous. These are observed only in *M. heureka*, *Protanilla* TH01, and −03 (Fig. 28B).
12. Parapsidal signa present: the presence (1) (Fig. 29B) or absence (0) (Fig. 29A) of the parapsidal signa can be difficult to discern, varying from a distinct impressed signum to a stripe of glabrous cuticle. Some form of parapsidal signum is present in *M. heureka*; *O. hungvuong*; *Protanilla* zhg-vn01; *Yavnella* zhg-th01, cf. *indica* and *argamani*, *Yavnella* TH02, −4 and −6; *Yavnella* MM01; *Yavnella* TH01; *Noonilla* zhg-my06; the Bornean morphospecies-group; and *Leptanilla* GR01.
13. Oblique mesopleural sulcus adjoining posterior mesopectal margin: this character state is observed (1) in *O. hungvuong*, all Anomalomyrmini (Fig. 30B) and most *Leptanilla s. str.* for which this character can be scored, except for *Leptanilla* GR01, −03, TH09, and *Leptanilla* zhg-bt01. Complete bisection of the mesopectus by the oblique mesopleural sulcus is seen in the Anomalomyrmini. The alternative character state (0) encompasses a morphocline from the near-complete loss of the oblique mesopleural sulcus (as in *Leptanilla* zhg-bt01) to the termination of this feature immediately anterior to the upper metapleuron (e.g., *Yavnella* TH02: Fig. 30A) or propodeum (as in the Bornean morphospecies-group). This character could not be scored in *Leptanilla* zhg-au02.
14. Pterostigma present: this character state is observed (1) only in *M. heureka*, *O. hungvuong*, and the Anomalomyrmini (Fig. 31B). Rf and 2s-rs+Rs+4-6 are confluent in the Bornean morphospecies-group and in *Noonilla* zhg-my06, producing an infuscation of the wing membrane that resembles a pterostigma (0). No infuscation or pterostigma (0) is observed in all other terminals scored (Fig. 31A). Wings are lost in all available specimens of *Noonilla* zhg-my02, *Leptanilla* zhg-th01, and *Leptanilla* GR03; therefore, this character could not be scored in these terminals.
15. Mesoscutellum densely pubescent: the mesoscutellum is covered with sparse setae (0) in all leptanilline males sampled herein except for *Leptanilla* TH01 and zhg-th01, and the Bornean morphospecies-group (Fig. 32B); in these cases, the mesoscutellar vestiture is densely pubescent (1) (Fig. 32A). This character could not be scored in *Yavnella* TH04.
16. Mesoscutellum projecting posteriorly in profile view: this character state is observed (1) either as a dorsoventrally robust cuneiform process (*Leptanilla* TH01) or as a recurved spine (*Leptanilla* zhg-th01) (Fig. 6B). Under the alternative character state, the posterior margin of the mesoscutellum is rounded (0) (Fig. 6A). This character could not be scored in *Yavnella* TH02.
17. Propodeum concave in profile view: this character state (1) (Fig. 7A) is an autapomorphy of *Yavnella s. l.* Under the alternative character state (0) the propodeum is convex in profile view (Fig. 7C) or produced into a right angle, with largely planar dorsal and posterior faces (the Bornean morphospecies-group; Fig. 7B).
18. Abdominal tergite II produced into distinct node: there is a shallow to pronounced dorsal node (Fig. 33B) present on the petiole (1) in *O. hungvuong*, *Protanilla* zhg-vn01 and TH01-2, *Yavnella* TH08, *Leptanilla* zhg-th01, the Bornean morphospecies-group, and *Leptanilla s. str.* except for *Leptanilla* zhg-au02. Under the alternative character state (0) the dorsal surface of the petiole is slightly convex (Fig. 33A), or planar without any supra-axial projection (as in *Leptanilla* zhg-au02).
19. Abdominal sternite II with ventral process: a ventral rounded to angular process (1), shallow or well-produced, is present on abdominal sternite II in *Protanilla* zhg-vn01 and TH02, *Leptanilla* zhg-my02 (Fig. 34C) and −5, and *Leptanilla s. str.* except for *Leptanilla* zhg-au02, and *javana*. Under the alternative character state (0) there is no ventrally projecting process on abdominal sternite II (Fig. 34A). A moderate ventral bulge without a distinct anterior and/or posterior face may be present under this character state (Fig. 34B). This character could not be scored in *Protanilla* TH01.
20. Petiole higher than long including peduncle: this character state (Fig. 35B) is observed in profile view (1) in *Protanilla* zhg-vn01 and TH01-2, *Yavnella* MM01, *Yavnella* TH05, *Yavnella* cf. *indica* and zhg-th01, *Leptanilla* TH01, the Bornean morphospecies-group, and *Noonilla*. This includes cases in which there is no distinct dorsal node. Under the alternative character state (0) the distance between two lines drawn tangential to the dorsal- and ventral-most points of the petiole in profile view is no greater than petiole length in profile view (Fig. 35A). This character could not be scored in *Yavnella* TH02.
21. Cinctus present on abdominal segment III: the corollary of this character state (1) is the existence of a petiole (Fig. 36B), which has been secondarily lost (0) in *Yavnella* zhg-th01 (Fig. 36A), *Yavnella* TH02 (as noted by Boudinot 2015: p. 14), and *Noonilla* zhg-my02. There is a tendency towards petiolar reduction in *Yavnella s. l.* and *Noonilla*, but in many cases a cinctus on abdominal segment III is still discernible.
22. Cinctus present on abdominal segment IV: the corollary of this character state (1) is the presence of a post-petiole. This character state is unique to *Protanilla* TH03 (Fig. 37B), although the anterior margin of abdominal segment IV may be slightly constricted relative to more posterior abdominal segments (0); otherwise, there is no constriction whatsoever (Fig. 37A).
23. Abdominal sternite IX with posteromedian filiform process: while a posteromedian process of abdominal sternite IX is present in all male Anomalomyrmini and *Opamyrma hungvuong* (0), its filiform condition (1) is unique to *Yavnella* TH03. Abdominal sternite IX is not thus produced medially in all other male leptanillines sampled herein (0).
24. Abdominal sternite IX with posterolateral filiform processes: these “bizarre, elongate, filamentous extensions” of the metasoma were noted by Boudinot (2015: Fig. 10D) as being extensions of the gonocoxae *sensu* Boudinot (2018). Detailed examination and micro-CT segmentation (Griebenow, Fischer and Economo in prep.) demonstrate that these processes are in fact extensions of abdominal sternite IX (Fig. 38B). This character state is unique to the Bornean morphospecies-group. Under the alternative character state (0) the posterior margin of abdominal sternite IX may be medially indented (Fig. 38A), entire, or with a posteromedian process, as noted above. This character could not be scored in *Leptanilla* zhg-au02.
25. Abdominal tergite XIII broader than long: this character state is observed (1) in all male Leptanillinae scored (Fig. 17A) except for *Noonilla*, to which elongation of abdominal tergite XIII (0) is unique (Fig. 17B). This character could not be scored in *Yavnella* MM01 and *L. javana*.
26. Gonocoxae ventromedially fused along entire length: this character state (Fig. 8C) is observed in *O. hungvuong*, *Yavnella* TH03, and in all terminals within the Bornean morphospecies-group that could be scored (1). The alternative character state (0) encompasses partial (Fig. 8B) to complete (Fig. 8A) ventromedian fusion of the gonocoxae. This character could not be scored in *Protanilla* TH01, *Yavnella* MM01, *Leptanilla* TH01, *Leptanilla* (Bornean morphospecies-group) zhg-my05, and *L. javana*.
27. Gonocoxae dorsomedially fused along entire length: this character state is observed (1) in *hungvuong*, *Yavnella* TH03 and the Bornean morphospecies-group (Fig. 39B). Under the alternative character state (0) the gonocoxae are fully (Fig. 39A) to partly separate medially. This character could not be scored without dissection in *Noonilla* (in which abdominal tergite XIII conceals the gonocoxal dorsum) or in *Leptanilla* zhg-au02, -bt01, ZA01, and *javana*.
28. Gonocoxa with ventral lamina: a ventral laminate margin, variably produced and shaped, is present (1) on the gonocoxa (Fig. 40B), or on the basal part of the gonopodite in those cases in which the gonocoxa and stylus are insensibly fused, in *Yavnella* cf. *indica*; *Leptanilla* zhg-th01; *Leptanilla* (Bornean morphospecies-group) zhg-my02 and −5; and *Leptanilla* TH09, GR01-2, ZA01 (Fig. 40B), and zhg-au02. Under the alternative character state (0) no lamina is discernible whatsoever on the gonocoxa (Fig. 40A). I do not assert primary homology *sensu* de Pinna (1991) of laminate portions of the gonopodites in *Leptanilla* zhg-my02 and −5 with styli, since this does not meet the criterion of conjunction (Patterson 1982; de Pinna 1991) in CASENT0178838, a heterospecific member of the Bornean morphospecies-group (misattributed to *Protanilla* by Boudinot [2015]). This character could not be scored in *Protanilla* TH01 or in *Leptanilla* zhg-bt01 and *javana*.
29. Stylus articulated to gonocoxa: this character state (1) includes cases in which the stylus is sharply deflexed relative to the gonocoxa (Fig. 41B) or a conjunctiva is visible between the gonopodital sclerites. Under the alternative character state (0) a suture might be visible (as in many *Yavnella s. l.*) or the gonocoxa and stylus insensibly fused (as in the Bornean morphospecies-group; Fig. 41A). Gonopodital articulation is fully present in *O. hungvuong*, *Protanilla* zhg-vn01, *Yavnella* zhg-bt01, *Leptanilla* zhg-th01, all *Leptanilla s. str.* for which this character can be scored and both *Noonilla* included in this study. This character could not be scored in *Leptanilla* zhg-bt01.
30. Gonopodital apex with vestiture: this character and the next are so termed in order to encompass cases in which the stylus is insensibly fused to the gonocoxa (figs. 41A, 42A-B). The only terminals sampled here in which setal vestiture is not present on the gonopodital apex (0) are the Bornean morphospecies-group (Fig. 42A) except for *Leptanilla* zhg-my04. Otherwise (1) there are at least some setae present on the gonopodital apex (Fig. 42B). This character could not be scored in *Leptanilla* zhg-bt01.
31. Gonopodital apex bifurcated: this character state is observed (1) only in *Yavnella* TH08 (Fig. 43B), *Leptanilla* ZA01 and GR02. Under the alternative character state (0) the stylus may be entire (Fig. 43A) or may have a subapical tooth. This character could not be scored in *Leptanilla* zhg-bt01.
32. Penial sclerites enclosed dorsally by gonopodites at base: in this character state (1) the gonopodites may completely enclose (Fig. 44C) or partially overlap with (Fig. 44B) the penial sclerites. This character state is observed in *M. heureka*, *Yavnella* TH03, *Yavnella* zhg-th01 and zhg-bt01, the Bornean morphospecies-group, and *Leptanilla* zhg-au02. Under the alternative character state (Fig. 44A) (0), the penial sclerites are never dorsally surmounted by any portion of the gonopodites. This character could not be scored in *Noonilla* zhg-my06 and *L. javana*.
33. Penial sclerites dorsally recurved at base in profile view: among the terminals sampled here, this bizarre character state (1) is only present in *Leptanilla* (Bornean morphospecies-group) zhg-my02 and −5 (Fig. 45B). In these cases the penial sclerites are curved at the base so that in preserved specimens the apex is situated dorsally of the gonocoxae. Otherwise (0) in profile view the penial sclerites are slightly curved at the base towards the venter of the genital anteroposterior axis (Fig. 45A) or are parallel to that axis.
34. Penial sclerites dorsoventrally compressed at base: this character state is observed (1) in *M. heureka*, and all *Yavnella s. l.* (Fig. 46B) and *Leptanilla s. str.* for which this character can be scored. Under the alternative character state (0) the penial sclerites are basally wider along the dorsoventral axis, exclusive of any ventromedian processes, than along the lateromedial axis (Fig. 46A). This character could not be scored in *O. hungvuong*, *Protanilla* TH02-3, *Yavnella* TH03-4 and zhg-bt01, *Leptanilla* TH01, *Noonilla* zhg-my06, and *L. javana*.
35. Penial sclerites dorsoventrally compressed at apex: this character state is observed (1) in *M. heureka*, *Yavnella s. l.* except for *Yavnella* TH03, *Leptanilla s. str.*, and *Leptanilla* zhg-my03 (Fig. 47B). Under the alternative character state (0) the penial sclerites are apically wider along the dorsoventral axis, exclusive of any ventromedian processes, than along the lateromedial axis (Fig. 47A). The alternative character state (0) encompasses cases in which the penial sclerites are lateromedially compressed to varying extents (e.g., Anomalomyrmini) or are subcircular in cross-section (e.g., *Noonilla*). This character could not be scored in *O. hungvuong* or *Protanilla* TH02.
36. Lateral margins of penial sclerites laminate: this character state is observed (1) in *Yavnella* TH02-5, *Yavnella* MM01, *Yavnella* zhg-th01, cf. *indica* and *argamani*; *Leptanilla* zhg-my02 (Bornean morphospecies-group) and *Leptanilla* zhg-my05; and all *Leptanilla s. str.* (Fig. 48B) for which this character can be scored, except for *Leptanilla* zhg-bt01. In the Bornean morphospecies-group the lateral laminae, when present, are strongly produced ventrally relative to the remainder of the penial sclerites. Under the alternative character state (0) (Fig. 48A) lateral flanges may be present or absent, but when present are not laminate. This character could not be scored in *M. heureka* and *Leptanilla* zhg-au02.
37. Penial sclerites with dorsomedian carina: this character state is observed (1) only in *Leptanilla* TH01 and *Leptanilla* (Bornean morphospecies-group) zhg-my04 (Fig. 47A). In both cases the penial sclerites are strongly lateromedially compressed. Under the alternative character state (0) there is no dorsomedian penial carina, such that the dorsum of the penial sclerite(s) is/are rounded in cross-section (Fig. 47B). This character could not be scored in *Protanilla* TH03.
38. Penial sclerites with ventromedian projection: this character state is observed (1) in *Leptanilla* zhg-my03-4; *Leptanilla* zhg-th01; and *Leptanilla* zhg-bt01, GR01-2, and zhg-au02. When present and discernible, the volsellae flank this projection, which can be rounded (as in *Leptanilla* zhg-my03-4; Fig. 49B) or carinate. Under the alternative character state (0) the penial sclerites are entirely separated, or if fused then lacking any ventromedian process (Fig. 49A). This character could not be scored in *M. heureka*, the Anomalomyrmini, *Leptanilla* TH01 and *Yavnella* TH03-8 and *Yavnella* MM01.
39. Phallotreme flanked with vestiture: this character state (1) occurs only in *Noonilla* (Fig. 50B). Under the alternative character state (0) the phallotrematic rim is visibly bare of any setae (Fig. 50A). This character could not be scored in *Yavnella* MM01.
40. Phallotreme preapical: under the alternative character state (0) the phallotreme is situated adjoining the posterior penial margin or, if the penial sclerites are lateromedially compressed, at the penial apex (Fig. 19A). This includes cases in which the phallotreme is situated well basal to the penial apex but has a distal margin that extends to the penial apex. The phallotreme is therefore preapical (1) in *Leptanilla* (Bornean morphospecies-group) zhg-my02-3 and –5 (Fig. 19B), and in *Noonilla* zhg-my06. This character could not be scored in *Protanilla* TH02-3; *Yavnella* TH04, −8 and *Yavnella* zhg-bt01; and *Yavnella* MM01, and *Leptanilla* zhg-bt01 and *L. javana*.
41. Penial apex entire: the alternative (0) to this character state encompasses cases in which the penial sclerites are medially separated at the apex (as in *Protanilla* TH01-2), strongly bifurcated (Fig. 51A). Under this character state (1) none of these observations apply (Fig. 51B), encompassing cases in which the distal phallotrematic margin forms a narrow slit-like indentation in the penial sclerites (e.g., *Yavnella* cf. *indica*: Fig. 44A). The penial apex is entire in *M. heureka*; *Protanilla* TH03 and zhg-vn01; *Yavnella* TH02, −5-8, *Yavnella* cf. *indica*, zhg-bt01, and zhg-th01; *Leptanilla* TH01; *Leptanilla* zhg-th01; the Bornean morphospecies-group; and *Leptanilla s. str.* except for *Leptanilla* ZA01.

**Fig. 20.**
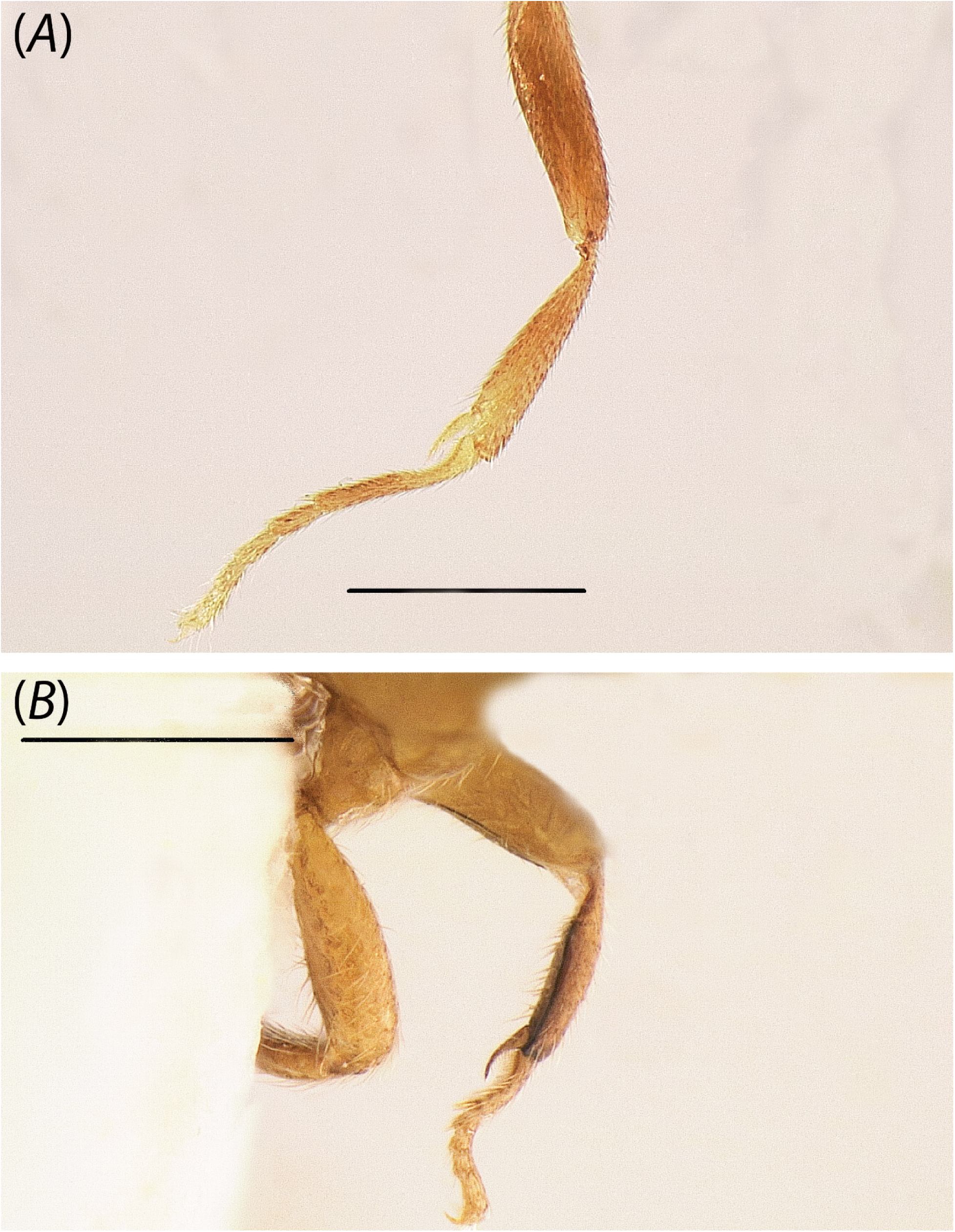
Mesal view of protibia of (A) *Protanilla* zhg-vn01 (CASENT0106382) and (B) *Noonilla* zhg-my01 (CASENT0842587; not sequenced in this study). Scale bar = 0.3 mm.

**Fig. 21.**
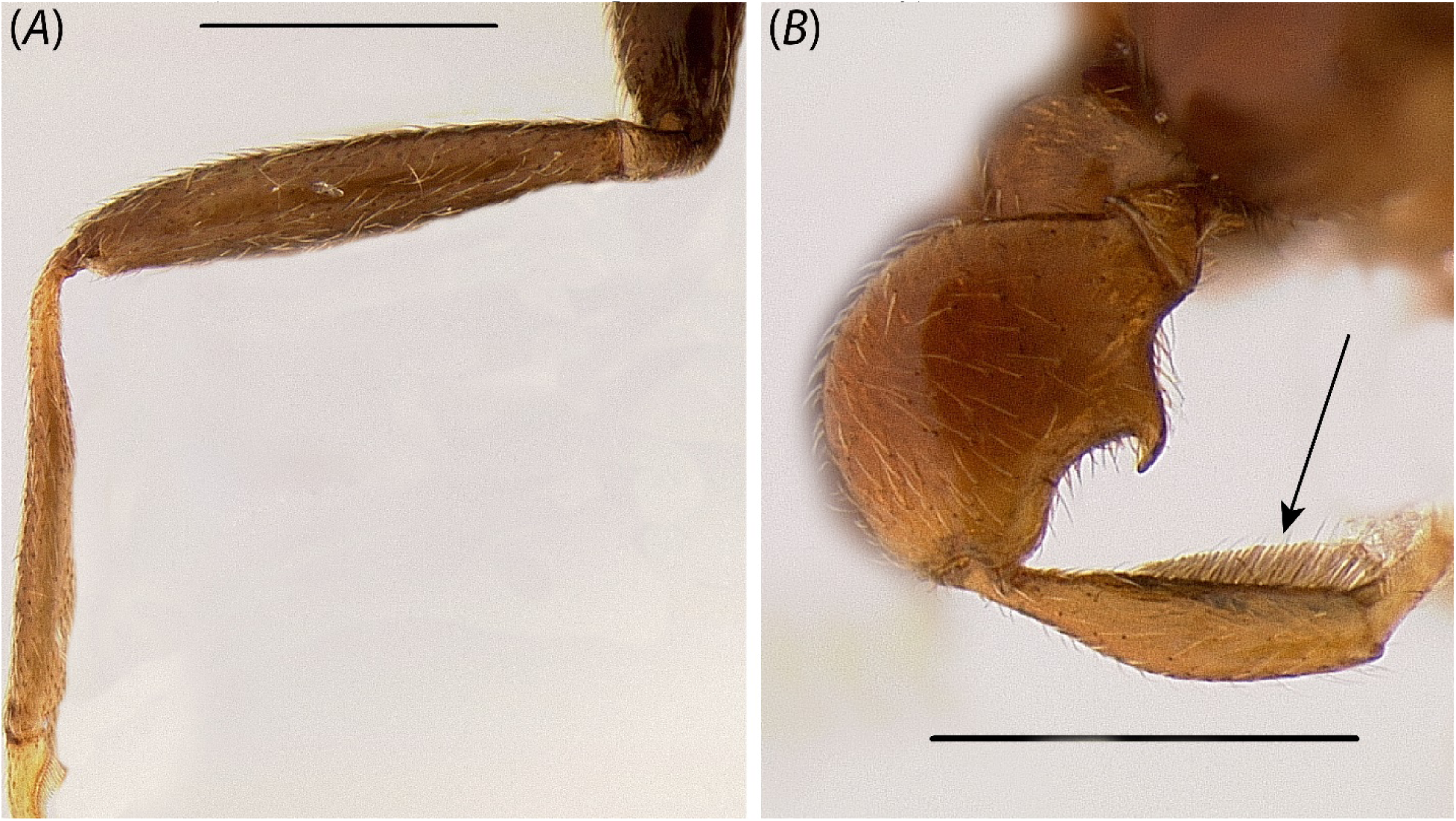
Foreleg of (A) *Yavnella argamani* (CASENT0235253) and (B) *Leptanilla* zhg-id01 (CASENT0842626; not sequenced in this study). Scale bar = 0.3 mm.

**Fig. 22.**
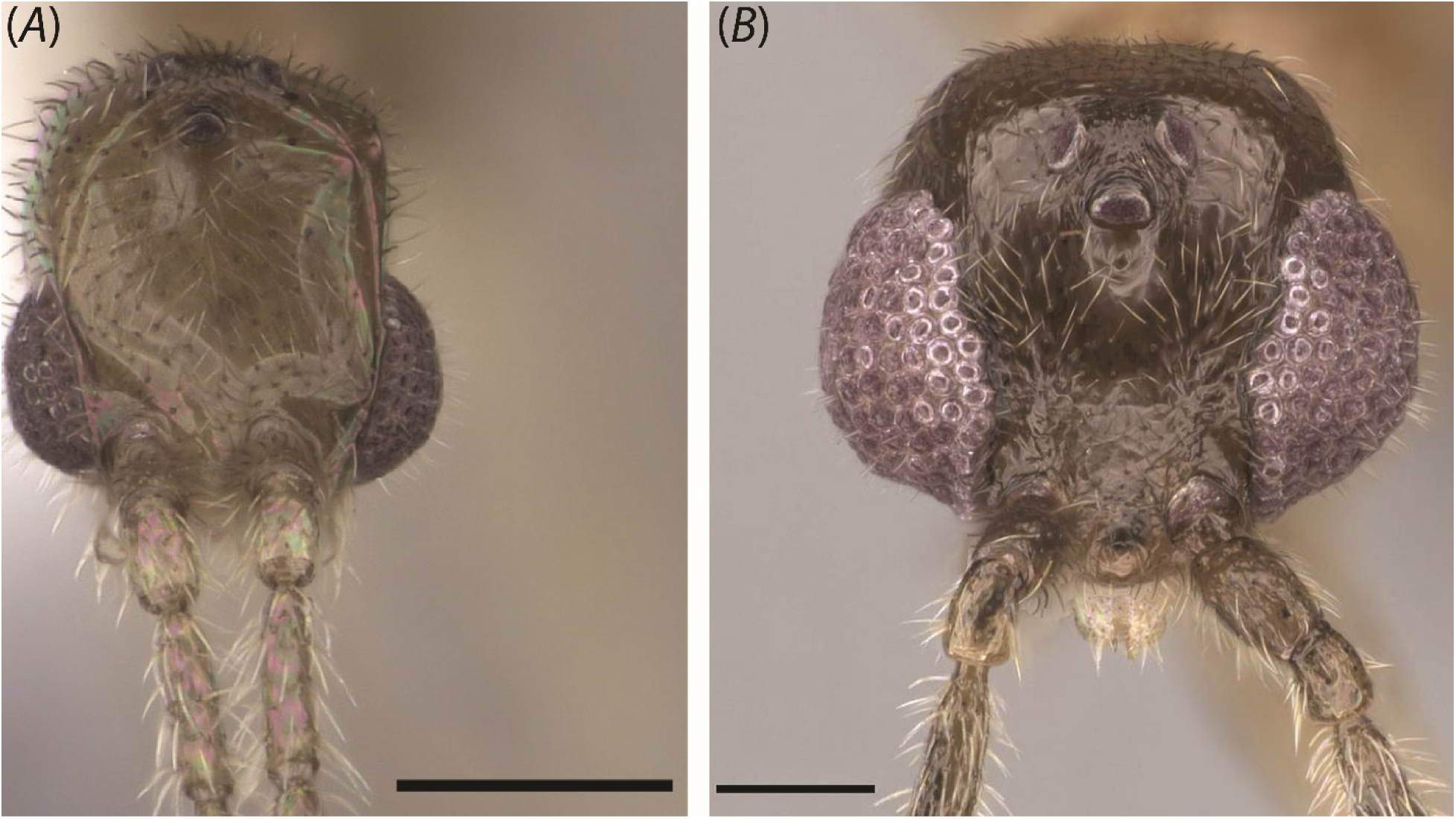
Full-face views of (A) *Yavnella* TH08 (CASENT0119531; Michele Esposito) and (B) *Yavnella* TH02 (CASENT0227555; Shannon Hartman). Scale bar = 0.1 mm.

**Fig. 23.**
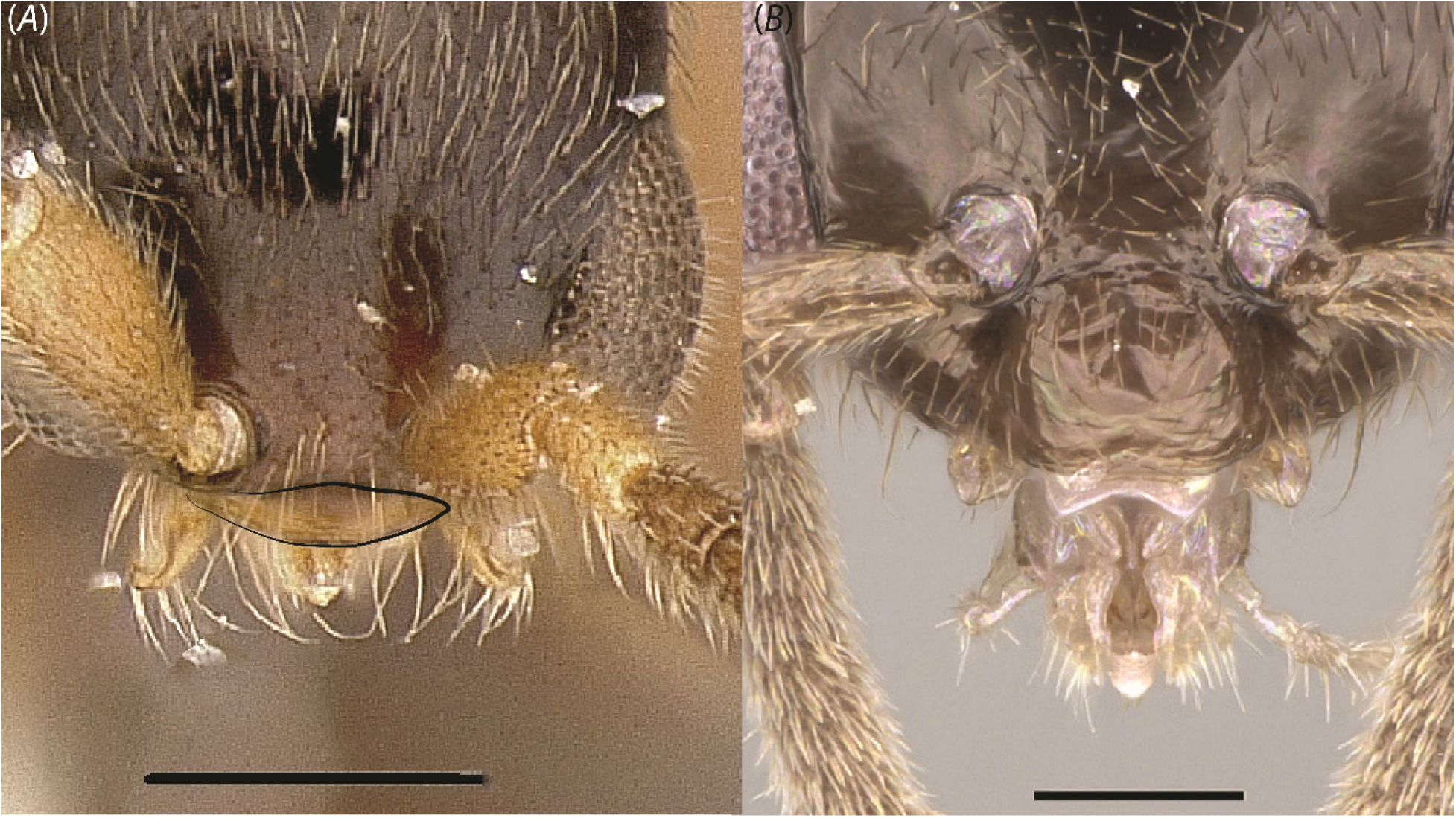
Full-face views of (A) *Leptanilla* zhg-my04 (CASENT0842558) and (B) *Protanilla* TH01 (CASENT0119776; Michele Esposito). Scale bar A = 0.1 mm.; B = 0.2 mm.

**Fig. 24.**
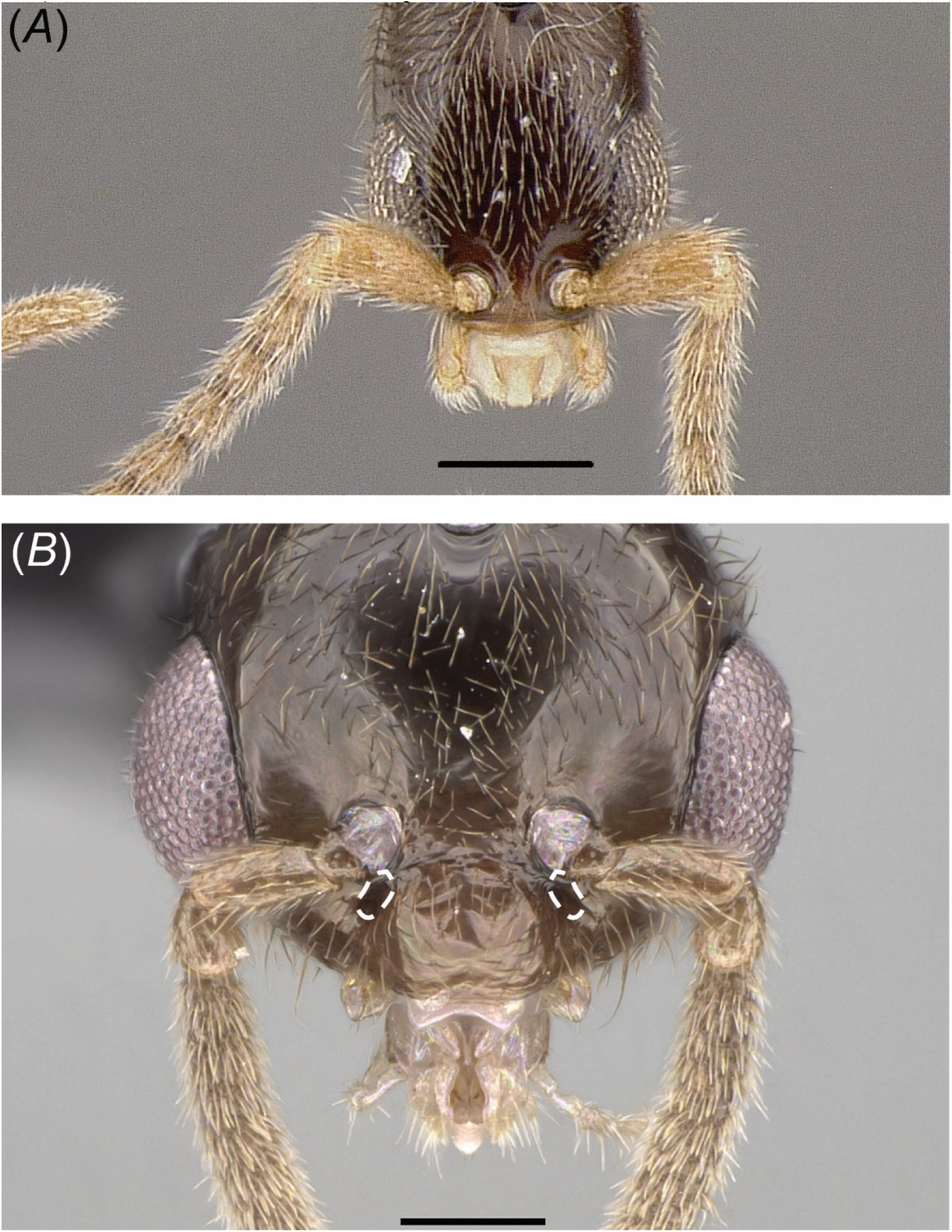
Full-face views of (A) *Leptanilla* zhg-my03 (CASENT084545) and (B) *Protanilla* TH01 (CASENT0119776; Michele Esposito). Scale bar A = 0.1 mm.; B = 0.2 mm.

**Fig. 25.**
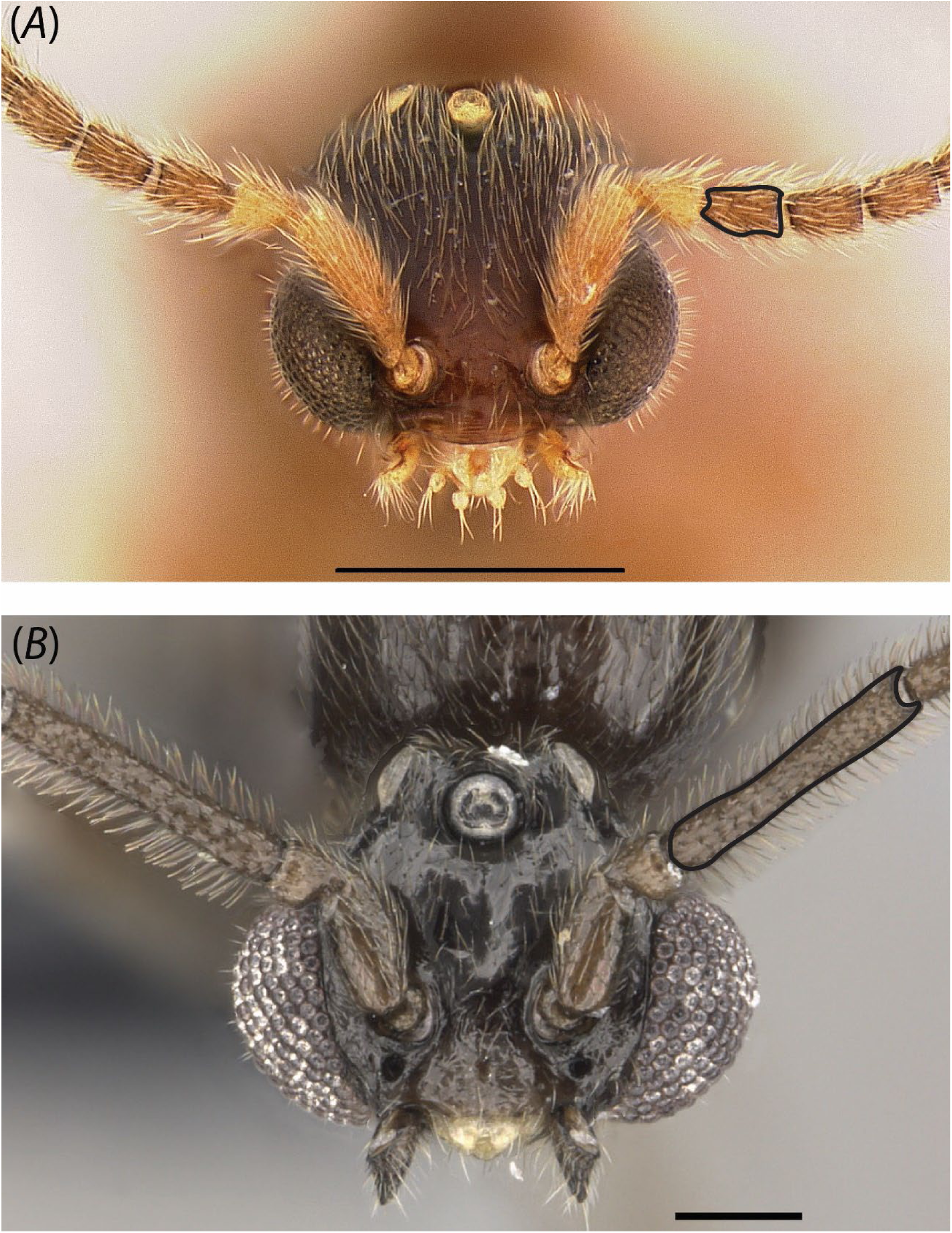
Full-face views of (A) *Leptanilla* zhg-my04 (CASENT0842548) and (B) *Yavnella argamani* (CASENT0235253; Shannon Hartman). Scale bar A = 0.3 mm.; B = 0.1 mm.

**Fig. 26.**
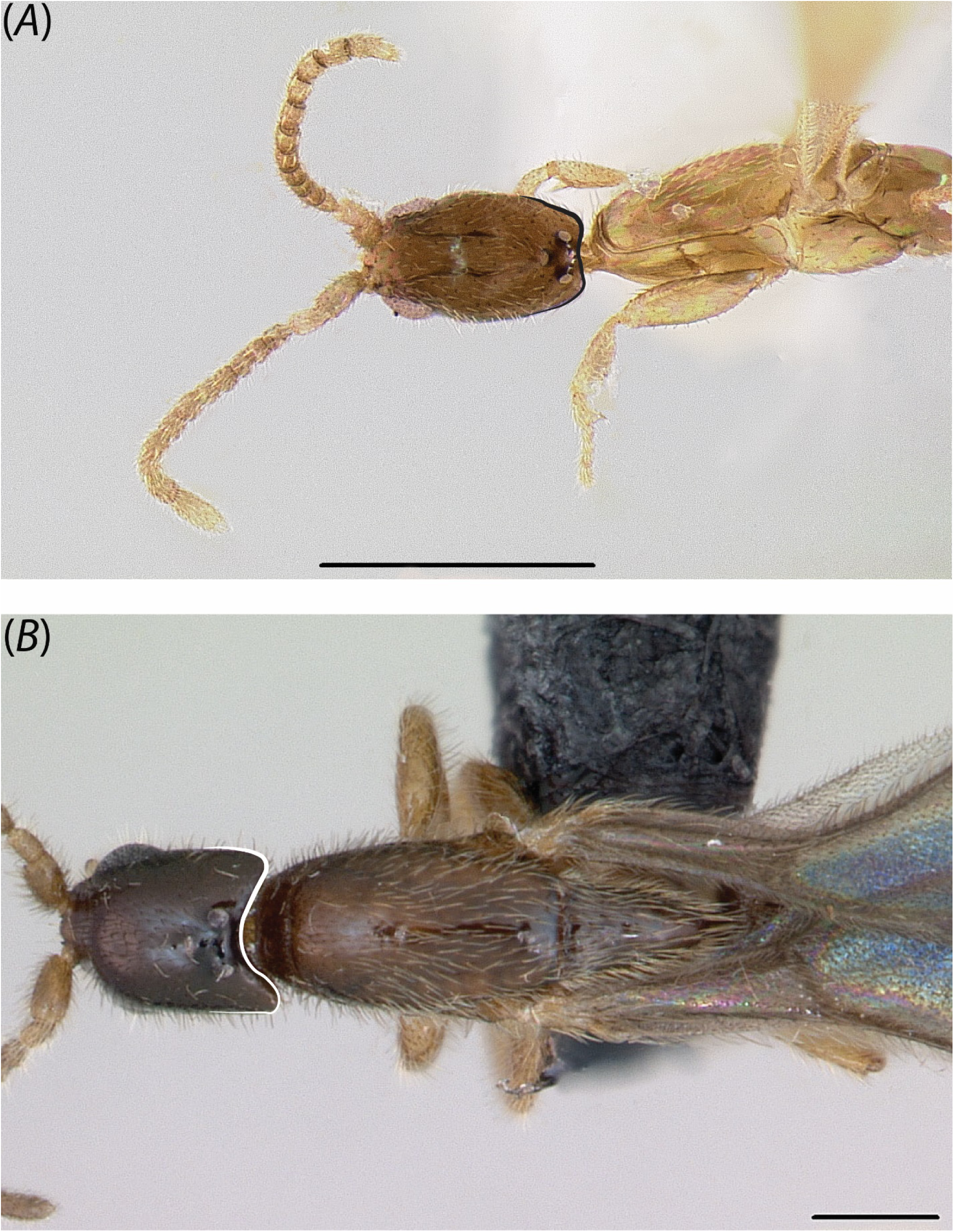
Dorsal view of occipital margin in (A) *Leptanilla* TH09 (CASENT0842664) and (B) *Leptanilla* TH01 (CASENT0119792; April Nobile). Scale bar A = 0.3 mm.; B = 0.2 mm.

**Fig. 27.**
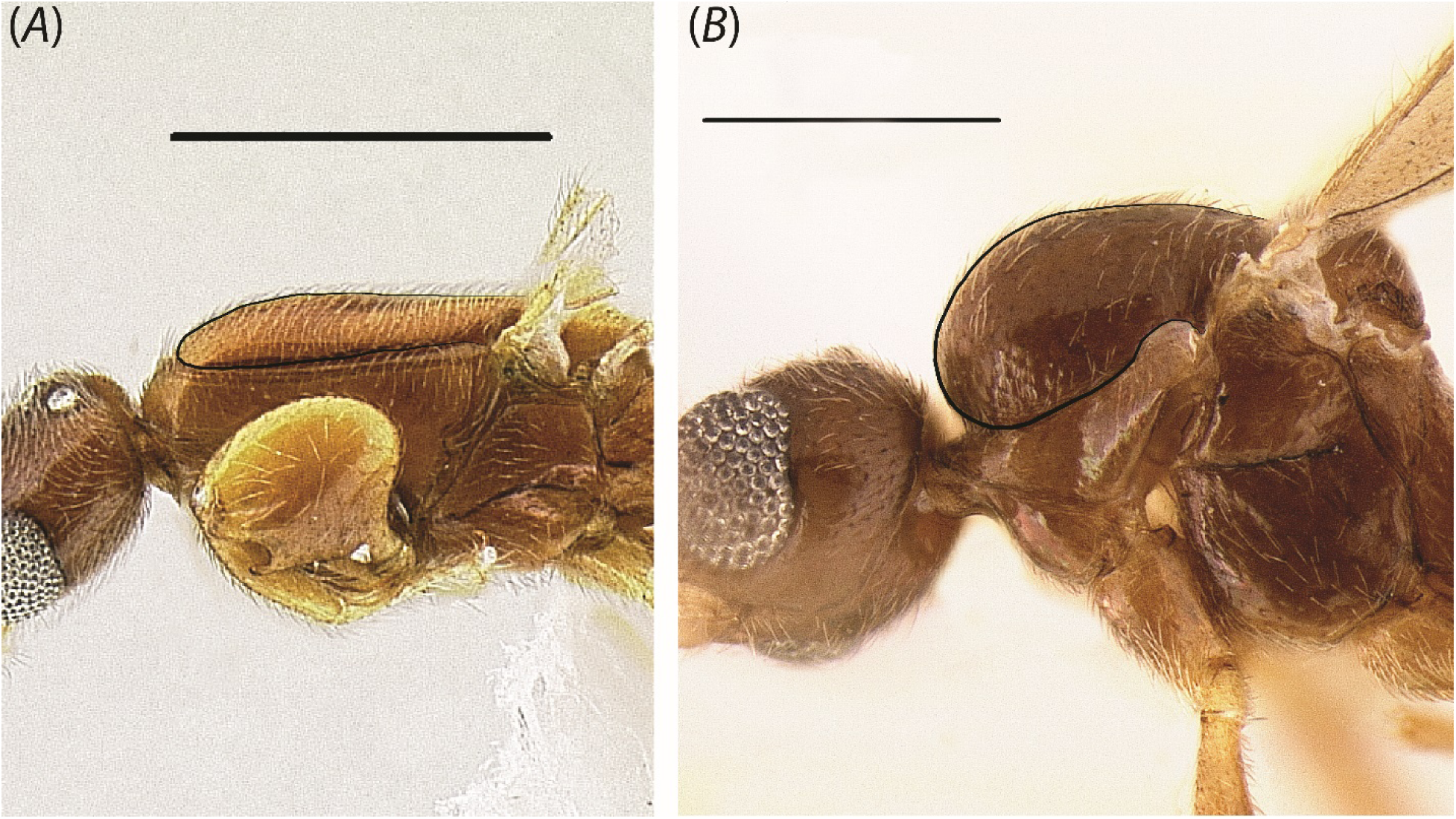
Profile view of mesosoma in (A) *Leptanilla* zhg-my02 (CASENT0106416) and (B) *Yavnella* zhg-th01 (CASENT0842621). Scale bar A = 0.5 mm.; B = 0.3 mm.

**Fig. 28.**
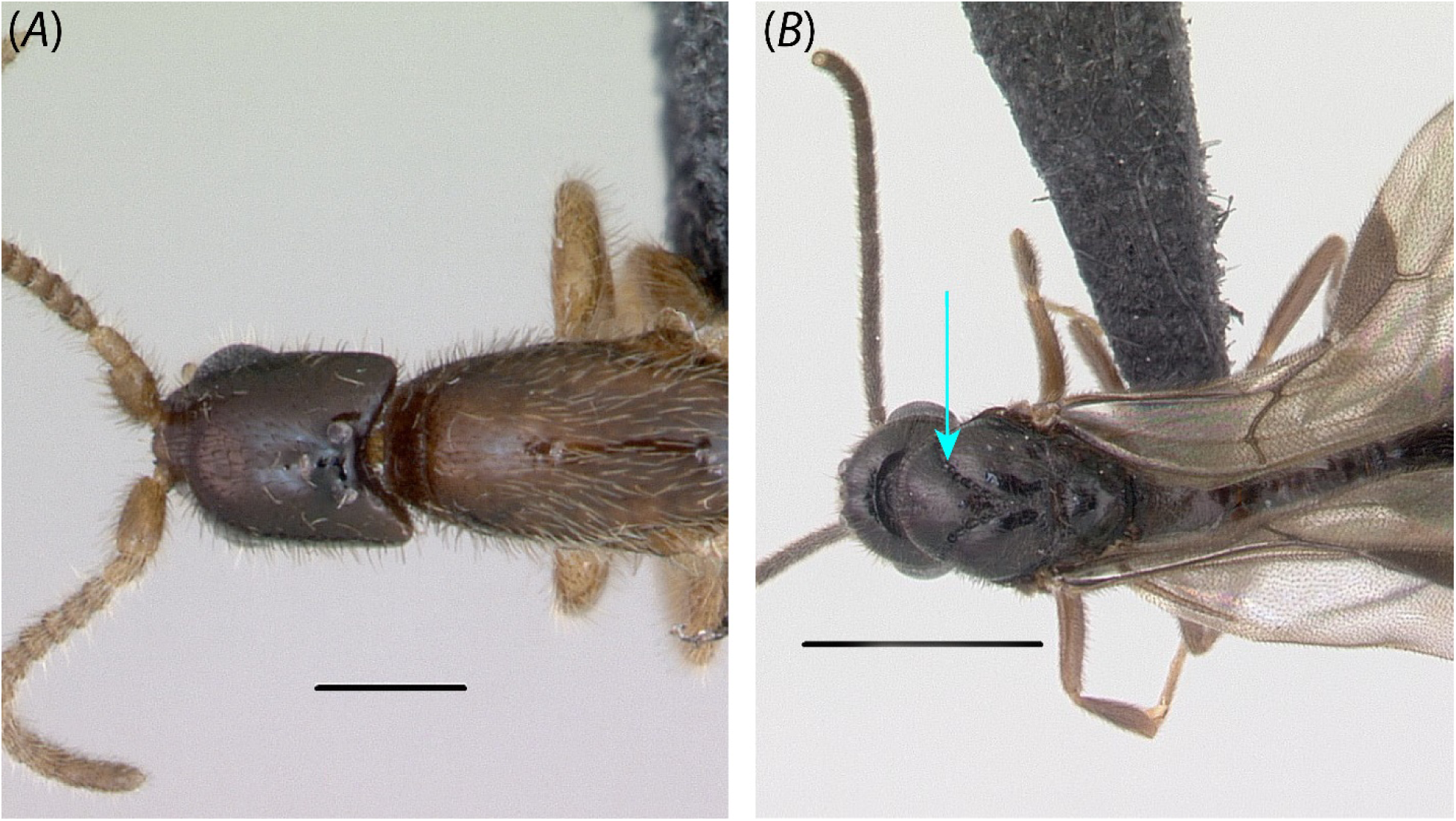
Dorsal view of (A) *Leptanilla* TH01 (CASENT0119776; April Nobile) and (B) *Protanilla* TH03 (CASENT0119791; Erin Prado). Scale bar A = 0.2 mm.; B = 1 mm.

**Fig. 29.**
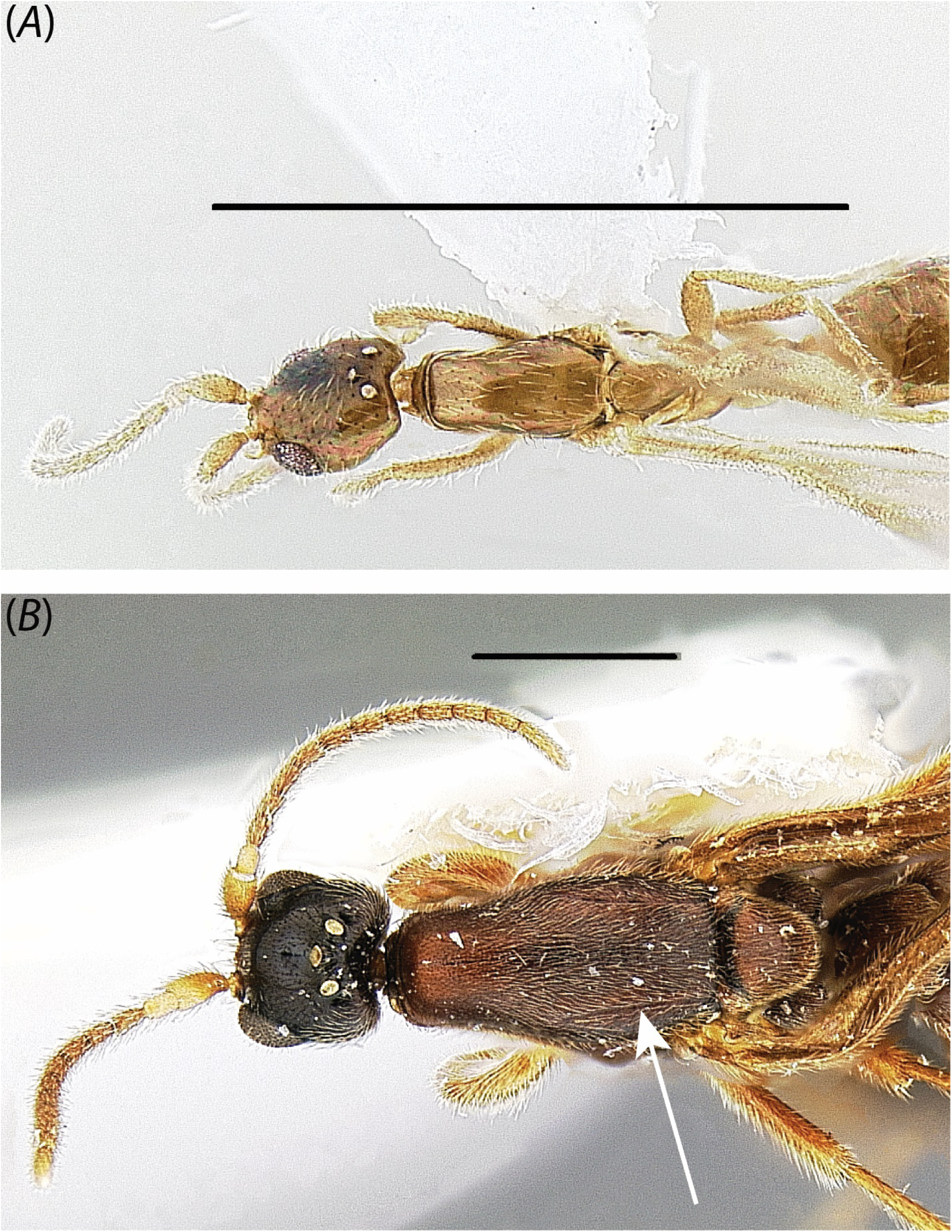
Dorsal view of (A) *Leptanilla* zhg-au01 (CASENT0758873; not sequenced in this study) and (B) *Leptanilla* zhg-my04 (CASENT0842558). Scale bar A = 1 mm.; B = 0.5 mm.

**Fig. 30.**
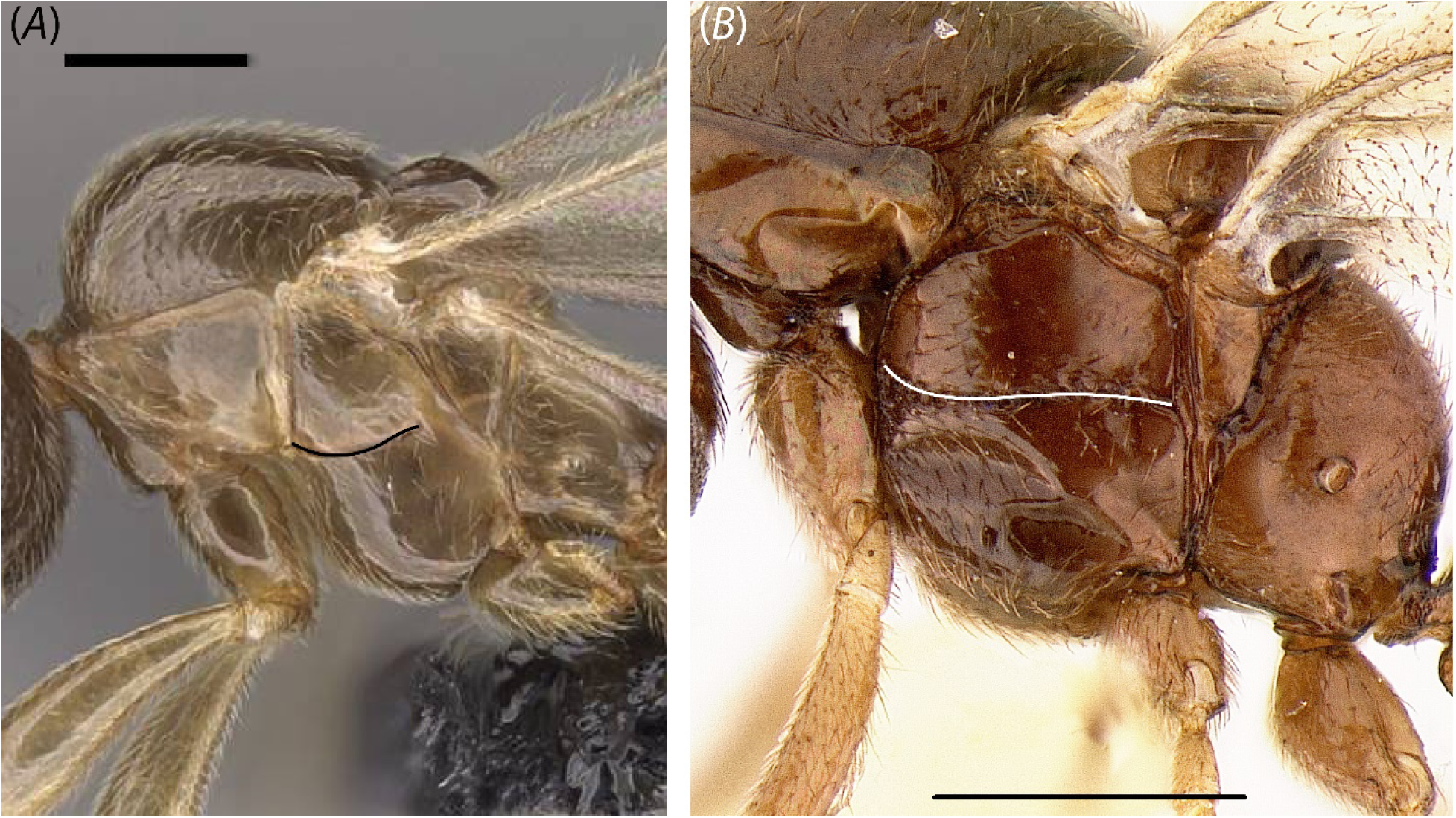
Profile view of mesosoma in (A) *Yavnella* TH02 (CASENT0119531; Michele Esposito) and (B) *Protanilla* zhg-vn01 (CASENT0842656). Scale bar A = 0.2 mm.; B = 0.3 mm.

**Fig. 31.**
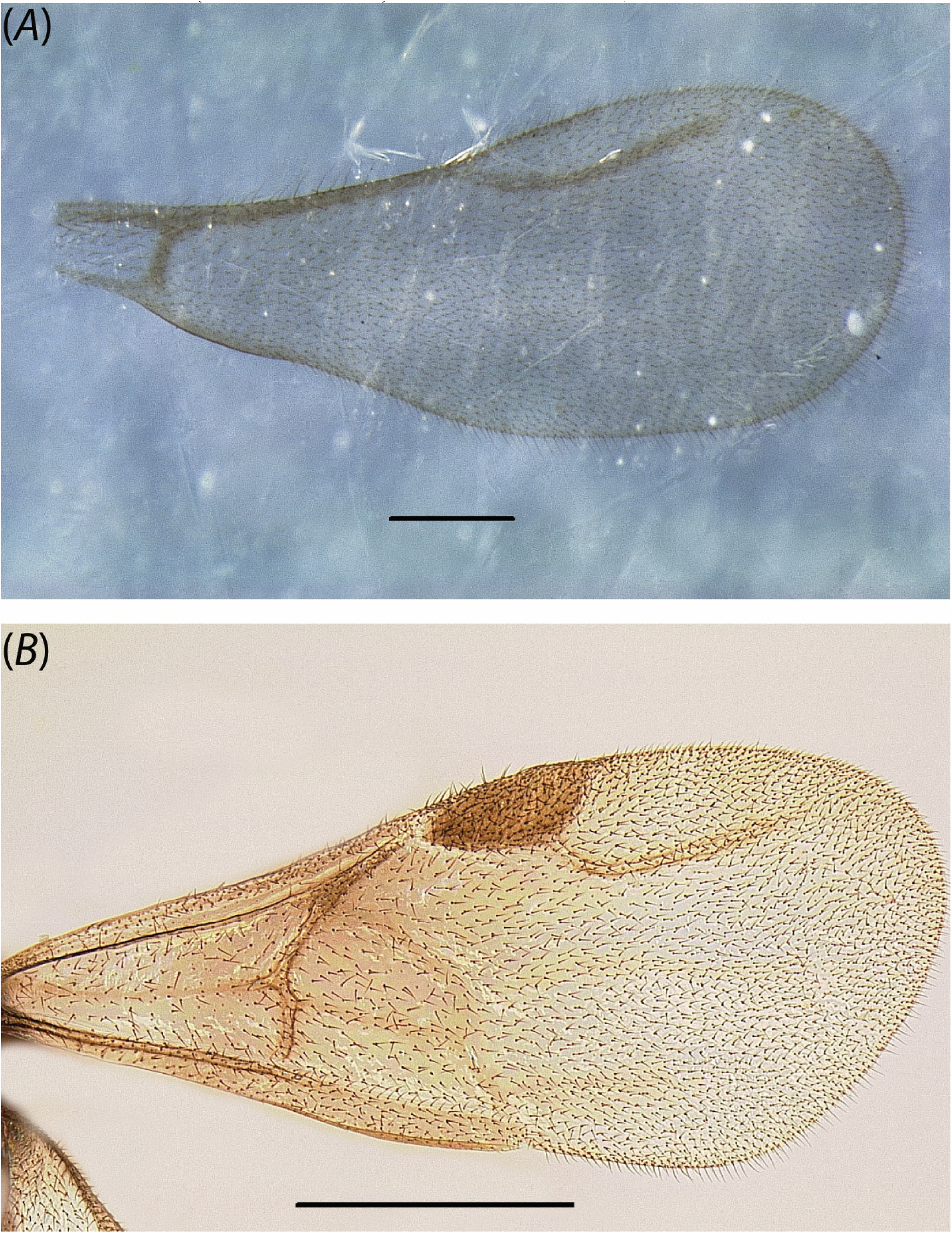
Forewing of (A) *Phaulomyrma javana* (MCZ:Ent:31142) and (B) *Protanilla* zhg-vn01 (CASENT0842613). Scale bar A = 0.2 mm.; B = 0.5 mm.

**Fig. 32.**
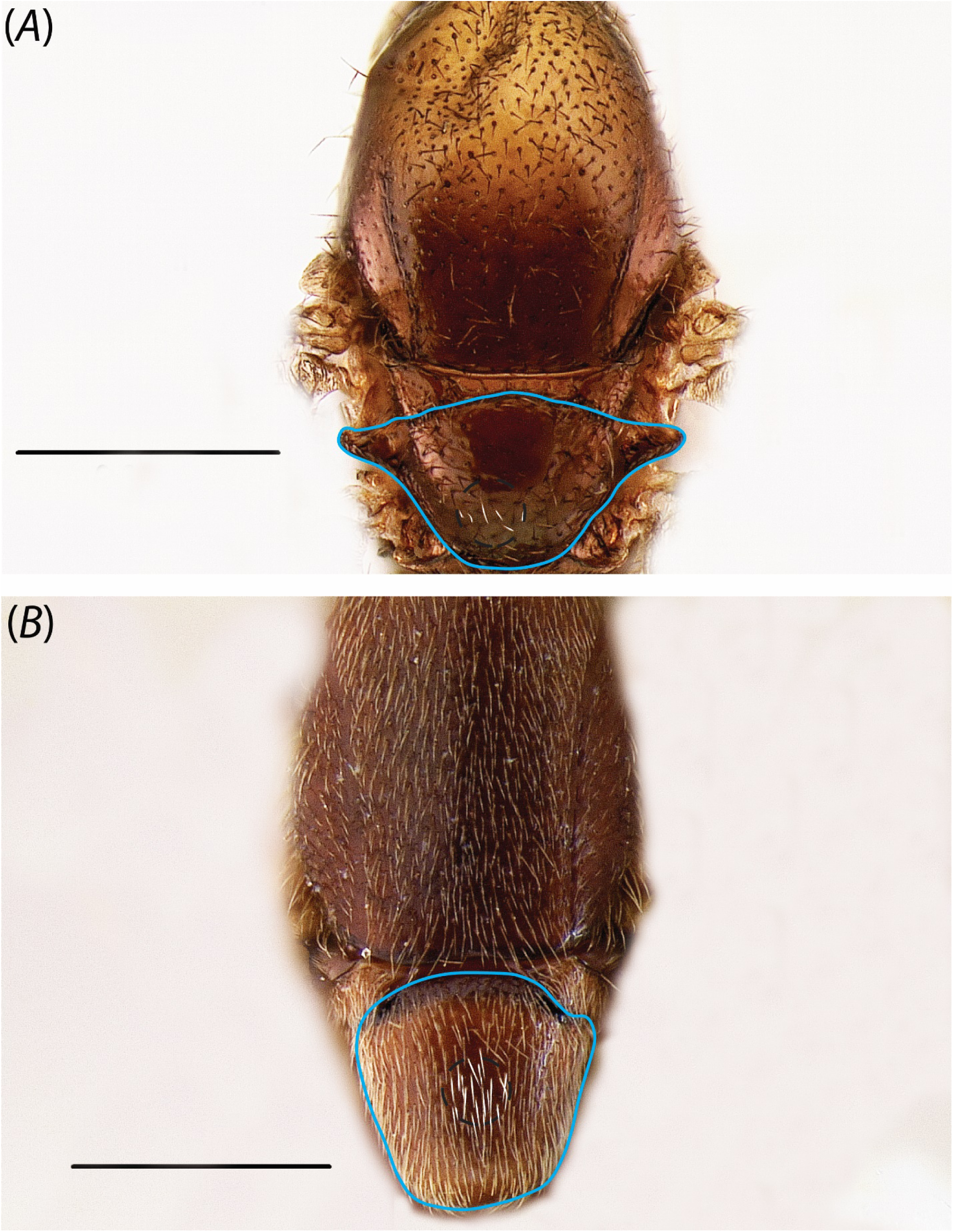
Dorsal view of mesosoma in (A) *Protanilla* zhg-vn01 (CASENT0842613) and (B) *Leptanilla* zhg-my04 (CASENT0842548). Scale bar = 0.3 mm.

**Fig. 33.**
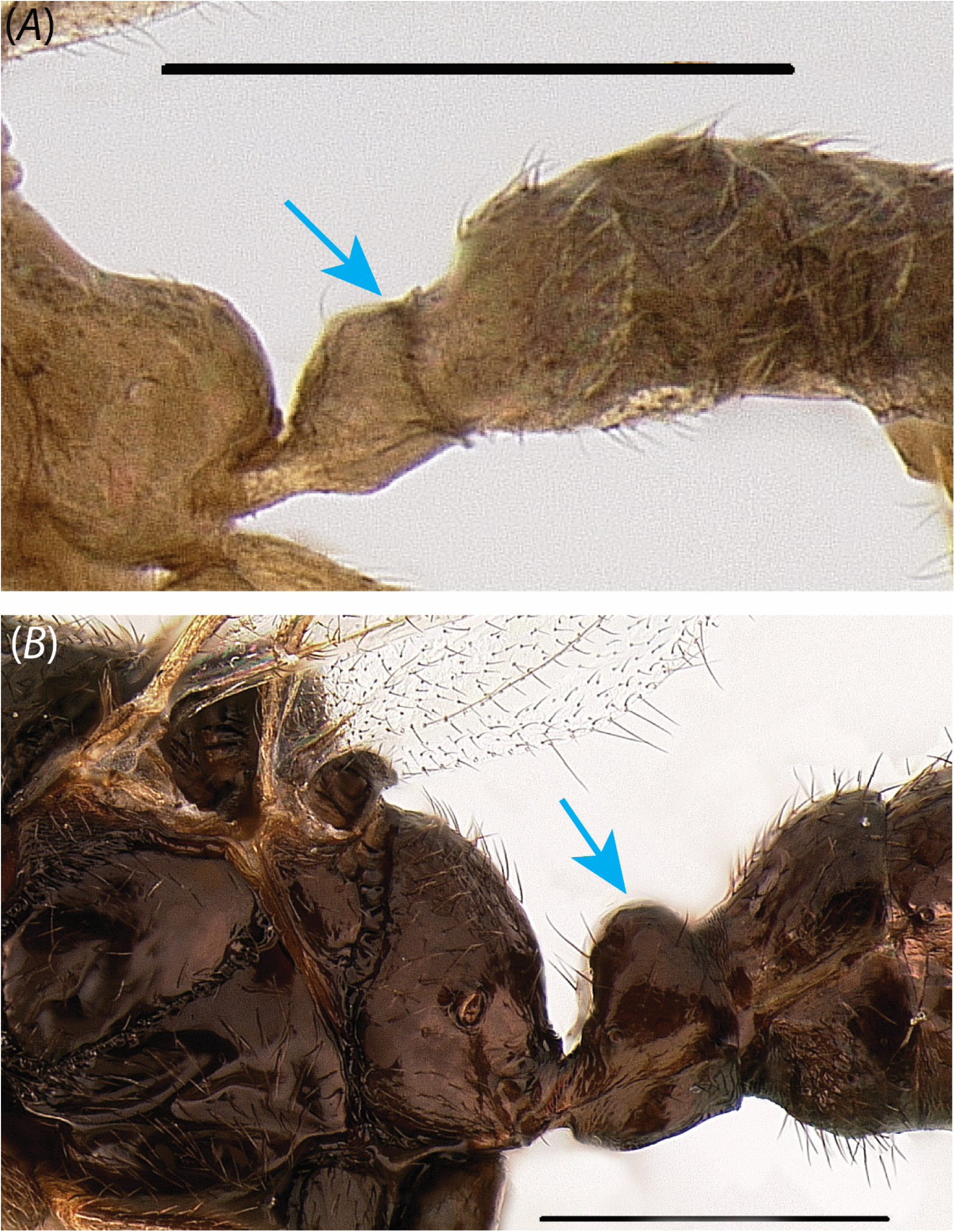
Profile view of petiole in (A) *Yavnella* zhg-bt01 (CASENT0106384) and (B) *Protanilla lini* (OKENT0011097; male described by Griebenow, in press) (not sequenced in this study). Scale bar A = 0.3 mm.; B = 0.5 mm.

**Fig. 34.**
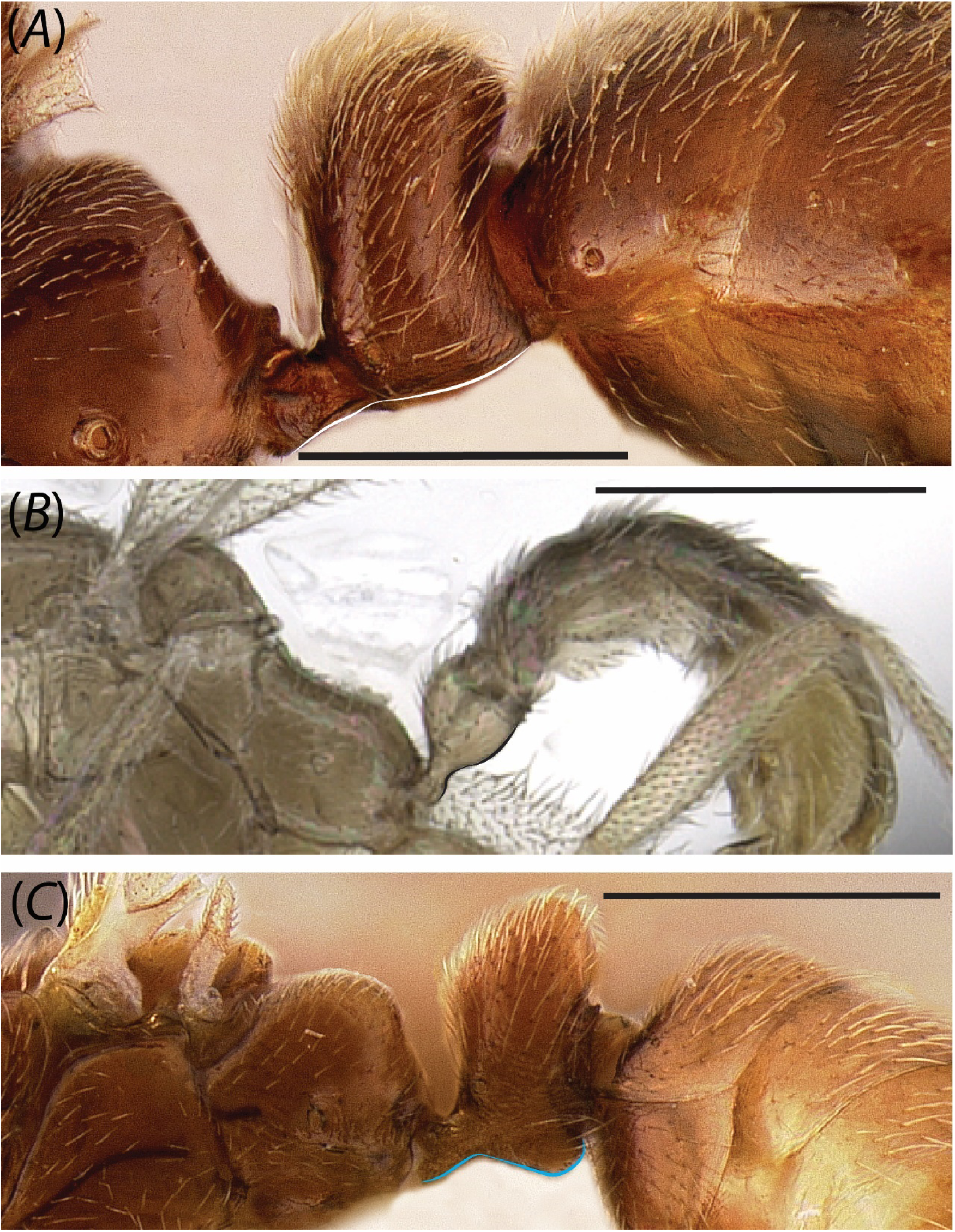
Profile view of petiole in (A) *Leptanilla* zhg-my04 (CASENT0842553), (B) *Yavnella* TH08 (CASENT0227555) and (C) *Leptanilla* zhg-my02 (CASENT0106417). Scale bars A, C = 0.5 mm.; scale bar B = 0.2 mm.

**Fig. 35.**
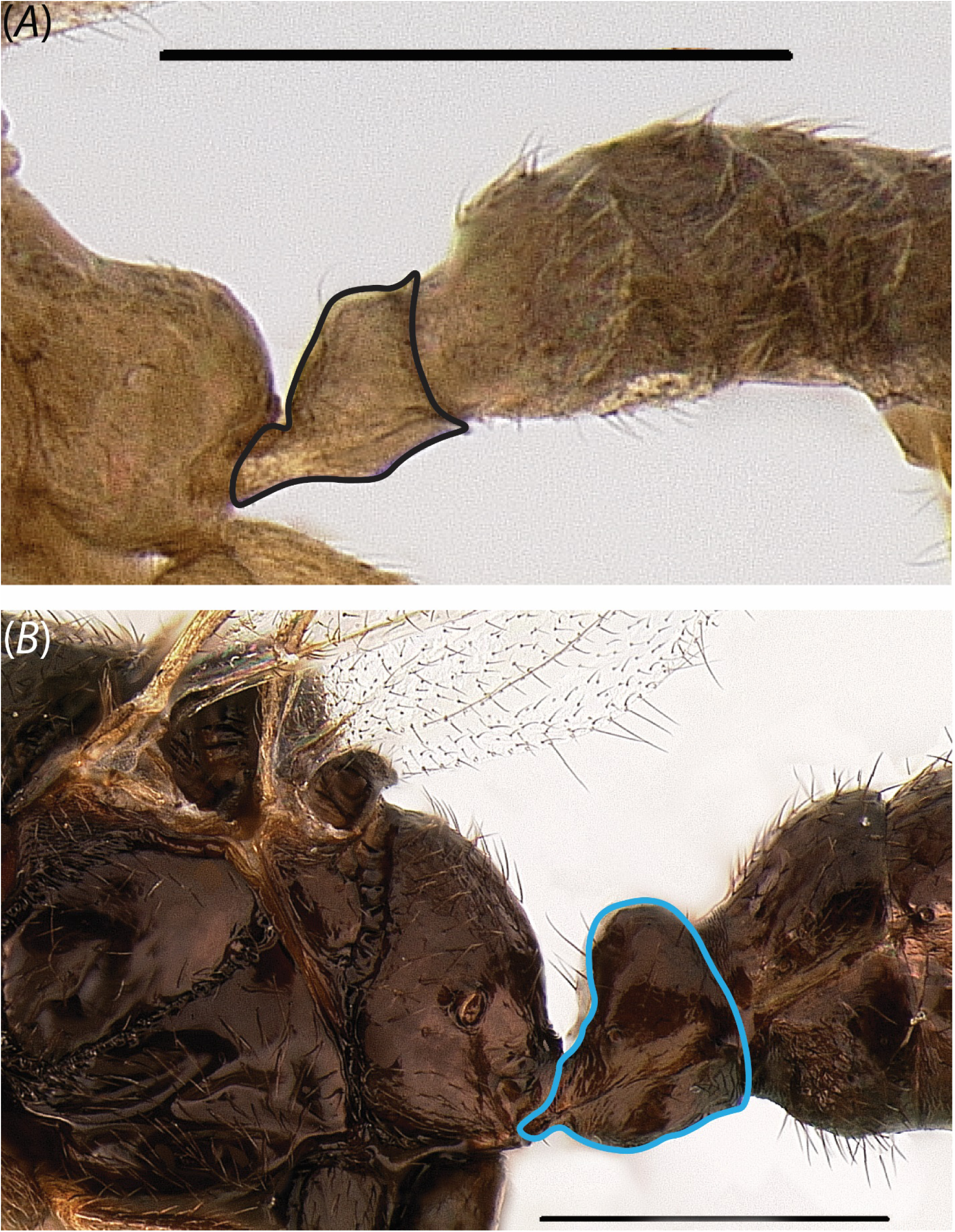
Profile view of petiole in (A) *Yavnella* zhg-bt01 (CASENT0106384) and (B) *Protanilla lini* (OKENT0011097; male described by Griebenow, in press) (not sequenced in this study). Scale bar A = 0.3 mm.; B = 0.4 mm.

**Fig. 36.**
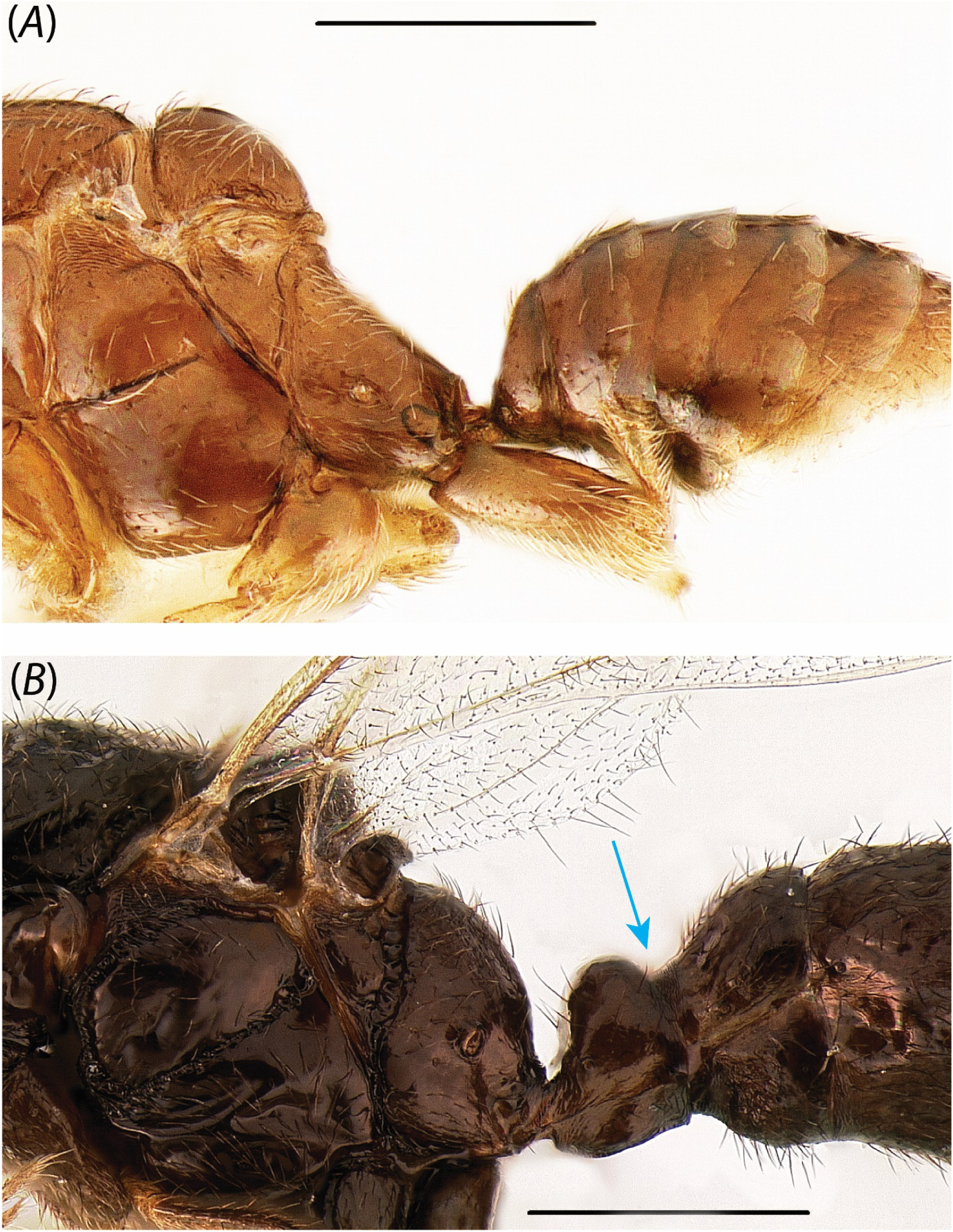
Profile view of petiole in (A) *Yavnella* zhg-th01 (CASENT0842621) and (B) *Protanilla lini* (OKENT0011097; male described by Griebenow, in press) (not sequenced in this study). Scale bar A = 0.3 mm.; B = 0.4 mm.

**Fig. 37.**
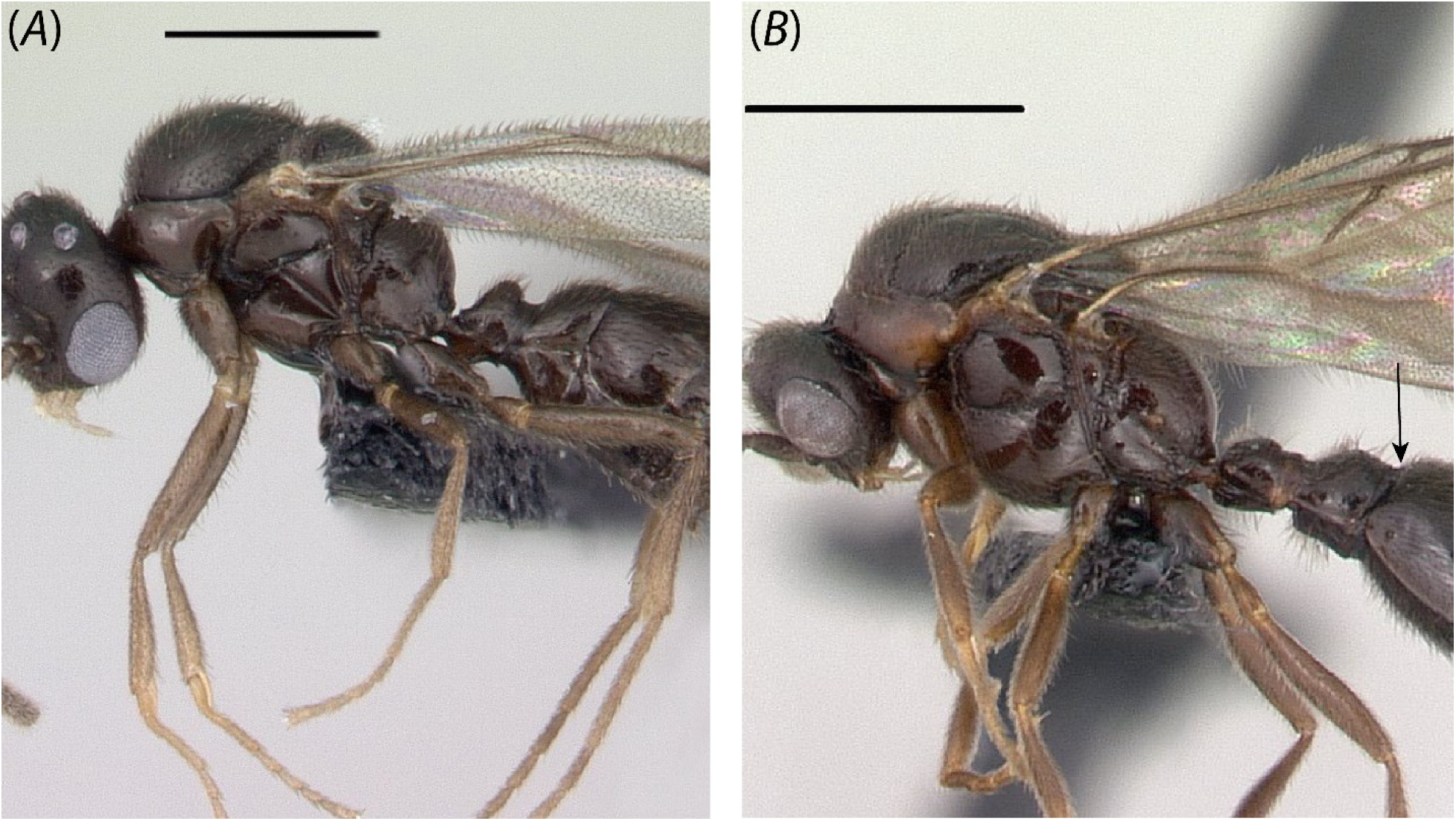
Profile view of (A) *Protanilla* TH02 (CASENT0128922) and (B) *Protanilla* TH03 (CASENT0119791). Scale bar A = 0.5 mm.; B = 1 mm.

**Fig. 38.**
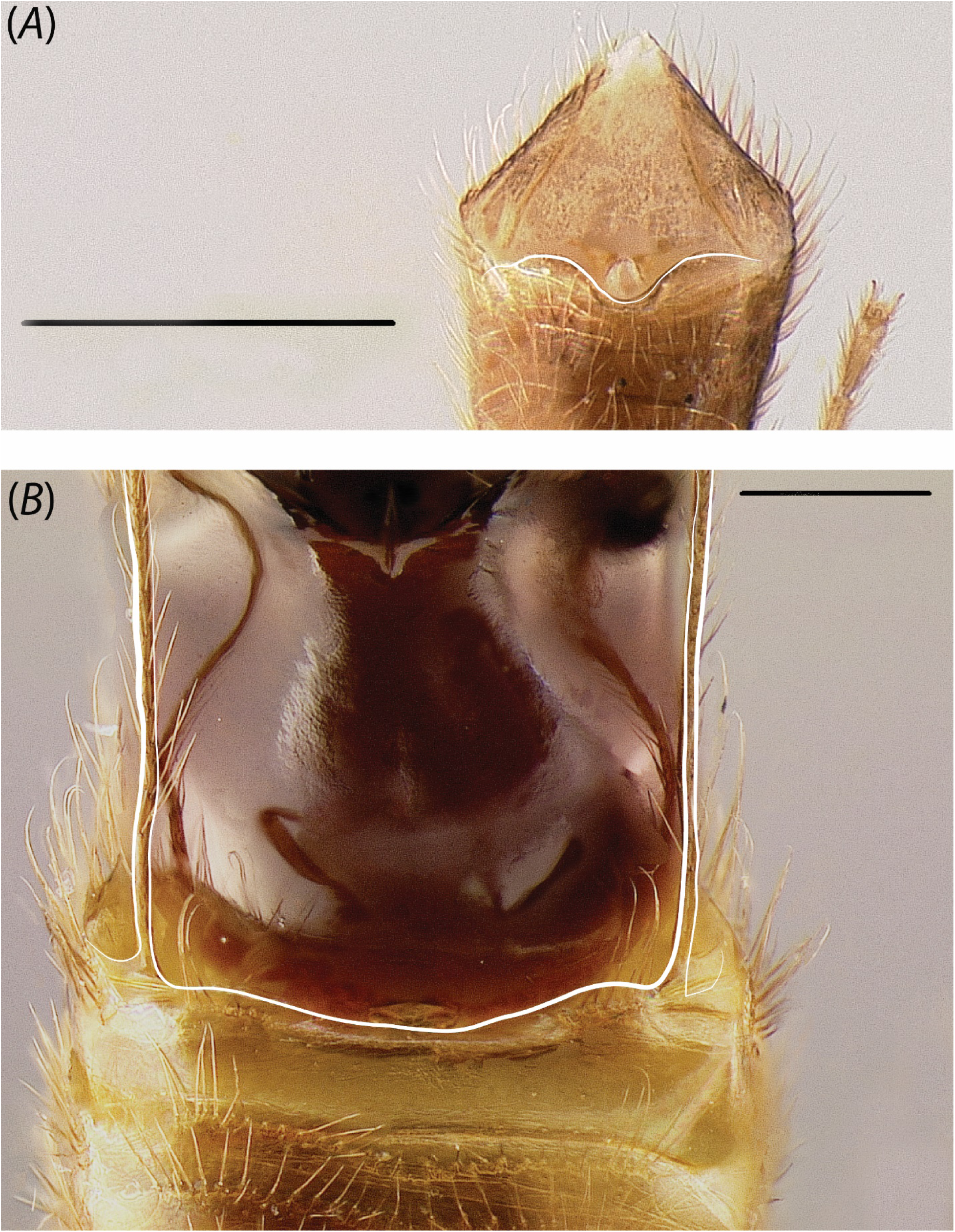
Ventral view of abdominal sternite IX in (A) *Leptanilla* zhg-th01 (CASENT0842619) and (B) *Leptanilla* zhg-my04 (CASENT0842553). Scale bar A = 0.3 mm.; B = 0.2 mm.

**Fig. 39.**
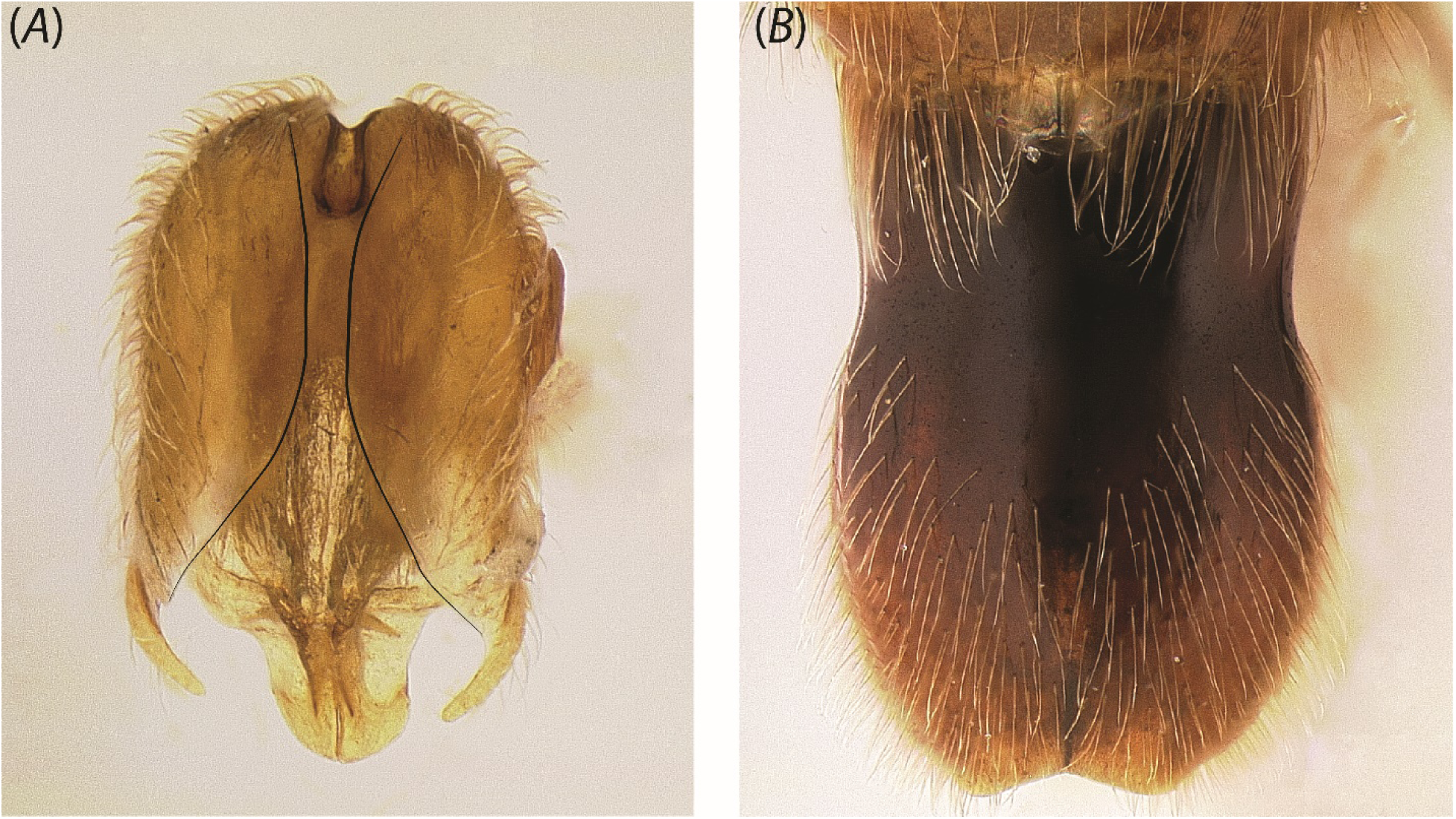
Dorsal view of genitalia in (A) *Yavnella* zhg-th01 (CASENT0842620) and (B) *Leptanilla* zhg-my04 (CASENT0842565). Scale bar A = 0.3 mm.; B = 0.4 mm.

**Fig. 40.**
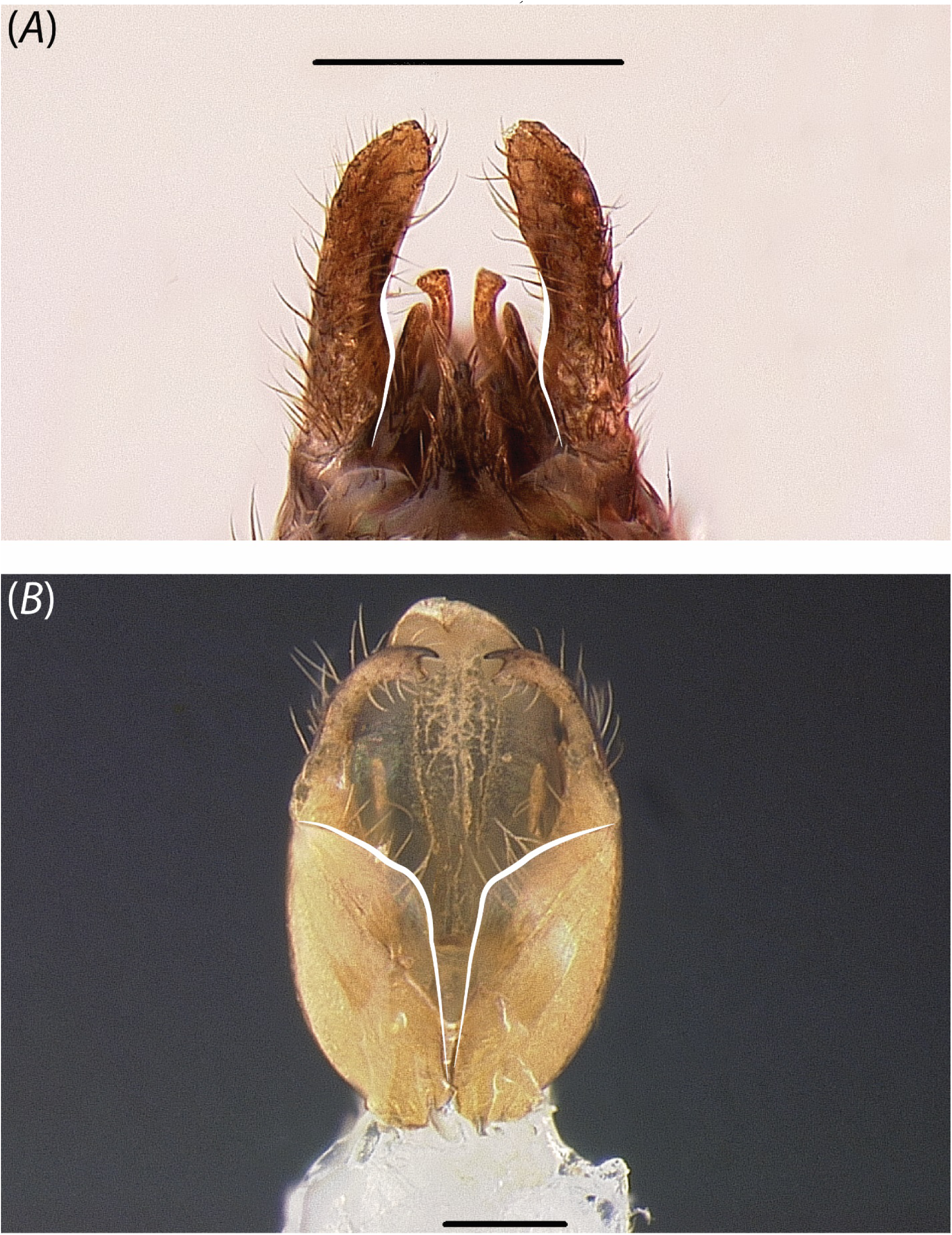
Ventral view of genitalia (A) *Protanilla lini* (OKENT0018456; male described by Griebenow, in press) (not sequenced in this study) and (B) *Leptanilla* ZA01 (CASENT0106354). Scale bar A = 0.3 mm.; B = 0.1 mm.

**Fig. 41.**
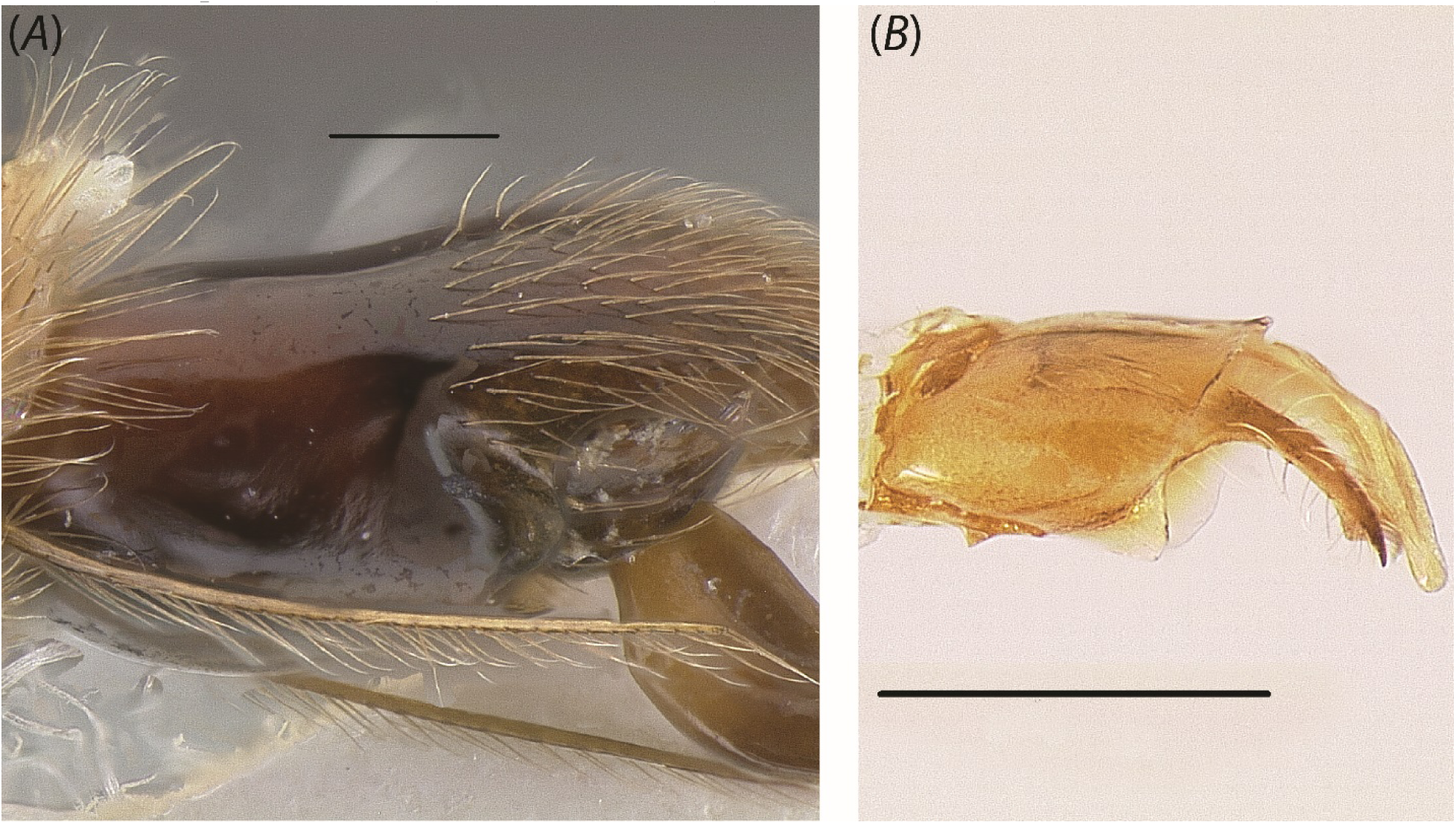
Profile view of genitalia in (A) *Leptanilla* zhg-my04 (CASENT0842558) and (B) *Leptanilla* ZA01 (CASENT0106354). Scale bar A = 0.2 mm.; B = 0.3 mm.

**Fig. 42.**
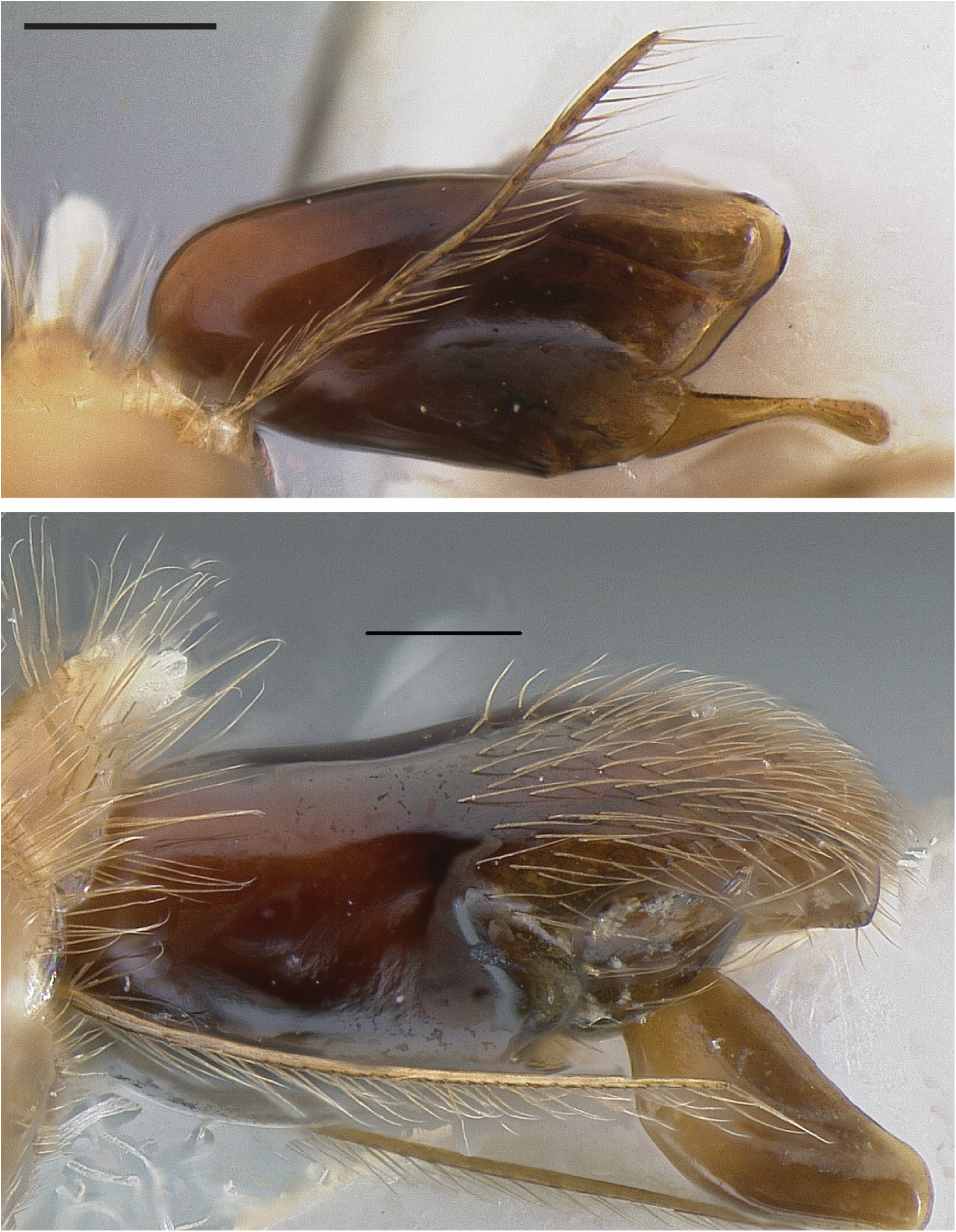
Profile view of genitalia in (A) *Leptanilla* zhg-my03 (CASENT0842545) and (B) *Leptanilla* zhg-my04 (CASENT0842558). Scale bar = 0.2 mm.

**Fig. 43.**
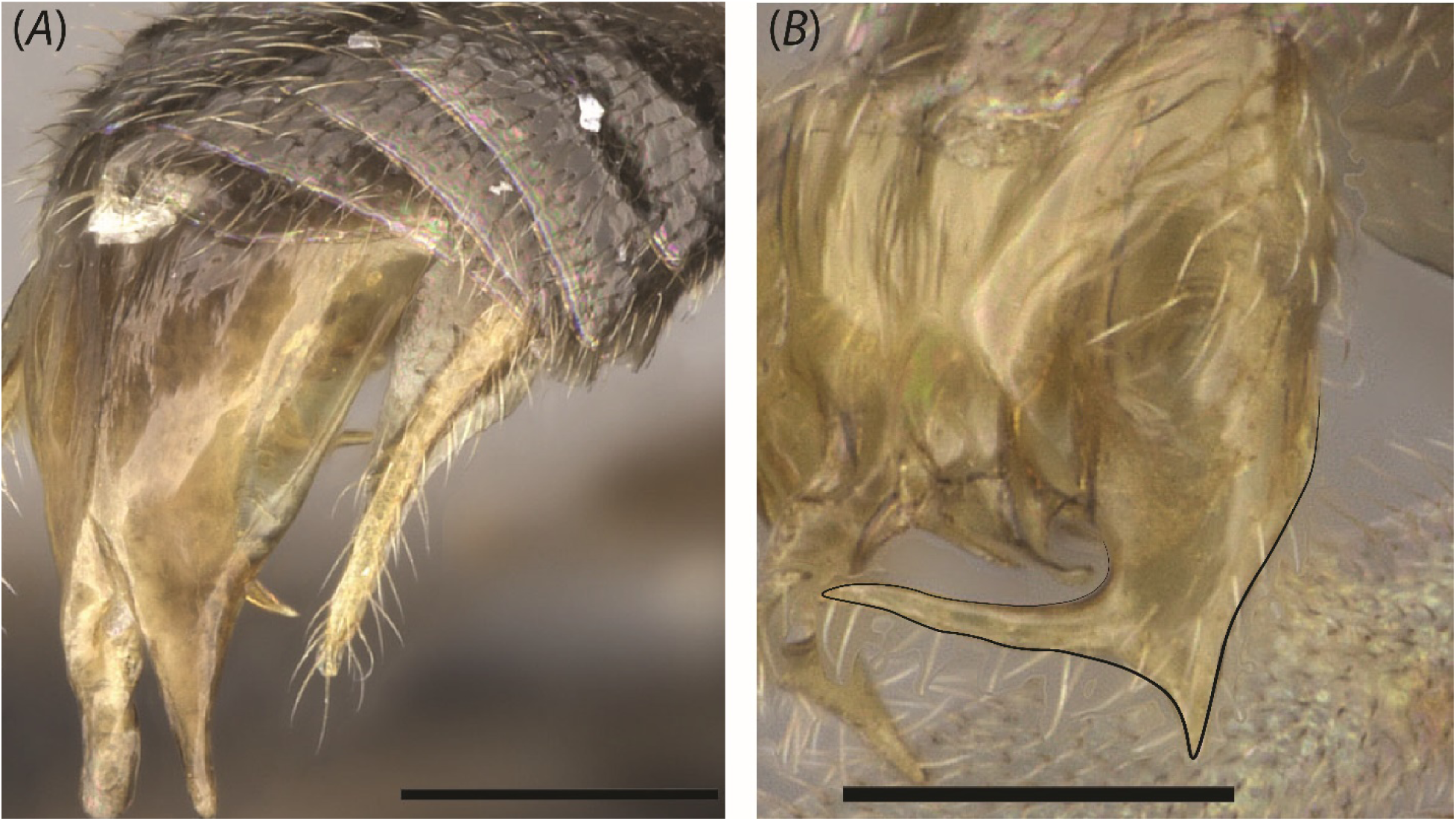
Posterolateral view of gonopodite in (A) *Yavnella argamani* (CASENT0235253) and (B) *Yavnella* TH08 (CASENT0227555) (both images by Shannon Hartman). Scale bar A = 0.2 mm.; B = 0.1 mm.

**Fig. 44.**
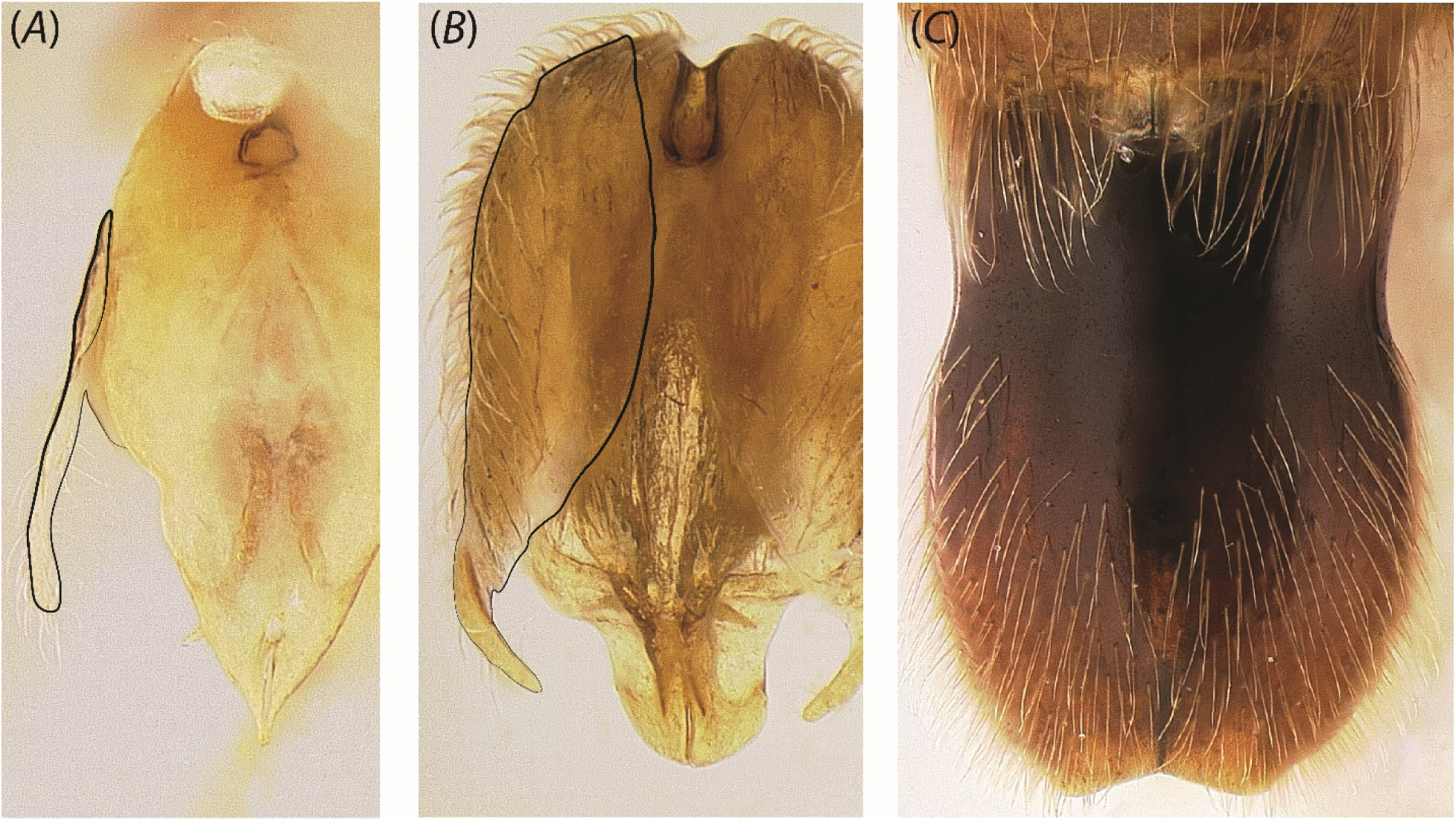
Posterior view of genitalia in (A) *Yavnella* cf. *indica* (CASENT0106378), (B) *Yavnella* zhg-th01 (CASENT0842620) and (C) *Leptanilla* zhg-my04 (CASENT0842565). Scale bar = 0.3 mm.

**Fig. 45.**
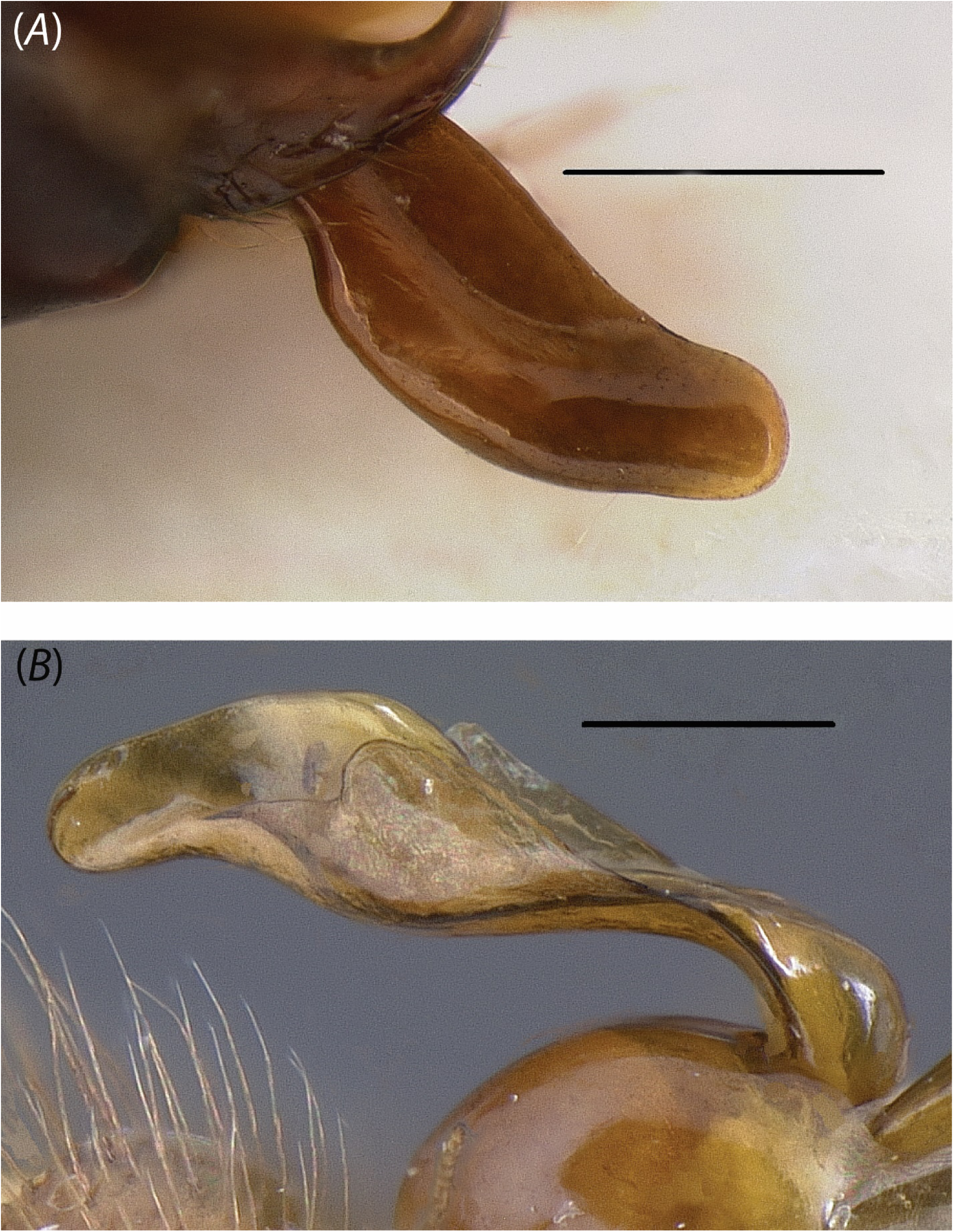
Profile view of penial sclerites in (A) *Leptanilla* zhg-my04 (CASENT0842550) and (B) *Leptanilla* zhg-my05 (CASENT0842571). Scale bar A = 0.3 mm.; B = 0.2 mm.

**Fig. 46.**
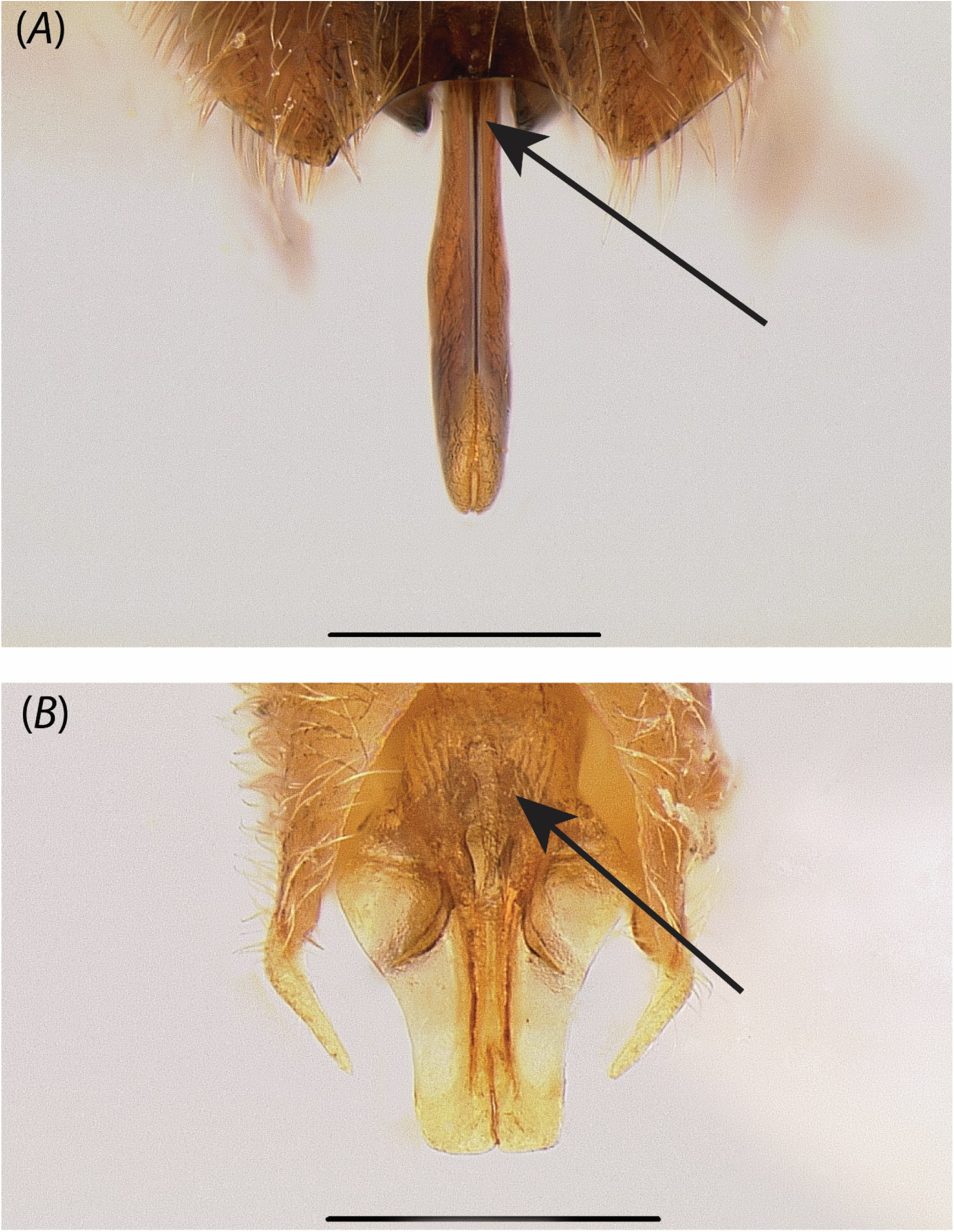
Posterior view of penial sclerites in (A) *Leptanilla* zhg-my04 (CASENT0842553) and (B) *Yavnella* zhg-th01 (CASENT0842620). Scale bar = 0.3 mm.

**Fig. 47.**
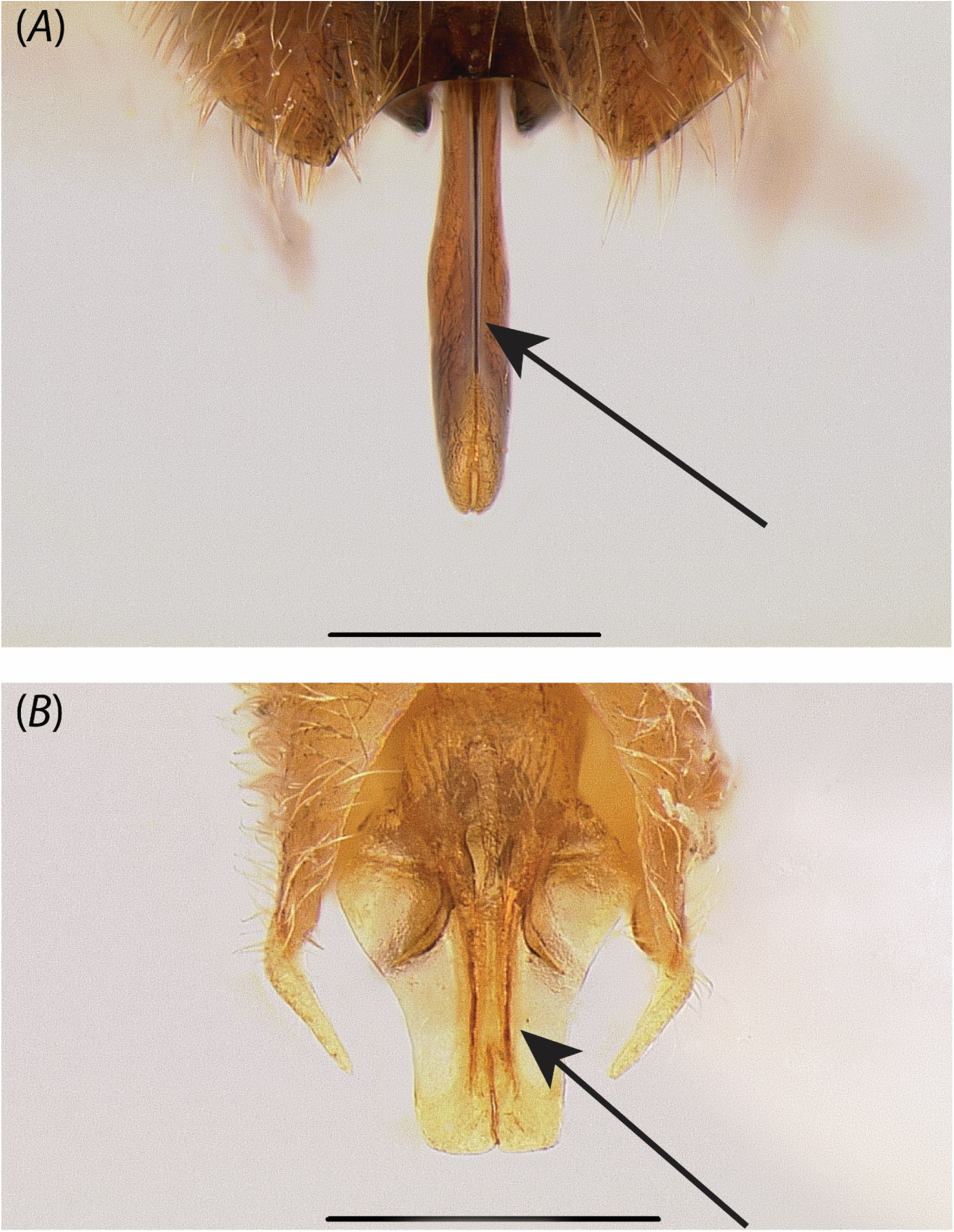
Posterior view of penial sclerites in (A) *Leptanilla* zhg-my04 (CASENT0842553) and (B) *Yavnella* zhg-th01 (CASENT0842620). Scale bar = 0.3 mm.

**Fig. 48.**
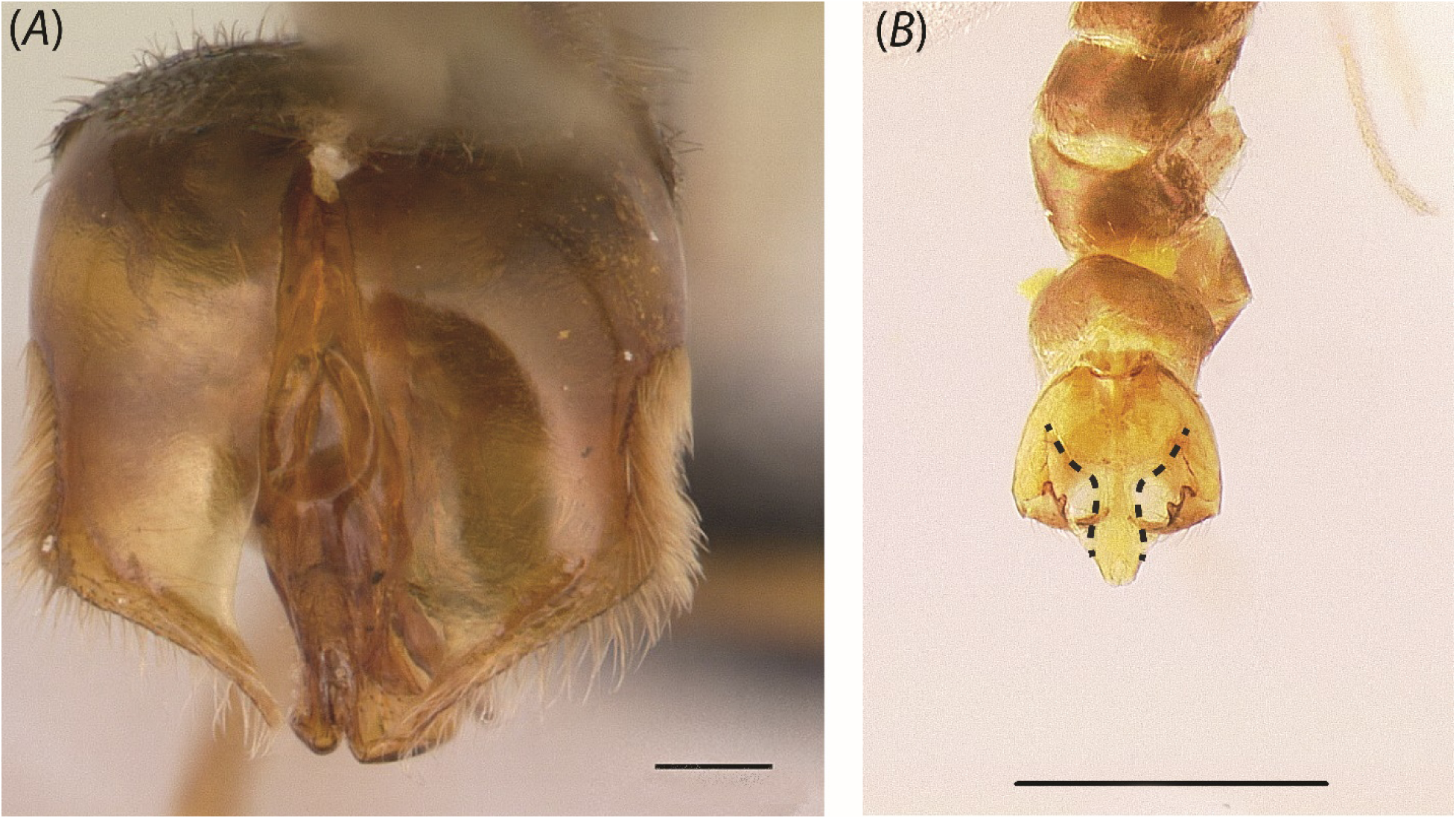
Posterior view of penial sclerites in (A) *Yavnella* TH06 (CASENT0129609; Erin Prado) and (B) *Leptanilla* GR02 (CASENT0106068). Scale bar A = 0.1 mm.; B = 0.3 mm.

**Fig. 49.**
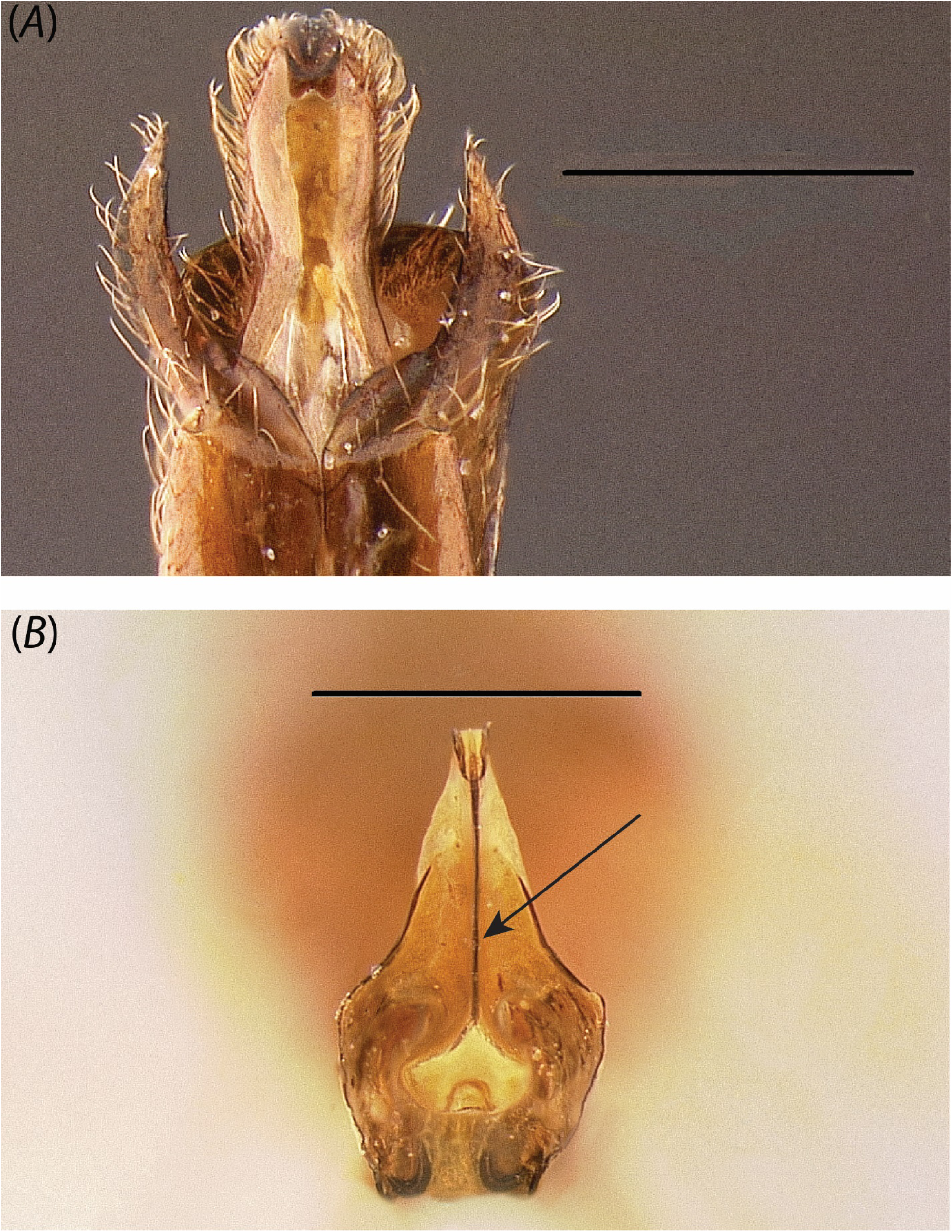
Posteroventral view of penial sclerites in (A) *Noonilla* zhg-my02 (CASENT0842595) and (B) *Leptanilla* zhg-my05 (CASENT0106432). Scale bar = 0.3 mm.

**Fig. 50.**
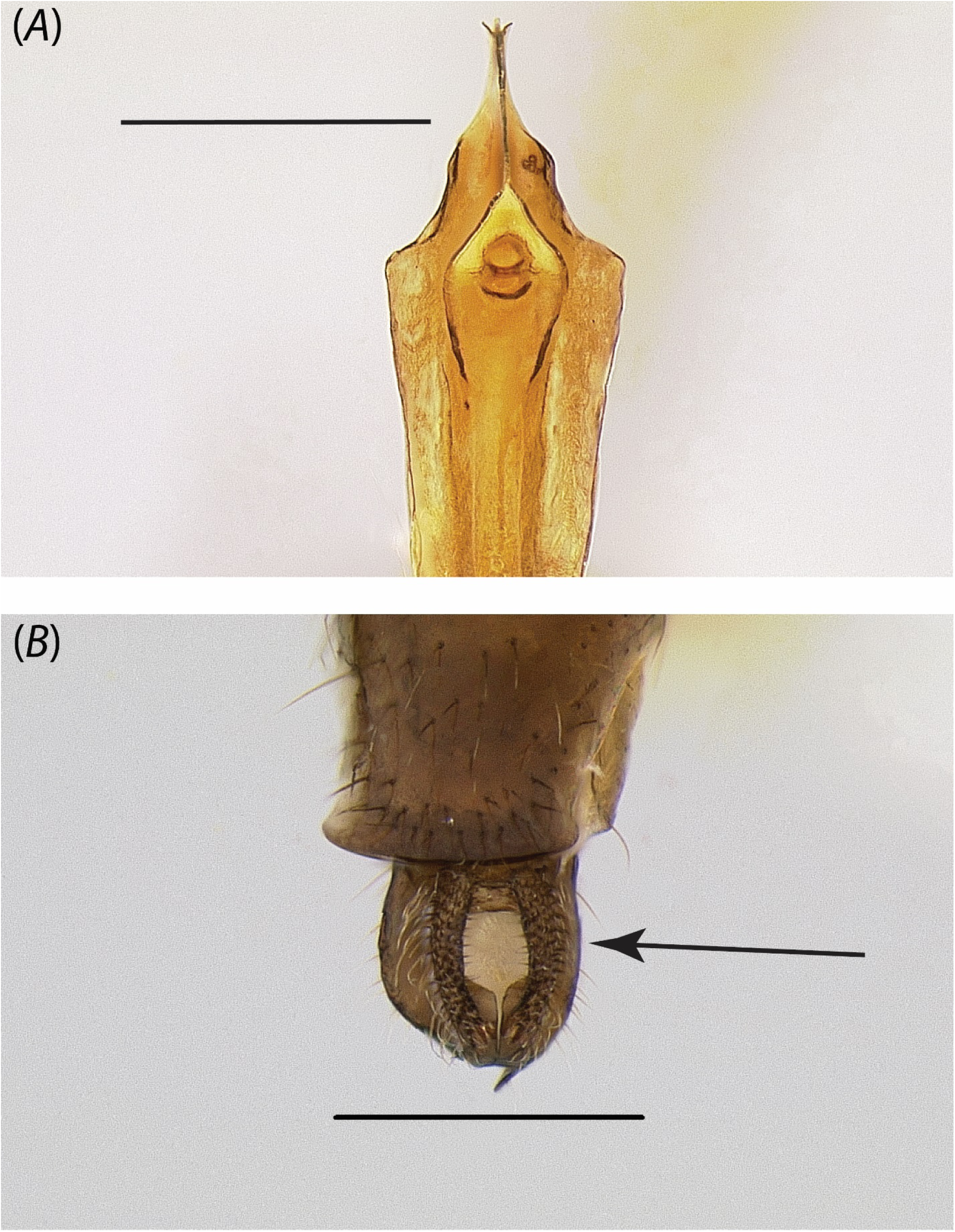
Posterior view of phallotreme in (A) *Leptanilla* zhg-my05 (CASENT0106432) and (B) *Noonilla* zhg-my02 (CASENT0842599). Scale bar A = 0.4 mm.; B = 0.3 mm.

**Fig. 51.**
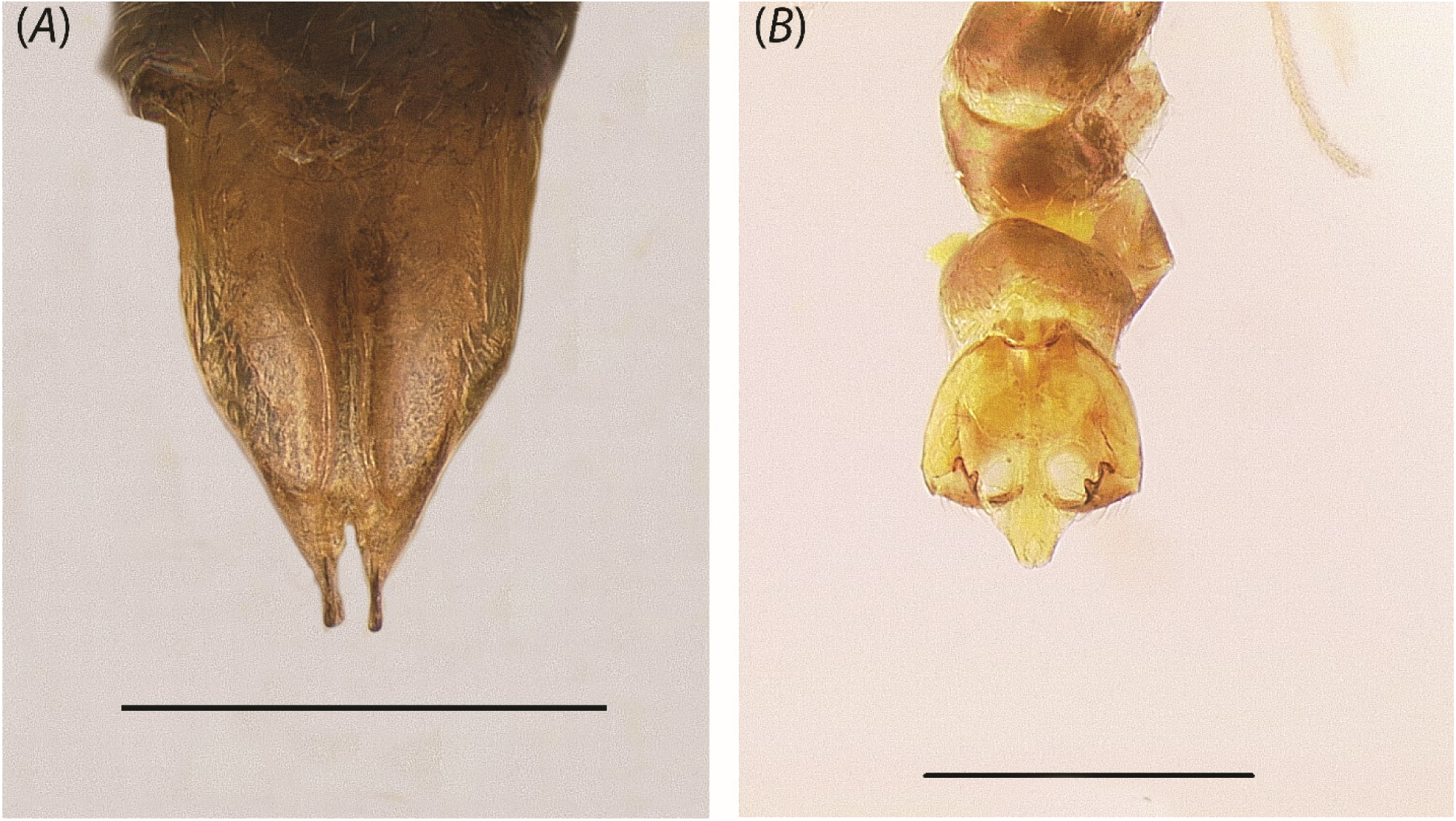
Posterodorsal view of the penial sclerites in (A) *Yavnella argamani* (CASENT0235253) and (B) *Leptanilla* GR02 (CASENT0106068). Scale bar = 0.3 mm.

